# Timing the onset of homologous recombination deficiency before breast cancer diagnosis

**DOI:** 10.1101/2025.10.17.683157

**Authors:** Michail Andreopoulos, Muchun Niu, Yang Zhang, Vinayak V. Viswanadham, Doga C. Gulhan, Hu Jin, Felipe Batalini, Gerburg Wulf, Chenghang Zong, Peter J. Park, Dominik Glodzik

**Affiliations:** Department of Biomedical Informatics, Harvard Medical School; Department of Molecular and Human Genetics, Baylor College of Medicine; Integrative Molecular and Biomedical Sciences Graduate Program, Baylor College of Medicine; Quantitative and Computational Biology Graduate Program, Baylor College of Medicine; Division of Medical Oncology, Mayo Clinic; Division of Hematology/Oncology, Beth Israel Deaconess Medical Center; Dan L. Duncan Comprehensive Cancer Center, Baylor College of Medicine; McNair Medical Institute, Baylor College of Medicine

## Abstract

Mutations in *BRCA1* and *BRCA2 genes*, whether inherited or somatically acquired, cause homologous recombination deficiency (HRD) in tumor cells. The timing of HRD onset in the emerging tumor lineage is unknown. Here, we present HRDTimer, an algorithm to infer the onset of HRD-driven mutagenesis prior to cancer diagnosis. We estimate that HRD arises at 34% of SBS1-based molecular time—corresponding to a median of 8.3 years (IQR 7.1–10.4) prior to diagnosis in triple-negative breast cancers, and 15.0 years (IQR 12.0–20.6) in ER-positive breast cancers. Bulk sequencing reveals accelerated SBS1 accumulation following neoplastic transformation compared to normal tissue, influencing the estimated age of HRD onset. Single-cell duplex sequencing confirms SBS1 acceleration in tumors and further shows that non-tumor cells largely lack the HRD signature, indicating that HRD is rare in pre-malignant cells, even in *BRCA1/2* mutation carriers. Together, our analysis pinpoints the onset of HRD before diagnosis, defining a window for detection and potential interception.

## Introduction

By age 80, *BRCA1* carriers face cumulative risks of 72% for breast cancer and 44% for ovarian cancer; the corresponding risks for *BRCA2* carriers are 69% and 17%.^1^ These risks are particularly elevated at younger ages, with age-specific hazard ratios up to 45-fold and 68-fold relative to the general population for breast and ovarian cancer, respectively.^2^ This elevated cancer risk underpins current clinical guidelines, which recommend risk-reducing surgeries such as prophylactic mastectomy and bilateral salpingo-oophorectomy.^3^ Because it remains uncertain which carriers will go on to develop cancer, many individuals undergo irreversible procedures that may not yet be necessary. This reliance on prophylactic surgery reflects a major clinical gap: the lack of reliable tools for risk stratification and early detection.

Earlier onset and elevated cancer risk suggest that neoplastic evolution progresses more rapidly in carriers of pathogenic *BRCA1/2* variants.^4^ This predisposition may result from an inherited inactivating mutation in one allele of a tumor suppressor gene, which makes bi-allelic loss more likely, and/or from elevated somatic mutagenesis. *BRCA1* and *BRCA2* are essential components of the homologous recombination (HR) DNA repair pathway; their loss forces reliance on error-prone mechanisms. The resulting HR deficiency (HRD) generates a characteristic mutational signature comprising structural variants, chromosomal copy number changes, insertions/deletions, and single-base substitutions.^5^ The full HRD phenotype^6^ and its associated signature are thought to arise only after bi-allelic inactivation,^5,7^ through a second somatic “hit” to the wild-type allele, after which HRD-driven mutagenesis can persist for years. Other studies, however, suggest that haploinsufficiency of *BRCA2* may allow features of HRD to emerge even before the second hit.^8,9^ Determining when HRD emerges during premalignant or tumor evolution is therefore a key unknown parameter in the evolutionary trajectory of HRD-associated cancers, with direct implications for precision-prevention strategies.

However, the timing of HRD emergence relative to clinical cancer diagnosis remains unknown in carriers of inherited *BRCA1/2* mutations and in patients with sporadic HRD tumors. Precursor lesions of HRD cancers are infrequently observed and poorly characterized, likely due to abrupt tumor growth and late diagnosis.^10,11^ Single-cell studies of premalignant tissue have found HRD-positive cells to be rare. These studies have largely relied on technologies that capture only gene expression^12^ or DNA copy number changes.^8,12^,13 As an alternative, cancer-archeology methods estimate the timing of key molecular events from tumor samples. Although existing algorithms can retrospectively infer the timing of mutations in cancer-implicated genes and whole-genome doubling,^14^ they cannot yet time the onset of a mutational process like HRD.

To pinpoint when HRD emerges before diagnosis, we developed the HRDTimer algorithm. Our analysis reveals that HRD often arises and remains active years before tumors become clinically detectable. Additionally, using a recent single-cell duplex sequencing technology, we show that HRD-driven mutagenesis is restricted primarily to tumor-lineage cells and, although exceptions do occur, is rarely detectable in non-tumor cells. These findings provide a framework for understanding the timing and cell-lineage specificity of HRD, and pinpoint when surveillance and preventive strategies might be most effective.

## Results

### Overview of the datasets

We analyzed five whole-genome sequencing (WGS) datasets: the PCAWG breast and ovarian cancer cohort,^15^ the SCANB triple-negative breast cancer cohort,^16^ the INFORM trial dataset comprising breast cancers from germline *BRCA1/2* carriers,^17^ the Genomics England (GEL) breast cancer cohort,^18,19^ and matched primary-relapse breast cancers.^21^ From these datasets, 142 cancers with both whole-genome doubling (WGD) and HRD (defined by HRDetect > 0.7)^5^ passed all quality-control criteria required for HRDTimer (Methods, “Sample quality control and curation before applying HRDTimer”) and were included in the timing analysis. Although the study focuses on breast cancer (115 of 142 tumors), we also included 27 ovarian cancers to assess generalizability to other WGD-positive, HRD tumor types. The breast cancer cohort comprised 32 tumors from germline *BRCA1/2* carriers, 15 with somatic mutations in HR-related genes, 21 with promoter hypermethylation, and 47 samples without known HRD-pathway alterations. We additionally included 70 breast cancer samples lacking both WGD and HRD for a comparative analysis of mutational burden and signatures. Additionally, to assess mutation rates during late tumor evolution, we included matched pairs of primary and relapsed breast tumors from 7 patients.^21^

To examine mutational burden and signatures in premalignant cells, we also analyzed wholegenome sequences from 32 single-cell-derived organoids of normal mammary epithelial cells obtained from 12 patients.^20^ Equivalent normal fallopian tube samples—the tissue of origin of highgrade serous ovarian cancer—were not available for this study.

Finally, to evaluate mutation burden and mutational signatures at single-cell resolution, and to investigate potential differential mutation rates between tumor and non-tumor lineages, we reanalyzed 21 breast cancers previously profiled using single-cell duplex sequencing (“scNanoSeq”).^22^ This approach enables the detection of cell-private somatic mutations with high accuracy, albeit at reduced sensitivity. To account for differences in mutation detection power due to sequencing coverage, ploidy, and mutation multiplicity, we developed a power-correction algorithm that integrates duplex coverage depth, DNA fragment counts, and per-cell copy number profiles (Supplementary Note 1). Six of the 21 scNanoSeq tumor samples were additionally characterized by deep WGS, enabling separation of clonal from cell-private mutations.

Together, these bulk and single-cell datasets span premalignant, primary, and relapsed states, as summarized in Figure S1.

### HRD-attributed mutations accumulate preferentially post-WGD

Inferring the timing of past events in both precancerous and cancer evolution requires a molecular clock, a mutational process that is active throughout life, accumulates at a steady rate, and is distinguishable from other mutational processes active in the tumor. Two single-base-substitution (SBS)^23^ signatures, SBS1 and SBS5, accumulate with age across diverse tissues and cancers^24^ and have been proposed as such clocks.^25^ SBS1 arises from spontaneous deamination of 5-methylcytosine at CpG dinucleotides, generating C>T substitutions. Importantly, the SBS1 mutational profile is largely distinct from other SBS signatures, including SBS3, the HRD-associated signature, enabling reliable attribution of mutations to each signature. Although SBS1 contributes a smaller fraction of the total mutational burden than SBS5, its context specificity enables accurate detection even at low mutation counts (Figure S2). In contrast, the relatively flat mutational profile of SBS5 makes it difficult to distinguish from SBS3, limiting its utility for HRD timing (Figure S3). In addition, SBS5 burden has been reported as elevated in smoking-associated tumors, indicating that its accumulation may be influenced by environmental exposures.^26^ We therefore adopted SBS1 as the molecular clock for HRD timing.

In the absence of longitudinal sampling, temporal reconstruction relies on genomic events that separate early from late mutations. Whole-genome doubling (WGD), which occurs in approximately half of HRD cancers,^27^ serves as one such anchor: mutations present at the time of WGD appear on two chromosome copies, while those acquired subsequently appear on only one. This logic holds only for samples with a single, clonal WGD event, so we restricted the cohort accordingly. To time individual somatic mutations relative to WGD, we applied the established Mutation-TimeR algorithm,^27^ which integrates somatic mutation allele fractions with allele-specific copy number profiles to classify mutations as early (pre-WGD) or late (post-WGD) according to their maximum a posteriori timing probability (Figure 1b).

**Figure 1.**
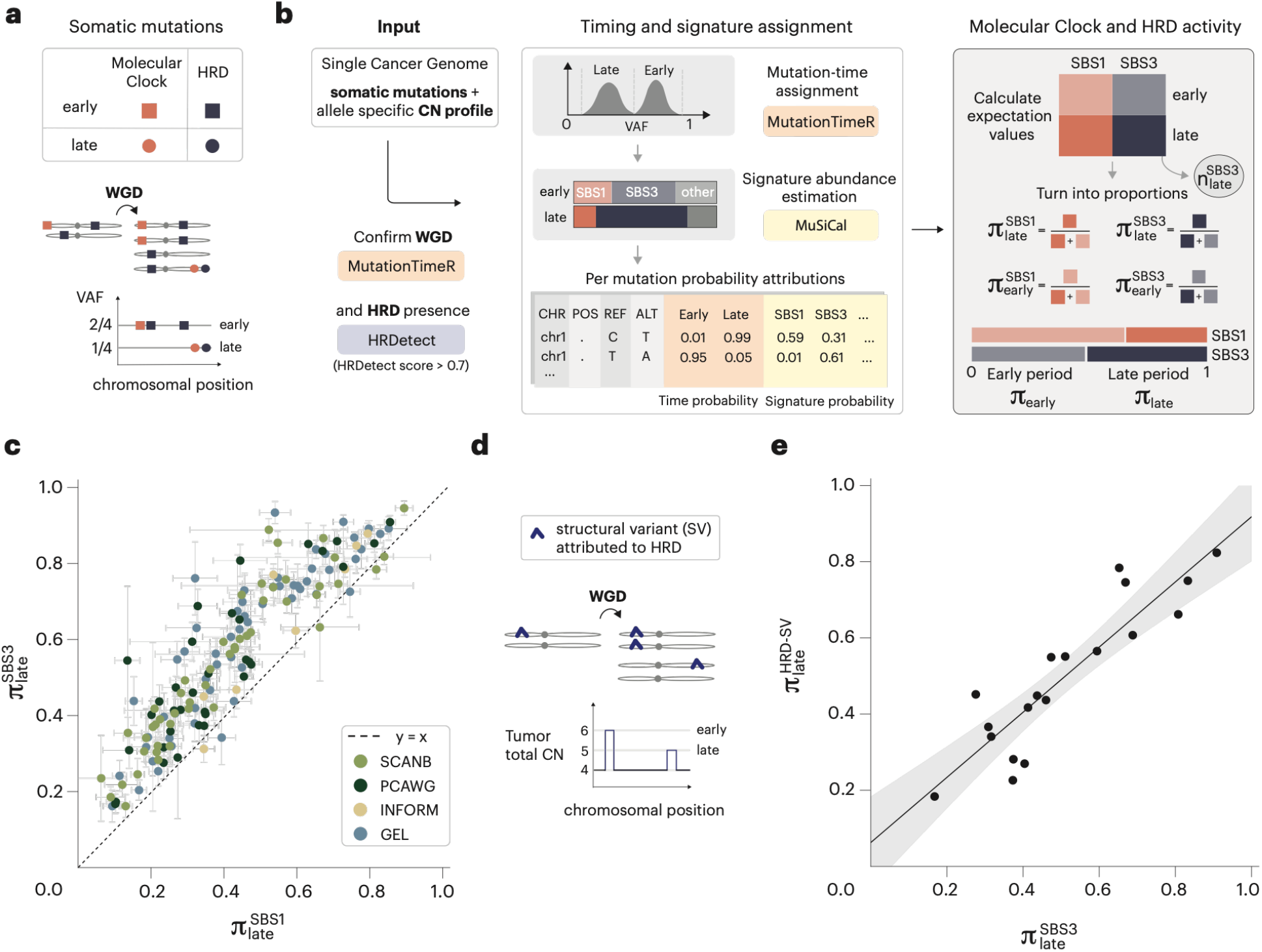
Shifts in the relative activity of the HRD signature over time. **(a)** Early and late mutations are distinguished based on variant allele fraction (VAF) and are attributed to homologous recombination deficiency (SBS3), the molecular clock (SBS1), or other mutational processes. Mutations that arose before whole-genome doubling (WGD) are present on two chromosomal copies, while those that occurred after appear on only one. **(b)** Schematic of mutation timing and signature assignment. Mutations are timed using VAF and allele-specific copy number profiles, followed by mutational signature assignment. Proportions of early and late mutations attributed to each mutational process (SBS1, SBS3) were computed. π_late_ denotes the proportion of late variants in regions with major copy number = 2 and minor copy number = 0 or 2, permitting retrospective timing relative to WGD. **(c)** Cohort-level comparison of the fraction of late mutations—defined by estimated mutation copy number of 1—attributed to SBS1 and SBS3. Error bars represent 95% confidence intervals from bootstrapping (see Methods). SBS3 shows a relative scarcity of mutations before WGD. See Figure S5 for interpretation. **(d)** Schematic illustrating how HRD-attributed structural variants (SVs), in PCAWG samples with both HRD and WGD, were classified as early or late based on their impact on copy number profiles. **(e)** Timing of HRD-attributed SVs relative to point mutations. HRD-associated SVs and point mutations show concordant timing.

We next inferred mutational signature exposures separately in the early and late mutation groups using MuSiCal,^28^ enabling estimation of the probability that individual mutations arose from specific mutational processes. This allowed us to examine the temporal distribution of HRD-associated mutations. SBS3 activity was detectable both before and after WGD, indicating that HRD-associated mutagenesis began prior to WGD (Figure S4); its distribution, however, was skewed toward the late period, with the proportion of late mutations among SBS3-attributed mutations 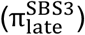 generally exceeding that among SBS1-attributed mutations (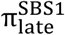; Figures 1c, S5). This suggests that although HRD mutagenesis begins before WGD, fewer SBS3 mutations accumulate during the pre-WGD period than would be expected under a model where SBS3 and SBS1 accumulate at constant, proportional rates throughout life.

To assess whether other HRD-associated variant types showed concordant timing, we timed structural variants (SVs), indels, and copy number losses. SVs—including tandem duplications and deletions with lengths <10 kb—were timed based on breakpoint-associated copy number changes (Figure 1, Figure S6; Methods).^29^ Indels attributed to ID6/ID8 were considered HRD-driven and timed using reported allele fractions. The proportion of late SBS3 mutations 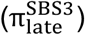 correlated strongly with that of late HRD SVs (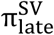, Pearson r = 0.89, P < 1 × 10^−6^; Figure 1d) and with late HRD indels (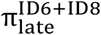, r = 0.91, P < 1 × 10^−12^; Figure S7). By contrast, the loss-of-heterozygosity (LOH) score—defined as the number of chromosomal segments affected by LOH^30^—was negatively correlated with 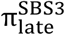 (r = −0.41, P = 0.02; Figure S7), as expected, since chromosomal losses occurring before WGD are more likely to produce persistent LOH detectable in the tumor sample. Together, these results indicate that HRD-attributed point mutations, indels, and structural alterations share a common temporal pattern, consistent with a single underlying mutational process.

### HRDTimer estimates HRD onset in units of SBS1-based molecular time

Having established that HRD onset preceded WGD, we next more precisely estimated when HRD began in molecular time between fertilization and cancer diagnosis. Our approach relied on the assumption that the SBS3:SBS1 relative mutation rate has remained steady since the onset of HRD. We validated this premise using post-WGD clonal, subclonal, and cell-private mutations in both bulk and single-cell data, confirming that SBS3 accumulates at a broadly constant rate relative to SBS1 within individual tumors throughout later stages of tumor evolution (Figure S8). We thus treated SBS3 as a second molecular clock that tracks HRD-associated mutagenesis from its emergence, which may occur post-zygotically. Because WGD occurred only once during the clonal period in each tumor in our cohort, it serves as a shared anchor point linking the SBS1 and SBS3 clocks, separating early (pre-WGD) from late (post-WGD) mutations for both processes.

HRDTimer classifies SBS1 and SBS3 mutations as pre- or post-WGD and uses this partitioning in three steps (Figures 2a, 2b, S9). First, it estimates the SBS1-based molecular time of WGD. Second, it calculates the sample-specific SBS3:SBS1 mutation rate from post-WGD mutations. Third, it applies this ratio to the number of pre-WGD SBS3 mutations to infer how many SBS1 mutations accumulated between HRD onset and WGD, thereby estimating the molecular time of HRD onset. HRDTimer reports both WGD and HRD times on a scale from 0 (fertilization) to 1 (the tumor’s most recent common ancestor, MRCA). These calculations rely exclusively on clonal mutations, which are reliably detected regardless of cancer cell fraction and greatly outnumber subclonal and cell-private mutations (Figure S8). The contribution of subclonal and cell-private mutations is incorporated later when translating molecular timing estimates into calendar time.

**Figure 2.**
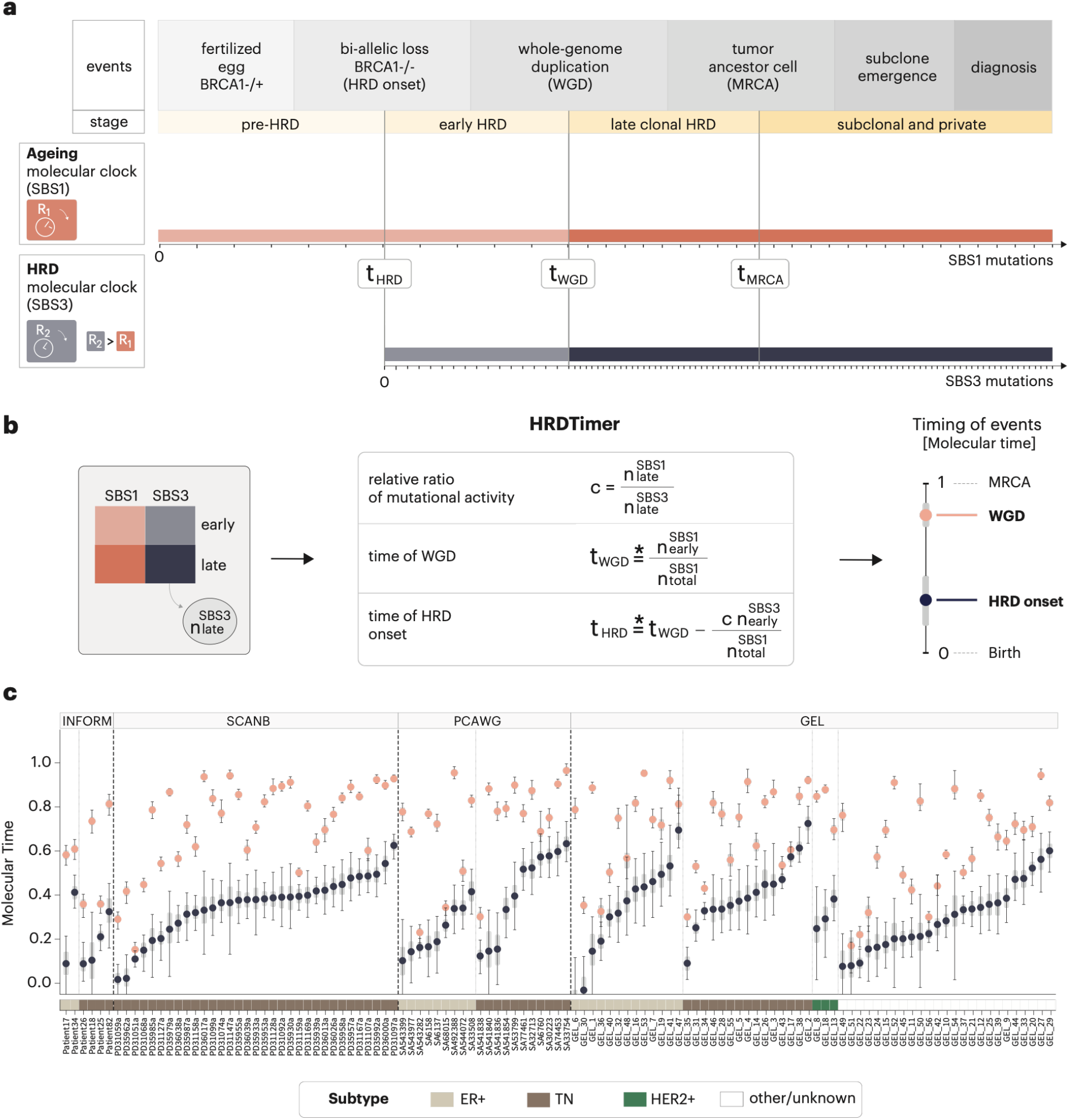
HRDTimer estimates the molecular (SBS1-based) timing of HRD onset. **(a)** Schematic time-line from fertilization to diagnosis, illustrating the unknown point of HRD onset (t_HRD_). Whole-genome duplication (WGD) serves as a temporal anchor separating the early and late clonal phases. The SBS1 mutation rate per Mb is assumed constant from fertilization, whereas the SBS3 rate is assumed stable from the onset of HRD. Tick marks on the x-axis illustrate the increase in mutation counts, with the rate changing at WGD if counts were not normalized by genome size; a correction for genome-size expansion due to WGD is applied throughout. **(b)** HRD onset is estimated from three components: the post-WGD SBS1-to-SBS3 mutation-rate ratio, the SBS1-based timing of WGD, and the number of early SBS3 mutations. Late mutations detected in bulk sequencing represent the tumor MRCA, but do not include subclonal and cell-private mutations. Asterisks denote genome-size–corrected quantities (see Methods for formulae). Confidence intervals are obtained by bootstrapping the early and late signature mutation catalogs (see Methods). **(c)** Estimated SBS1-based molecular timing of WGD (salmon) and HRD onset (navy) across four breast cancer cohorts (INFORM, SCANB, PCAWG, and GEL; n = 115 tumors). Samples are ordered within each cohort by the timing of HRD onset. Error bars denote 95% confidence intervals; gray bars indicate inter-quartile ranges from bootstrapping (see Methods and Figure S9). Colored bars below the x-axis indicate molecular subtype (ER+, TN, HER2+, or other/unknown). In all cohorts, HRD onset precedes WGD.

Across the 115 breast cancers that passed quality control (Methods), HRD arose after fertilization at a median time of 0.34 (IQR 0.20–0.44) of SBS1-based molecular time (Figures 2c, S10, S11). Median HRD onset times were similar in PCAWG (0.34), SCANB (0.38), and GEL (0.34), but substantially earlier in INFORM (0.16), a cohort composed exclusively of germline *BRCA1/2* carriers. Consistent with a predominantly post-zygotic origin of HRD, the 95% confidence interval for HRD onset excluded 0 (fertilization) in most tumors. Similar timing estimates were observed in the cohort consisting of 27 ovarian cancers, in which the median HRD onset was at 0.29 of SBS1-time (IQR 0.24–0.48; Figure S12).

Multiple analyses supported the robustness of these estimates. HRDTimer produced timing estimates for all samples, with uncertainty in both HRD and WGD timing quantified by bootstrapping (Figure S9). Estimates had the largest confidence intervals in tumors with very recent genome doubling, where the short post-WGD interval contains too few late clonal mutations to estimate the SBS3:SBS1 rate precisely; we therefore restricted the primary analysis to samples with sufficient numbers of early- and late-phase mutations and adequate purity and coverage (Methods, Figure S13). Furthermore, we evaluated our timing framework using additional sensitivity analyses and simulations. Although the accuracy of mutation timing by MutationTimeR has been validated,^27^ its performance also depends on the quality of the input purity and copy number estimates. Purity estimates derived from copy number analysis were consistent with the observed allele-fraction distributions of somatic mutations (Figure S14), and perturbing up to 10% of copy number segments had minimal impact on HRD timing estimates (Figure S15). Additionally, in simulations spanning a wide range of SBS3 rate regimes—including constant, gradual ramp-up, and oscillatory dynamics—our approach accurately recovered HRD onset across most scenarios, with accuracy reduced only when the SBS3 rate was unstable and either the pre- or post-WGD interval was short (Supplementary Note 3).

### Evidence for SBS1 acceleration from bulk WGS data

To translate SBS1-based molecular time into calendar time, we first evaluated whether SBS1 mutations accumulate at a constant rate or are instead influenced by proliferation and tumor evolution.^31^ We quantified clonal SBS1 mutations in WGD-positive breast cancers across four cohorts, capturing mutations accumulated up to the tumor’s most recent common ancestor (MRCA), and compared these with SBS1 burdens in normal breast epithelial cells. Mutation counts are reported per diploid genome throughout.

Tumor samples carried substantially higher SBS1 burdens than normal epithelial cells, with median burdens of 354 versus 93 SBS1 mutations per diploid genome. To estimate how much of the tumor burden could be explained by age-related accumulation, we fitted a linear regression of SBS1 burden against age in normal epithelial cells (1.11 SBS1 mutations per year) and applied it at the cohort’s median age. Tumor SBS1 burden was 3.8-fold above this age-matched expectation (Figure 3a). This excess is consistent with additional SBS1 accumulation prior to the MRCA, and is conservative because it excludes post-MRCA subclonal and cell-private mutations.

**Figure 3.**
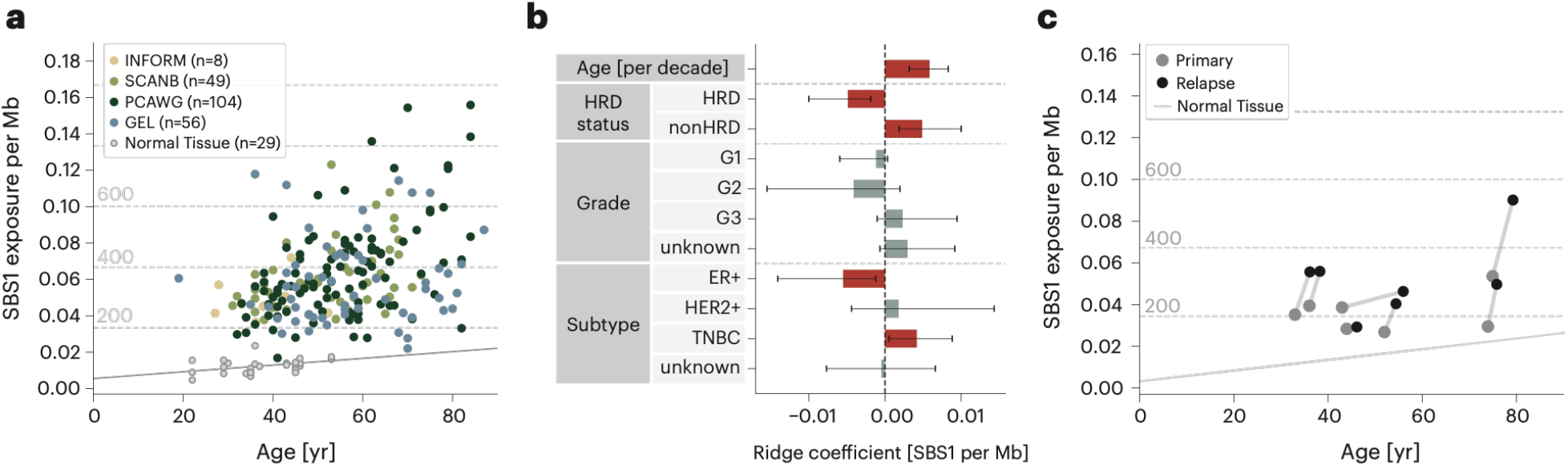
The SBS1 molecular clock scales with age in bulk WGS data. **(a)** Normalized SBS1 exposures in WGD-non-HRD (PCAWG, n = 104) and WGD-HRD breast cancers (INFORM, n = 8; SCANB, n = 49; GEL, n = 56) and in mammary epithelial organoids that were grown from single cells (Normal Tissue, n = 29), plotted against patient age. Tumors show elevated SBS1 burden relative to age-matched normal tissue. Horizontal dashed lines and gray numbers indicate SBS1 burden rescaled to a diploid genome. **(b)** Ridge-regression coefficients (bars) and 95% confidence intervals (whiskers) from a multivariate model of SBS1 burden across the tumor cohort in panel A (n = 217 tumors), with covariates grouped by age (per decade), HRD status, tumor grade and receptor subtype. Coefficients whose confidence intervals exclude zero are highlighted in red; non-significant coefficients are shown in gray. SBS1 burden is significantly associated with age, HRD status, and subtype. **(c)** Normalized SBS1 exposures in matched primary–relapse pairs from seven patients with relapsed breast cancer, compared with normal mammary epithelial organoids. Dark gray lines connect longitudinal samples from the same patient. Post-diagnosis SBS1 accumulation rates exceed those observed in normal tissue.

To assess whether variation in SBS1 burden among tumors is explained by patient and tumor-specific factors, we performed multivariable regression of SBS1 burden in tumors. Age was strongly associated with SBS1 accumulation, corresponding to an estimated increase of 37 SBS1 mutations per decade (P < 1 × 10^−5^). Additional independent contributions were observed from tumor grade (grade 3 vs grade 2: +80 SBS1 mutations, P = 0.045) and receptor status (TNBC vs ER+: +79 mutations, P < 0.001). Tumor-associated covariates therefore explain substantial variation in SBS1 burden, consistent with subtype-specific acceleration of SBS1 accumulation during neoplastic transformation prior to the MRCA. HRD tumors showed approximately 87 fewer SBS1 mutations than non-HRD tumors (P < 0.001). Effect sizes and P-values are derived from ordinary least squares regression; ridge regression, which accounts for collinearity among covariates, showed the same directions of effects (Figure 3b).

Finally, to evaluate SBS1 dynamics during late tumor evolution, we analyzed matched primary– relapse breast cancer pairs from seven patients.^21^ SBS1 burdens per diploid genome were quantified in each sample, and inter-sample accumulation rates were estimated as the difference in SBS1 counts divided by the time elapsed between the two samples (Figure 3c). The median SBS1 accumulation rate between primary MRCA and relapse was 39 mutations per year (range: 2.6– 64.9), approximately 35-fold higher than in normal cells. Excluding two low-mutation-rate outliers (PD11461 and PD13596) increased the median to 44.5 mutations per year, corresponding to a 40-fold higher rate. One of these outlier cases (PD13596), with relapse occurring 10 years after diagnosis following endocrine therapy and subsequent treatment interruption, illustrates that SBS1 accumulation reflects ongoing tumor growth dynamics during late evolution.

### Evidence for SBS1 acceleration from single-cell data

Bulk sequencing obscures cell-private mutations in tumor cells and non-clonal mutations in normal cells that have not undergone clonal expansion. To better calibrate the SBS1 molecular clock at single-cell resolution, we re-analyzed single-cell duplex sequencing (scNanoSeq) data from a recent study of 21 breast cancers.^22^

Single-cell analyses separate tumor from non-tumor lineages within a sample, enabling direct comparison of mutational burden. Although the single-cell study was designed to over-represent tumor cells,^22^ phylogenetic analysis using CellPhy^32^ identified 136 non-tumor cells and 585 cells in the tumor lineage across the 21 samples. After fitting mutational signatures for each cell and applying correction to account for sequencing depth and mutation detection sensitivity (Supplementary Note 1), we compared SBS1 burden between tumor and non-tumor lineages (Figures 4a, 4b). In seven samples with ≥5 non-tumor cells, tumor-lineage cells showed a median 1.8-fold higher SBS1 burden, even though both tumor and non-tumor cells were sampled from the same patients at the same time point (Figure 4c).

**Figure 4.**
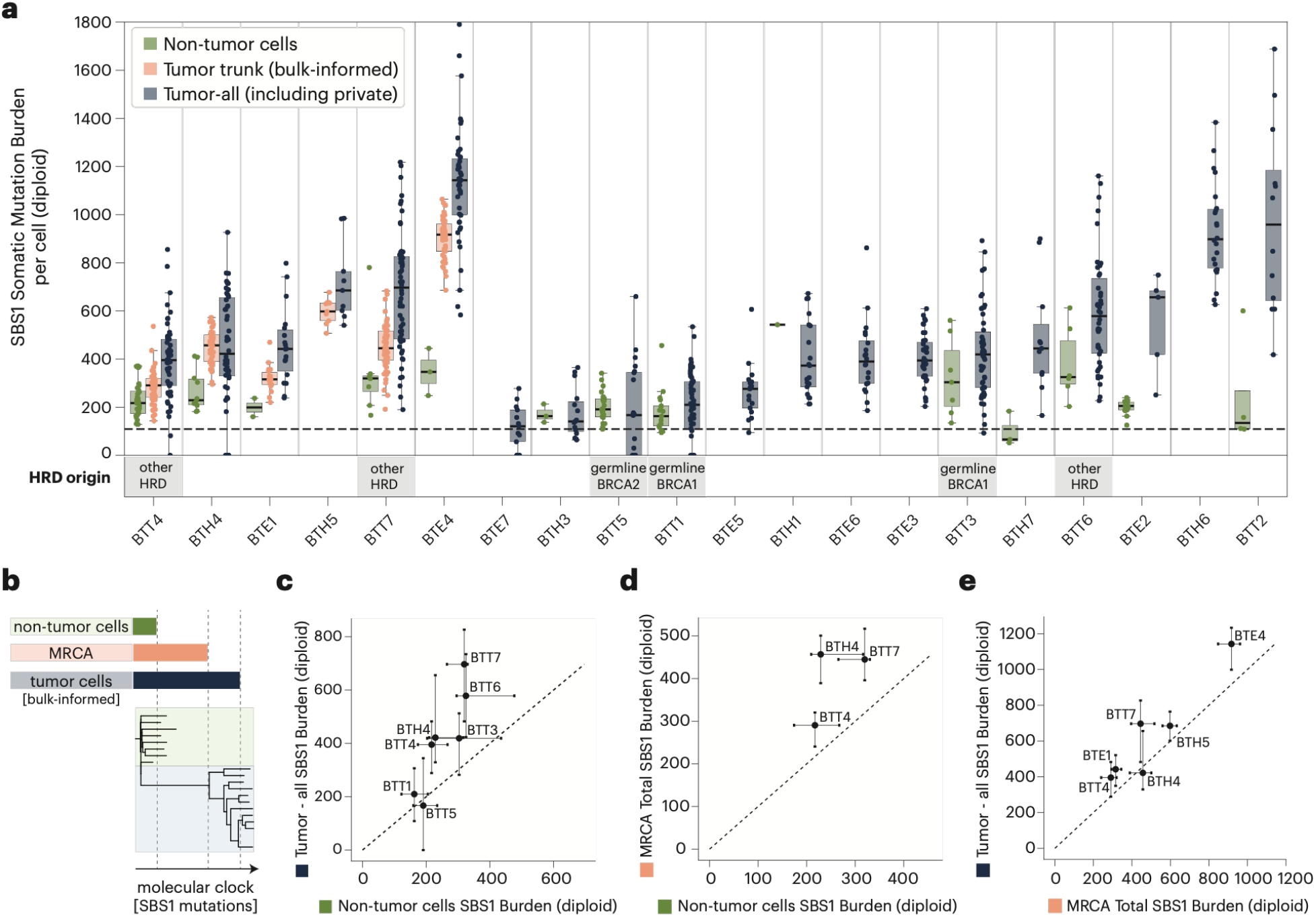
Single-cell scNanoSeq reveals SBS1 rate acceleration during tumor evolution. **(a)** SBS1 burden per cell across samples, stratified into non-tumor cells (green) and tumor cells, with mutations further subdivided as tumor truncal (bulk-informed, salmon), and all tumor mutations (navy). Counts are normalized to diploid genome size. Boxplots show median and IQR; points are individual cells. The dashed line marks the SBS1 burden expected in normal breast organoids (diploid-normalized) at the cohort’s median age at diagnosis. HRD origin is annotated below. **(b)** Schematic of how single-cell SBS1 burden captures non-tumor and post-MRCA accumulation, enabling reconstruction of SBS1 dynamics across tumor evolution. **(c)** Per-sample SBS1 burden, tumor versus non-tumor cells (samples with ≥5 cells per group). Points, per-sample medians; error bars, IQR; dashed line, y = x. **(d)** MRCA SBS1 burden (from matched bulk WGS) versus non-tumor single cells, for samples with ≥5 non-tumor cells and matched bulk data. Points above the diagonal indicate SBS1 acceleration before MRCA emergence. **(e)** Total versus truncal SBS1 burden per sample (cell medians), restricted to samples with matched bulk WGS. The extent of post-MRCA SBS1 accumulation differs across tumors, revealing rate heterogeneity that bulk sequencing cannot resolve.

Single-cell data further allow decomposition of SBS1 accumulation across evolutionary stages from premalignant cells to the tumor’s most recent common ancestor (MRCA) and up to diagnosis (Figure 4b). Truncal (MRCA) SBS1 burden was estimated using deep bulk WGS data available for six samples: mutations detected in both bulk and single cells were deemed truncal, and those unique to single cells were considered cell-private. To assess early acceleration in SBS1 accumulation, we compared MRCA SBS1 burden to that of non-tumor cells (Figure 4d). In three cases with both deep bulk data and ≥5 non-tumor cells, MRCA SBS1 burdens were higher than SBS1 burdens in non-tumor cells, with fold changes of 1.3, 1.4, and 2.0 (Figure 4e), consistent with accelerated mutagenesis during neoplastic transformation.

We next quantified post-MRCA SBS1 mutations, defined as mutations present in single cells but absent from bulk-inferred truncal profiles. In six samples where MRCA burden could be reliably estimated, tumor cells showed a median excess of 116 SBS1 mutations per diploid genome. Four samples showed larger excesses (105–252 mutations), while two showed similar tumor-cell and MRCA SBS1 burdens (Figure 4e). These estimates of cell-private SBS1 mutations were used in downstream age modeling. Together, single-cell and bulk data indicate that SBS1 accumulation is active in premalignant cells and accelerates during tumor evolution, with a substantial fraction of the excess present in the tumor’s MRCA.

### Age estimates place HRD onset years before diagnosis

Converting HRD onset from molecular (SBS1-based) time to chronological age requires a model relating SBS1 mutation burden to age. SBS1 mutations accumulate approximately linearly with age in normal tissue, yet breast tumors harbor a substantial excess of SBS1 mutations relative to normal mammary epithelium. This excess implies an increase in the rate of SBS1 accumulation during tumor development, and assuming a single lifelong rate would systematically lead to estimates of earlier age of HRD onset. We therefore constructed two SBS1-to-age models: the two-step model and the gradual model.

Both models are defined by two shared parameters (Figure 5a, Supplementary Note 2). The first is the early SBS1 accumulation rate, measured in normal mammary epithelial cells. The second is the late, tumor-associated rate, calibrated such that the trajectory reaches each tumor’s observed SBS1 burden at diagnosis. The late SBS1 rate is inferred over the tumor growth period— the interval between tumor initiation and diagnosis—using subtype-specific doubling times estimated from serial mammograms (Methods). Because triple-negative breast cancers (TNBCs) grow faster and over shorter periods than ER+ tumors, this interval is subtype-dependent (∼10.4 years for TNBC versus ∼16.3 years for ER+). The two models differ only in how SBS1 accumulation transitions between the early and late rates across this interval. In both cases, we propagate uncertainty in parameters and incorporate a fixed post-MRCA contribution of 116 SBS1 mutations per diploid genome, derived from scNanoSeq data, to account for cell-private mutations not detectable in bulk sequencing (Figure 4e).

**Figure 5.**
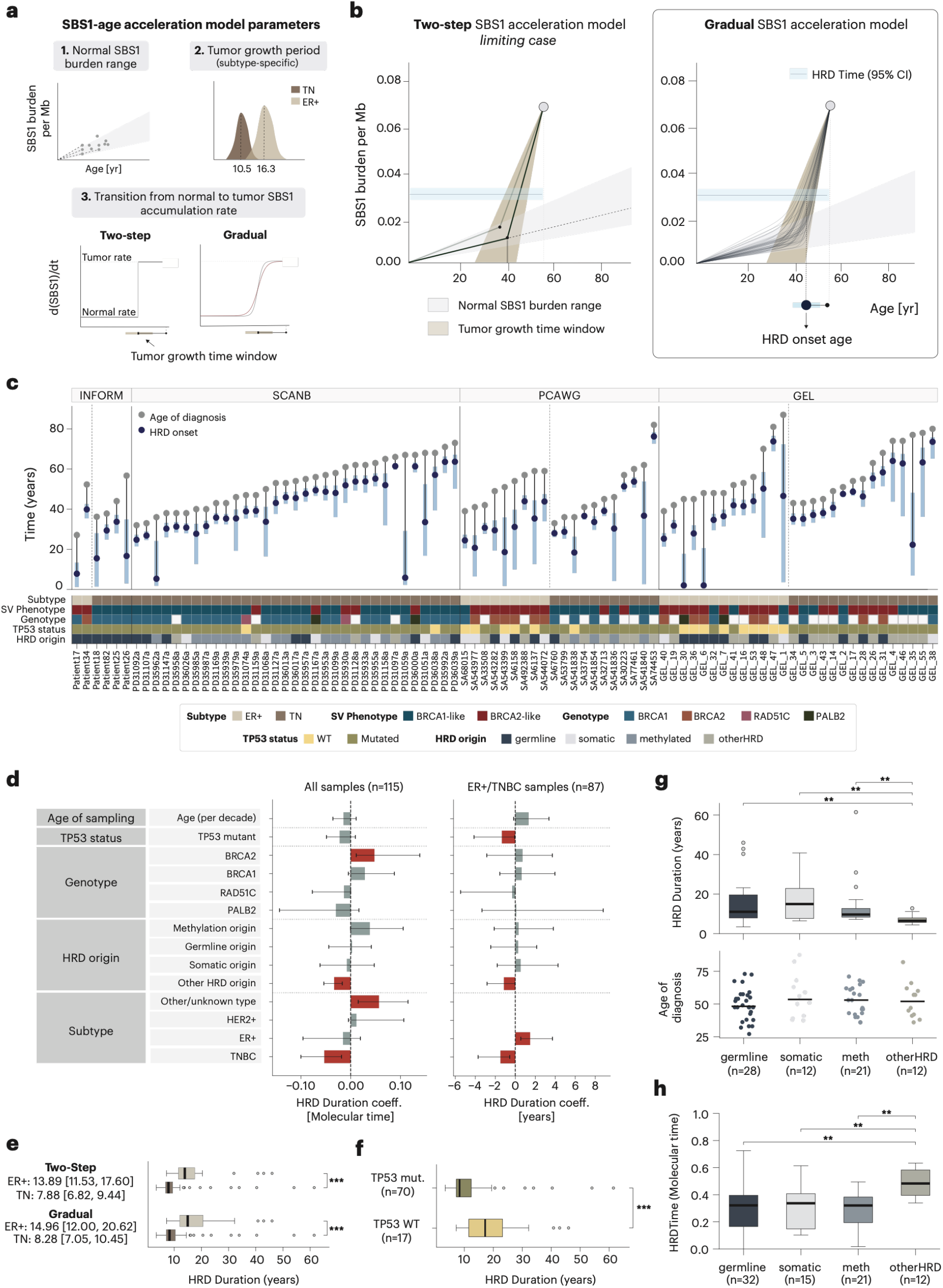
Age-resolved timing of HRD onset in breast cancer and its association with clinical and molecular features. **(a)** Parameters of the SBS1-to-age models: normal SBS1 burden accumulation with age and subtype-specific tumor growth periods (TN: 10.4 years; ER+: 16.3 years). These parameters are used by the two alternative SBS1-to-age models with different rate transitions: instantaneous (Two-step) or sigmoidal (Gradual). **(b)** Conversion of SBS1-based molecular time into calendar age under the Two-step and Gradual SBS1 acceleration models, with uncertainty propagated at every step (Supplementary Note 2). **(c)** Age at diagnosis (gray) and the age of HRD onset inferred using the Gradual model (blue, 95% CI) for 87 tumors across INFORM, SCANB, PCAWG, and GEL cohorts. Tracks below show the receptor sub-type, SV phenotype (Figure S19), HRD genotype, TP53 status, and HRD origin. **(d)** Multivariable ridge regression coefficients (95% CI) of HRD duration against clinical and molecular covariates in all samples (n = 115, HRD duration in molecular time) and samples with known receptor status (n = 87, HRD duration in years). Confidence intervals were estimated by 500 bootstrap resampling iterations; coefficients highlighted in red are considered statistically significant based on 95% bootstrap CIs excluding zero. **(e)** Univariate comparison of HRD duration (years) by receptor subtype under both transition models. ER+ tumors show longer HRD duration than TNBC under both Two-step (13.9 versus 7.9 years) and Gradual (15.0 versus 8.3 years) models. Significance: ***P < 0.001. **(f)** Univariate comparison showing TP53-mutant tumors (n = 70) have shorter HRD duration than TP53-WT (n = 17). Significance: ***P < 0.001. **(g)** HRD duration in years (top) and age at diagnosis (bottom) by HRD origin. Other/unknown HRD tumors show shorter duration compared to tumors with germline, somatic, and methylation alterations of HR genes. Significance: *P < 0.05, **P < 0.01. **(h)** t_HRD_ (SBS1-based molecular time of HRD onset) is later in other/unknown-origin HRD tumors than in those with known HR (epi)mutations. Significance: **P < 0.01. Box plots show median, IQR, and 1.5 × IQR whiskers; two-sided Wilcoxon rank-sum tests were used throughout.

The two-step model assumes an abrupt change from the early to the late mutation rate at tumor initiation; although biologically unrealistic, represents an upper-bound scenario of instantaneous acceleration. Because multiple oncogenic events more plausibly drive a progressive, rather than abrupt, increase in proliferation and SBS1 accumulation, the gradual model replaces this step change with a smooth sigmoidal transition between the same two rates, with steepness and midpoint of the sigmoid constrained by the tumor growth period and the observed SBS1 excess at diagnosis (Figures S16–S18, Supplementary Note 2). We applied both models to convert the SBS1-based HRD onset estimates into calendar age.

The two models yielded convergent estimates of HRD duration before diagnosis that varied by subtype. For TNBCs, we estimated 8.3 years (IQR: 7.1–10.4) under the gradual model and 7.9 years (IQR: 6.8–9.4) under the two-step model. For ER+ tumors, HRD mutagenesis was estimated to last 15.0 years (IQR: 12.0–20.6) and 13.9 years (IQR: 11.5–17.6) under the gradual and two-step models, respectively. By contrast, a linear model assuming a constant lifelong SBS1 rate implied a much longer HRD duration (median 36.7 years), underscoring the substantial impact of SBS1 acceleration and the need to model it explicitly. The approximately two-fold longer HRD duration in ER+ than in TNBC tumors was consistent across both models.

To identify determinants of HRD duration, we fitted multivariate ridge regression models relating the duration of HRD mutagenesis before diagnosis to tumor features (in calendar years, as estimated by the gradual model). Receptor status was the dominant correlate: ER+ tumors showed longer HRD duration, whereas TNBC status was associated with shorter duration (Figure 5e). This association persisted, albeit attenuated, when HRD duration was expressed in SBS1-based molecular time, in which ER+ tumors likewise showed significantly earlier HRD onset, indicating that this association is not merely an artifact of subtype-specific SBS1-to-age modeling (Figure 5d). *TP53*-wild-type tumors, enriched among ER+ cases (14 of 17), showed longer HRD duration in univariate analysis (P = 1.1 × 10^−4^; Figure 5f). In the multivariable regression, *TP53* mutation remained associated with shorter HRD duration in calendar time, reaching borderline significance (Figure 5d).

We investigated whether the type of lesion that caused HRD was a determinant of its duration. Among tumors with an identifiable HR-pathway lesion, duration was comparable whether deficiency arose through germline *BRCA1/2* mutation, somatic mutation, or promoter hypermethylation of *BRCA1* or *RAD51C* (Figures 5g, 5h). Furthermore, tumors of unexplained origin (lacking any germline, somatic, or epigenetic HR-pathway defect) sustained HRD for significantly shorter periods (median 6.5 years vs. 9.7–15.0 years; all P < 0.01, BH-corrected) than any of the other three groups. The unexplained cases may reflect a distinct or later-acting route to HRD.

Finally, the inferred duration of HRD activity was marginally shorter than the estimated tumor growth period: 8.3 years of HRD mutagenesis (gradual model) for TNBC tumors projected to grow for 10.4 years on average, and 15.0 years for ER+ tumors estimated to grow for 16.3 years. This concordance suggests that HRD onset typically coincides with, or shortly follows, the initiation of the clonal expansions that give rise to the tumor.

### The HRD signature is rare in cells outside of the tumor lineage

HRDTimer predicts that HRD lineages can emerge years before diagnosis, preceded by a latent phase during which HRD-associated mutagenesis was absent or slow. HRD cells could therefore already be present within premalignant tissue. However, to our knowledge, single-cell DNA sequencing with sufficient accuracy to resolve cell-private mutational signatures has not been applied to premalignant breast tissue from cancer-free women. We thus re-analyzed tumor and nontumor cells from breast cancer samples profiled with scNanoSeq, using non-tumor cells as a proxy for premalignant tissue.

Conventional HRD classification relies on the combined analysis of point mutations, indels, structural variants, and copy number alterations.^5^ Because scNanoSeq has limited per-cell sensitivity (median somatic mutation detection power of 2.5%) and does not capture structural variants, we developed HRDscout, a single-cell HRD classifier, trained on bulk WGS tumor data downsampled to the mutation-detection sensitivity of scNanoSeq. In cross-validation, HRDscout robustly distinguished HRD from HR-proficient tumors, achieving robust performance at scNanoSeq-relevant mutation detection sensitivity (see Methods, ROC-AUC=0.98–0.99; PR-AUC = 0.98 compared to HRD prevalence of 0.37; Figures 6a, S20; Supplementary Table S6). At a fixed 2% false-positive rate, HRD sensitivity was 83–88% with a false discovery rate (FDR) of 3–4%. Model predictions were primarily driven by indels with microhomology (Figure 6b), a hallmark of HRD.^5^

**Figure 6.**
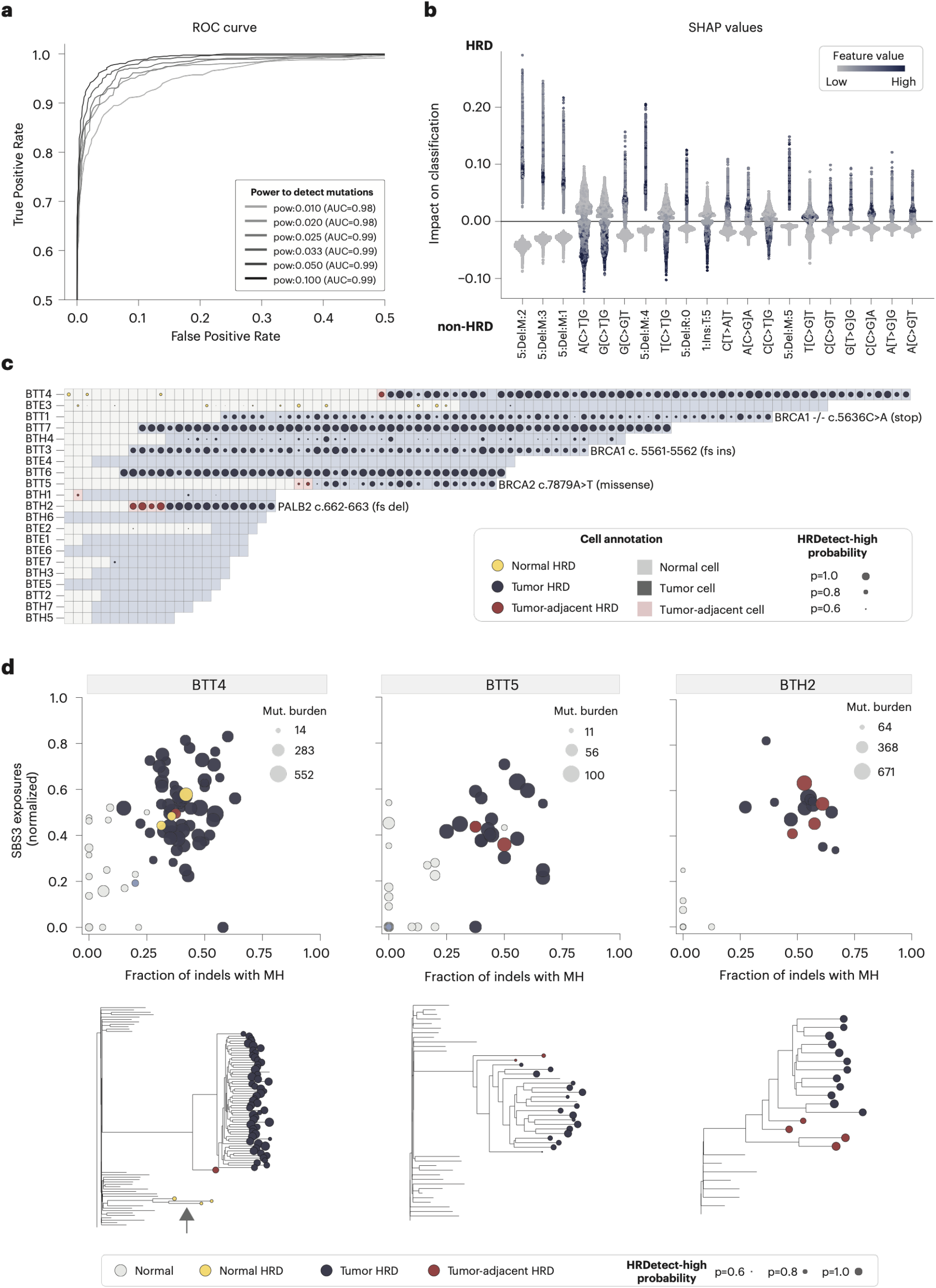
HRD classification of single cells profiled by scNanoSeq. **(a)** Receiver operating characteristic (ROC) curves for HRDscout, an HRD classifier optimized for low mutation-detection sensitivity. In cross-validation, HRDscout achieves an AUC of 0.98 at scNanoSeq-relevant mutation-detection sensitivity. **(b)** SHAP values identify the top features driving classifier performance, including short deletions with breakpoint microhomology (e.g., 5:Del:M:2, a 5 bp deletion with 2 bp microhomology) and point mutations in specific contexts (e.g., G[T>C]G). **(c)** Per-cell HRD classifications for tumor and non-tumor cells. Tumor cells from four germline mutation carriers (highlighted) consistently carry the HRD signature, as do cells from three additional tumors without known germline mutations. A small fraction of non-tumor cells is also confidently classified as HRD. **(d)** Top: HRD-associated signature exposures (SBS3, y axis) and indels with microhomology (MH, x axis) per cell, with HRDscout classifications overlaid. HRDscout classifications agree with HRD-associated signature exposures derived from point mutation and indel data. These three tumor samples were selected for harboring HRD cells originally labeled as non-tumor. Bottom: matched phylogenetic trees showing HRD-classified cells in tumor (navy) and non-tumor (yellow) lineages. BTT4 contains a tumor-independent HRD-positive lineage (arrow). All lineage trees with HRD classifications are shown in Figures S21–S23.

We applied HRDscout to 21 breast cancers profiled by scNanoSeq (Figure 6c). In all four tumors from carriers of HR-gene mutations (two *BRCA1*, one *BRCA2*, one *PALB2*) the tumor cells defined via phylogenetic analysis were predominantly classified as HRD; three tumors without a known germline mutation also showed predominantly HRD-classified tumor cells (Figure 6c). These single-cell classifications agreed with conventional per-cell signature fitting of point mutations and indels (Figures 6d, S21–S24).

Consistent with HRDTimer’s prediction of delayed HRD onset, the vast majority of non-tumor cells lacked a detectable HRD signature, including those from carriers of germline pathogenic mutations in HR genes. Because HRDscout may miss cells that have only recently acquired HRD and thus lack a detectable mutational signature, we performed a more sensitive, sample-level pooled analysis of indels across HRDscout-negative non-tumor cells. Even at this enhanced level of sensitivity compared to per-cell classifications, in non-tumor (HRDscout negative) cells from HRD tumor samples, we found no enrichment of HRD-associated microhomology-mediated deletions (median 5.3% of long deletions with MH, IQR 3.8–5.9%). This level of microhomology-mediated deletions which was indistinguishable from cells of non-HRD samples (median 5.9%, IQR 3.9– 7.2%; two-sided Mann–Whitney P = 0.81), and both were markedly lower than HRD tumor cells (median 35.5%, IQR 33.8–40.9%; two-sided Mann–Whitney P = 6 × 10^−3^; Figure S25). A single confident exception was a clone of three related HRD-classified non-tumor cells from tumor BTT4. These cells met stringent HRDscout criteria and carried sufficient mutations to confirm the HRD signature using both point mutations and indels (Figure 6d). Notably, their copy number profiles were distinct from those of the tumor cells, indicating that they originated from separate phylogenetic lineages.^22^

Together, these findings indicate that HRD mutagenesis is largely confined to the tumor lineage and only rarely arises in non-tumor cells before overt tumor expansion.

## Discussion

When HRD begins to shape an emerging tumor lineage has, until now, been unknown. Using SBS1 signature as a molecular clock and the activity of the HRD-attributed SBS3 as a second HRD-specific clock, HRDTimer dates the onset of HRD on both a molecular and a calendar time-scale. Across four breast cancer cohorts, HRD onset preceded WGD and arose post-zygotically, at a median of 34% of SBS1-based molecular time, even in carriers of germline *BRCA1/2* mutations. Translated to calendar time, HRD-associated mutagenesis was already active a median of 8.3 years (IQR 7.1–10.4) before diagnosis in triple-negative breast cancers and 15.0 years (IQR 12.0–20.6) in ER-positive tumors. These estimates reveal a previously unrecognized, clinically silent interval—on the order of a decade—during which an HRD lineage exists yet remains undetected.

The timing of HRD relative to other early genomic events suggests a recurrent evolutionary trajectory in these tumors. In tumors that underwent WGD, HRD consistently preceded the genome-doubling event, and where timing was possible, *TP53* mutations almost always occurred beforehand as well (43 of 44 cases; data not shown). Together, these findings support a model in which *TP53* inactivation and bi-allelic loss of *BRCA1/2* generate a viable HRD lineage, followed by genome doubling and clonal expansion. However, the *TP53*-wild-type HRD tumors are enriched among ER-positive cancers and show the longest HRD durations. Their presence indicates that alternative evolutionary routes to HRD also exist.

Beyond establishing when HRD begins, we identified factors that influence how long it persists before diagnosis. Receptor status emerged as the strongest determinant: ER-positive tumors exhibited substantially longer HRD durations than triple-negative tumors, a difference that was evident in both molecular and chronological time and therefore cannot be attributed solely to our subtype-specific SBS1-to-age model. The most parsimonious explanation is that ER-positive tumors progress more slowly from an HRD lesion to a clinically detectable cancer, thereby extending the observable HRD interval. Surprisingly, the duration of HRD was broadly similar whether the deficiency arose through germline mutation, somatic mutation or promoter hypermethylation of *BRCA1* or *RAD51C*. The similar HRD durations in hereditary and sporadic tumor lineages suggest that, once established, HRD lineages follow broadly comparable trajectories regardless of the initiating mechanism. In germline carriers, comparable HRD durations further indicate that HRD remains lineage-restricted even in this context. On its own, it does not account for their markedly elevated, age-skewed cancer risk.^1^ One possible explanation is that germline mutations—rather than altering the duration of HRD once established—increase the likelihood of HRD initiation, as fewer additional events are required for bi-allelic inactivation. If so, this could contribute to the earlier onset of HRD-associated cancers in germline carriers without necessitating a longer interval between HRD initiation and cancer diagnosis.

Placing HRD onset on a calendar timescale required accounting for a fundamental property of the SBS1-based molecular clock: its rate of accumulation accelerates during tumor evolution. Tumors carried a median 3.8-fold excess of clonal SBS1 mutations relative to age-matched normal mammary epithelium, and single-cell duplex sequencing data indicated that a substantial proportion of this acceleration occurs before the tumor’s most recent common ancestor. We therefore modeled the transition from normal to neoplastic SBS1 rates using measurements from both normal mammary epithelial cells and tumor samples, while propagating uncertainty in all underlying parameters to infer the relationship between SBS1 burden and chronological age. Estimates obtained under a conservative two-step model and a more biologically realistic gradual-acceleration model were highly concordant, whereas a naive constant SBS1-rate assumption would have overestimated HRD duration by approximately four-fold (median 36.7 years). More broadly, this framework extends molecular-time approaches from ordering genomic events to dating the emergence of a specific mutational process.

Single-cell duplex sequencing allowed us to determine which cells within the tumor and its microenvironment carry the HRD signature. Across 21 breast cancers, HRD-associated mutagenesis was largely confined to the tumor lineage: the vast majority of non-tumor cells lacked a detectable HRD signature, including cells from carriers of pathogenic germline variants. The sole confident exception was a small phylogenetically distinct non-tumor clone identified in a single case. This lineage restriction is consistent with the latent pre-HRD phase inferred by HRDTimer in tumor lineages. However, the conclusion about lineage specificity is constrained by the limited cellular sampling achievable with current single-cell sequencing. Although hundreds of non-tumor cells were examined, this represents only a minute fraction of the epithelial cells at risk of transformation, and the mutation-detection sensitivity of scNanoSeq could miss cells that acquired HRD too recently to accumulate a detectable signature. Nevertheless, the absence of microhomology-mediated indel enrichment argues against a large hidden population of HRD cells. The rarity of a full HRD signature also does not exclude more subtle haploinsufficiency-associated phenotypes.^6,8^ Clarifying these possibilities will require larger single-cell studies that couple mutational profiling with direct cell-type annotation and lineage reconstruction.

HRDTimer is broadly applicable but limited by available data. The framework uses WGD as a shared temporal anchor, yet in tumors without WGD, effective-genome-size–normalized SBS3 burden matches that in WGD tumors, consistent with similar HRD durations, although their precise onset times cannot be dated (Figure S26). Converting molecular time to calendar age also depends on tissue-specific baseline mutation rates: ovarian cancers show similarly early HRD onset in molecular time (median 0.29), but the lack of somatic mutation-rate estimates for fallopian tube epithelium precludes direct age calibration, a gap emerging normal-tissue atlases will address.^33^ Directly timing the second *BRCA1/BRCA2* hit would refine the model, but most second hits occur via copy number loss, which cannot yet be temporally resolved. Finally, validating SBS1-based molecular clocks will require longitudinal sampling, as available HRD cell-line data accumulate too few SBS1 mutations to confirm the SBS3:SBS1 relative mutation rate over time.

HRD-deficient cells rely on error-prone DNA repair, a vulnerability already exploited therapeutically in the clinic.^34^ By dating HRD emergence and showing that HRD lineages can persist in a lineage-restricted manner for years before diagnosis, our analysis revealed an extended premalignant window during which HRD states may be detectable. As genomic surveillance of high-risk individuals matures, the duration and timing of HRD mutagenesis could serve as a dynamic biomarker of cancer risk, enabling more precise, individualized interception that may reduce reliance on irreversible risk-reducing surgery.

## Methods

### Datasets

#### PCAWG

From the PCAWG dataset of 2,778 cancer samples, we selected 61 breast and ovarian cancers with HRD signature (HRDetect > 0.7) and evidence of whole-genome doubling (evaluated by MutationTimeR) for the main analysis; of these, 20 breast and 27 ovarian cancers passed all HRDTimer quality-control filters (see “Sample quality control and curation before applying HRDTimer”) and entered the main analysis. An additional 190 breast cancers lacking either HRD signatures or genome doubling were included for comparative mutational signature analysis and for estimating the SBS1 exposure in tumor samples compared to premalignant samples. We used the consensus calls obtained from multiple single nucleotide variant callers produced by the PCAWG consortium, where evidence from at least 2 out of 4 callers (CaVEMan v1.5.1, MuTect, MuSE v1.0rc, DKFZ) was required per variant.^15^ Somatic mutation calls from the PCAWG study (.vcf files dated ‘20160830’) were accessed through the ICGC data portal, and we included all mutations with the ‘PASS’ flag.^36^ We used consensus PCAWG indels derived by merging Platypus (v0.7.4), cgpPindel (v1.5.7), and SvABA calls.^15^ PCAWG’s consensus allele-specific copy number profiles dated ‘20170119’, as downloaded from the ICGC data portal, were used for timing mutations. To distinguish early from late structural variants, we accessed the structural variant and copy number variant calls generated by the Hartwig Medical Foundation using Gridss (v2.9.3) and Purple (v2.53) algorithms.^37,38^ The annotation of pathogenic mutations and the classification of the samples as HRD were retrieved from publications.^5,39^,40 As HRD, we considered samples with HRDetect probability over 0.7, according to published probabilities^.16,39^ The ICGC data portal has since been retired, but somatic mutation calls remain accessible from the ICGC Argo website.^41^ We also make the somatic mutation calls available for visualization with Chromoscope (see Data Availability section).

#### SCANB

Point mutations, indels, and copy number profiles were accessed from the Mendeley repository.^16,42^ Copy number profiles had been estimated using both the Battenberg and Ascat algorithms. We used a previously published classification of pathogenic mutations in HR genes, as well as HRD classifications using mutational signatures and HRDetect.^16^ Of the 233 published TNBC samples, we included those meeting the following criteria: consistent clonal ploidy estimates (within 0.2) between Ascat and Battenberg algorithms; tumor purity ≥ 0.45, based on Battenberg estimates; manual curation of copy number profiles to ensure closeness to integer levels, excluding samples with subclonality or contamination as flagged in the ‘Ascat.Curation’ column in the original publication;^16,42^ tumor whole-genome sequencing coverage ≥ 25x; HRD (HRDetect probability > 0.7); and genome doubling as evaluated by MutationTimeR’s classWGD function. A total of 49 SCANB samples met these eligibility criteria, of which 33 passed all HRDTimer qualitycontrol filters (see “Sample quality control and curation before applying HRDTimer”) and were included in the HRD timing analysis.

#### INFORM

The TBCRC-031 clinical trial (INFORM trial, NCT01670500) enrolled 118 patients and included patients with Stage I-III HER2-negative invasive breast cancer across 13 US sites. The trial evaluated different neoadjuvant chemotherapy regimens with curative intent.^17^ Each patient carried a germline pathogenic or likely pathogenic variant in *BRCA1* or *BRCA2* (*BRCA1/2*). Pretreatment tumor samples and matched normal blood were sequenced at a median effective whole-genome coverage of 38x (tumor). Somatic SNVs, indels, and copy number profiles were derived using Sentieon TNScope (v202112.01).^43^ Structural and copy number variants were called by Gridss (v2.13.2) and PURPLE (v3.8.1).^38^ Only samples with PURPLE QCStatus not marked as ‘noisy’ in PURPLE’s ‘QCStatus’ were included. Among 33 WGS tumor samples, 19 passed PURPLE QC. Of these, 8 exhibited genome doubling per MutationTimeR’s classWGD function, of which 6 passed all HRDTimer quality-control filters (see “Sample quality control and curation before applying HRDTimer”) and were retained for the main timing analysis. In addition to the pathogenic germline variants based on which the patients were enrolled, all the 8 wholegenome duplicated samples displayed the pattern of structural variants attributed to HRD, and thus were considered as HRD for the timing analysis.

#### ICGC matched primary-relapse samples

We accessed previously published whole genome data from seven breast cancer patients with longitudinal primary-relapse sampling.^21^ While the original study included data from more patients, we restricted our analysis to those profiled by whole-genome sequencing. Somatic mutation calls were obtained from the repository linked to the publication, together with annotations of total copy number and mutation multiplicity.^44^ No further filtering of the mutation set was performed.

#### Pre-malignant epithelial breast cells

Somatic point mutation data from 32 single-cell-derived organoids, obtained from 12 premenopausal women, were provided by the study authors.^20^

#### Single-cell nano-seq (scNanoSeq) data from 21 breast cancers

Mutation calls, read counts, unique DNA fragment counts, and copy number profiles from 781 cells were obtained from the authors of the study.^22^ This dataset included both tumor-lineage and non-tumor cells. Somatic mutation filtering differed from the original publication to better remove germline variants without distorting somatic mutational signatures (see “Single cell (scNanoSeq) re-analysis”). Duplex tag information was leveraged to generate an additional dataset of somatic indels for mutational signature analysis.

#### Filtering somatic mutations from bulk WGS to remove germline contamination

Filtering somatic mutations against population databases can disproportionately remove C>T mutations at CpG sites, one of common germline mutational signatures.^45^ Conversely, sample contamination can skew somatic mutational signatures. To enable cross-dataset comparison, we reprofiled each somatic dataset using the gnomAD (v2.1) germline reference, counting overlaps with gnomAD variants across decreasing population allele frequency thresholds (Figure S27).

SCANB somatic mutation calls were available in two versions: one filtered against population databases, and one unfiltered. In the filtered version, variants overlapping dbSNP v147, ExAC03, or dbNSFP3.0a germline datasets were excluded. Previously applied filtering markedly reduced SBS1 exposure estimates; thus, we used the unfiltered dataset (aside from normal panel filtering), as no evidence of germline contamination was found (Figure S27). We posit that filtering based on population databases containing rare variants (frequency < 0.001) inadvertently removes true somatic SBS1 mutations. Comparison against gnomAD variants enabled assessment of the degree of germline filtering applied across somatic mutation datasets (Figure S27).

### HRDTimer

#### Fitting mutational signatures

We fitted previously established breast cancer mutational signatures using the MuSiCal package, with likelihood optimization as the objective and a likelihood-based threshold for exposure sparsity.^28^ The following mutational signatures were considered, based on prior analysis of breast cancer samples: SBS1, 2, 3, 5, 8, 13, 17a, 17b, 18, 34, 51, 60 (COSMIC v3p2).^28^ For MuSiCal’s fitting function, we used a likelihood threshold of 0.001.

The probability that a mutation was generated by a signature was calculated from sample-level exposures and normalized vectors representing the abundance of mutations in each of the 96 channels^46^ via Bayes’ rule: P(S|C) = P(C|S)P(S)/P(C), where P(S|C) is the probability that a mutation in 3-nucleotide context C was generated by mutational signature S; P(C|S) is the probability that a mutational signature S generated a mutation in context C (known signatures normalized to 1); P(S) is the probability of a signature generating a mutation in a given sample (equivalent to its normalized exposure); and P(C) is obtained by marginalizing P(C|S)P(S).

#### Timing mutations with respect to gains and whole genome duplications

We first verified whether the copy number profile in each sample is consistent with a whole genome duplication, using MutationTimeR’s classWGD function that considers total ploidy and the proportion of the genome with loss of heterozygosity (2.9 – 2h ≤ p, where h is the proportion of genome with loss-of-heterozygosity and p is genome-wide ploidy; average total copy number).^27^ We timed somatic mutations against copy number gains, resulting in predictions of mutations as early or late, using allele-specific copy number profiles. In the first pass, we followed the MutationTimeR methodology and all mutations were included regardless of their probable signature of origin.^27,47^ The first pass estimated the timing of each copy number gain region separately, using the ‘mutationTime’ function from the ‘MutationTimeR’ package with 1000 bootstraps. We used the timing of copy number regions using all mutations to identify regions of gain whose timing is consistent, as suggested previously.^27^ Next, the timing of whole genome duplication was estimated through a search from 0 to 1, so that most of the genome has gains whose time was estimated in a consistent range. This procedure resulted in a set of copy number segments consistent with whole genome duplication (WGD) as a singular event before the most recent common ancestor cell (MRCA). HRDTimer uses only the mutations in the gain regions whose timing is consistent with the timing of the WGD.

#### Sample quality control and curation before applying HRDTimer

For each sample, we assessed whether the distribution of variant allele fraction was consistent with the theoretical distribution, given estimated sample purity and copy number profiles. We applied MutationTimeR’s recommended quality check by calculating the tail probability for each mutation.^27,47^ For each mutation, factors such as multiplicity, local coverage, total copy number, and sample purity determined the mean of the negative binomial distribution that models the number of supporting reads. Using the distribution defined by MutationTimeR for each mutation, we computed the tail probability based on the observed read support. In well-behaved samples with accurate purity and copy number estimates, about 1% of mutations are expected to have tail probabilities outside [0.005, 0.995], and about 5% outside [0.025, 0.975]. We excluded samples where a higher-than-expected fraction of mutations had tail probabilities outside these intervals, indicating likely errors in purity or copy number estimation.

We additionally excluded samples where the signature of homologous recombination deficiency could not be confidently determined. We relied on previously published HRDetect bootstrapping results for the PCAWG dataset and excluded samples with a 95% lower confidence bound for HRDetect below 0.7—a previously recommended value.^39^ In the absence of HRDetect bootstrapping results, we required that at least 40% of indels exhibited microhomology to classify a sample as confidently HRD.

HRDTimer depends on accurate estimation of the SBS3-to-SBS1 relative mutation rate, based on late mutations occurring after genome doubling. If genome doubling occurred relatively late, the resulting moderate number of post-doubling SBS1 and SBS3 mutations leads to uncertainty in relative mutation rates and the timing of HRD. Consequently, we excluded samples in which the confidence interval for late SBS1 exposure included zero in any of the mutation groups— early, late, or undefined (NA; MajorCN = 2, MinorCN = 1)—provided that each group contained a sufficient number of mutations. Additionally, we excluded samples where the mean estimated number of SBS1 mutations was less than 15, as these cases showed wide confidence intervals for the estimated HRD onset time. We excluded one sample (INFORM Patient 101) with wide 95% confidence intervals and a negative estimated time of HRD onset, as this suggests an elevated SBS3:SBS1 ratio prior to genome doubling—violating the assumption of a constant relative mutation rate since HRD onset. Figure S13 shows all the samples before filtering.

#### Estimates of molecular time of whole genome duplication

In regions consistent with WGD, each somatic mutation in a sample with whole genome duplication was assigned a timing probability (early, preceding WGD, or late, following WGD) and a probability of originating from a specific mutational process (SBS1, SBS3, or others). Specifically, the probability of a mutation *i* originating from SBS1, a process regarded as a molecular clock, is denoted as P_SBS1|i_, while the probabilities of a mutation being early and late are denoted as P_Gain|i_ and P_Single|i_ respectively. Each mutation was assigned a probabilistic weight representing the combined likelihood of being classified as early or late and originating from a specific mutational signature (for example, SBS1, SBS3, etc.). These weights were calculated separately for early mutations 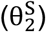 and late mutations 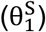 using the following approach:

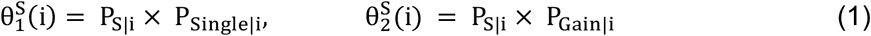

where S denotes the signature of interest, and the *P*_*S*|*i*_ values were determined based on estimated mutational signature exposures and the assignment of each mutation as early, late, or undefined (for regions where the major copy number is 2 and minor copy number is 1). By summing over all mutations *i*, the expected total number of early and late mutations attributed to a given mutational signature, 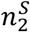 and 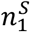 respectively, were calculated as:

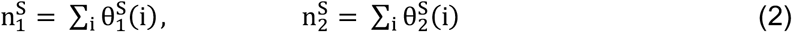

These expected counts of early and late mutations per signature were then used to establish the relative mutation rate.

Mutations could be directly timed only in gain regions, specifically where the total copy number is 4 with a minor copy number of 2, or where total copy number is 2 with a minor copy number of 0, indicating loss of heterozygosity. In regions with a total copy number of 3 and a minor copy number of 1, mutations on one copy cannot be unambiguously timed with respect to WGD, because early mutations detectable on one copy are possible. However, assuming that in (2,1) regions the number of early mutations is twice the detected number of mutations on two copies, the relative proportions of mutations on one and two chromosome copies can be used to estimate the timing of WGD and HRD.

For each sample and region with allele-specific copy numbers of (major = 2, minor = 0), (2,1) and (2,2), we calculated the proportions of early 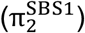 and late 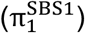 mutations out of all SBS1-attributed mutations (note that 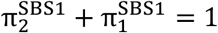). Since SBS1 is predominantly associated with [C>T]pG mutations, these proportions were calculated exclusively for [C>T]pGs, with weights reflecting the probability that each mutation originated from SBS1.

The proportions of late mutations (1 denotes multiplicity of mutations post WGD) among mutations attributed to signature SBS1 are defined by:

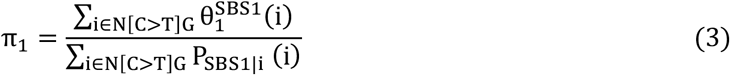

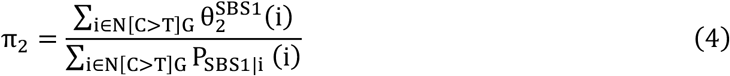

where N = {A, C, G, T}. This method improves upon previous approaches, which either treated all [C>T]pGs as a molecular clock or relied on maximum a posteriori assignment of mutations to signatures. The resulting *π* estimates were subsequently used to time WGD for each of the (2,2), (2,1), and (2,0) regions. After normalizing for ploidy across copy number regions following the MutationTimeR ploidy normalization45, the WGD timing estimates for the (2,0), (2,1), and (2,2) regions are:

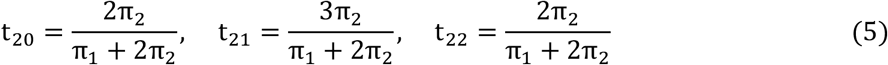

A final WGD estimate was obtained for each sample, similar to the MutationTimeR^47^ algorithm, as the weighted mean of the three individual estimates, with weights based on the proportion of SBS1-derived mutations in each copy number state, as shown in Eq. 6. The final WGD timing (t_WGD_) for a given sample was calculated as the mean of all individual bootstrapped estimates, with 95% confidence intervals as described in the section “Confidence intervals for the timing estimates using bootstrapping”.

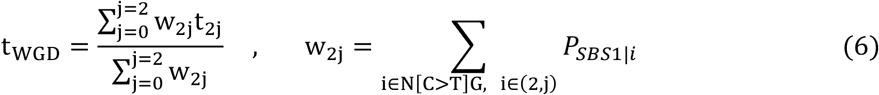

#### Estimates of molecular time of the HRD onset

The presence of early (pre-WGD) SBS3 mutations—a mutational signature associated with HRD—suggests that HRD onset preceded WGD. Adapting our previous approach used to estimate WGD timing, we first focused on (2,0) and (2,2) genomic regions, where mutations present in one copy of a chromosome can be unambiguously characterized as late. Assuming that the relative rate of SBS1 and SBS3 mutation acquisition, denoted as c, remains constant following HRD onset (see “Subclonal and cell-private SBS3:SBS1 ratio analysis”), we calculated this rate for late mutations as:

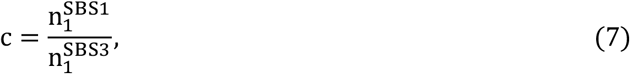

where 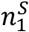 is defined in Eq. 2, and 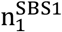, and 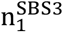 represent the expected late-stage mutations (post-WGD) attributed to SBS1 and SBS3, respectively.

To time the onset of HRD, the expected number of early SBS1 mutations 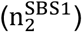 was adjusted by subtracting the contribution of “molecular-clock” mutations that occurred after the HRD onset and before the time of WGD. This adjustment ensures that the newly calculated expected number of SBS1-attributed mutations 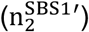 reflects only those that occurred prior to the HRD onset, providing an estimate of HRD onset:

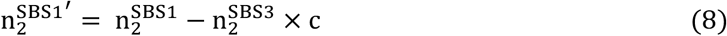

The prime notation (′) is consistently used throughout this section to denote variables modified to estimate the onset of HRD instead of WGD. To implement the HRDTimer workflow, Eq. 8 was normalized by the total expected SBS1 mutation burden and expressed as proportions (see the next section for derivation):

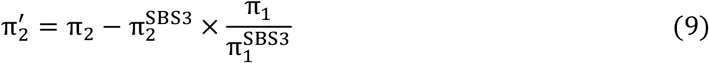

Here, the proportions of early 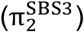 and late 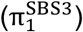 mutations out of all SBS3-attributed mutations were calculated similarly to SBS1, with the key distinction that all mutations, i, were included rather than exclusively [C>T]pGs:

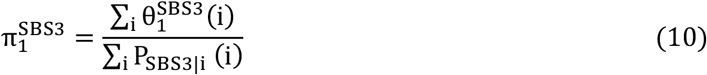

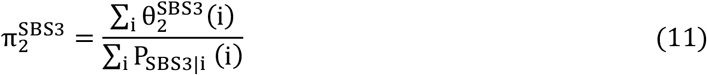

Eq. 9 was applied to the (2,0) and (2,2) regions. For the (2,1) regions, where late mutations cannot be confidently assigned, the average value of c from the (2,0) and (2,2) regions was used. In this case, Eq. 9 simplifies to:

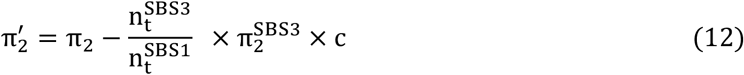

where 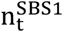 and 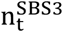 represent the total number of mutations attributed to SBS1 and SBS3, respectively (e.g., 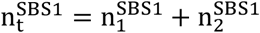). The ploidy-normalized timing estimate of HRD onset per copy number state was then calculated following the MutationTimeR ploidy correction:

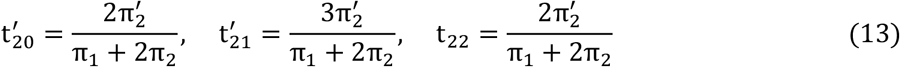

Using an approach analogous to that described in the section “Estimates of molecular time of whole genome duplication”, a single estimate for the timing of HRD onset was derived by calculating the weighted mean of individual timing estimates across copy number regions. As shown in Eq. 14, the weights correspond to the total number of mutations within each copy number region rather than exclusively N[C>T]pGs. To estimate the confidence interval (CI) for HRD timing we used bootstrapping, as described in the section “Confidence intervals for the timing estimates using bootstrapping” below.

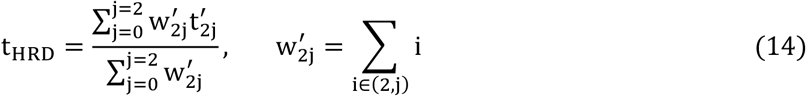

##### Derivation

Using the result obtained in Eq. 8, we express the adjusted expected number of early SBS1 mutations as:

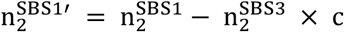

where the correction factor c is defined as:

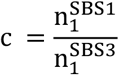

To convert this to a normalized proportion, we divide both sides by the total expected number of SBS1 mutations, 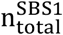, yielding:

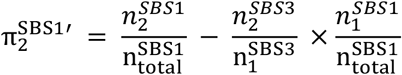

Here, n denotes the expected number of early or late mutations attributed to the relevant mutational signatures. Using the following definitions:

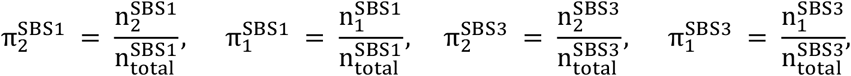

we arrive at the final expression:

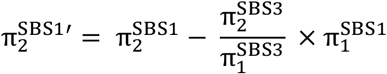

#### Confidence intervals for the timing estimates using bootstrapping

Mutations from each sample are categorized into three subgroups—Early, Late, and Undefined—based on the maximum a posteriori (MAP) timing estimates obtained using MutationTimeR. These subgroups constitute the reference mutation sets for downstream analysis. In a single iteration of the HRDTimer signature-analysis module, each subgroup is analyzed independently to estimate the probabilities of signature attribution of a given mutation type (e.g., A[C>T]G) to a mutational signature (e.g., SBS1), thereby capturing distinct mutational trends across temporal phases.

To assess the robustness of our timing estimates and quantify uncertainty due to variability in signature attribution and the observed mutational profiles, we employ a resampling-based strategy (Figure S9). For each subgroup (Early, Late, and Undefined), the observed mutation profile is normalized to create a probability vector, which serves as the mean for multinomial sampling. New mutation profiles are then generated by sampling from the multinomial distribution; estimation of signature abundance, signature attribution, and posterior probability calculations are performed for each resampled subset, and the resulting signature probabilities are applied back to the original mutation set to recompute timing estimates. This approach isolates variation to the signature attribution step, allowing us to assess how fluctuations in the inferred signature probability vectors influence downstream timing estimates. While the mutation sets and their timing remain fixed, the probabilistic sampling and downstream signature analysis inherently introduce variability in the mutation profiles, capturing the uncertainty present in the observed data.

This procedure is repeated 200 times. The reported timing estimates for t_WGD_ and t_HRD_ represent the mean across resampling iterations, with 95% confidence intervals computed to reflect the uncertainty in the assignment of mutations to signatures.

Of note, the per-mutation timing probabilities from MutationTimeR depend only on the variant allele fraction of a mutation, sample purity, and local copy number. Only the sample purity and local copy number are subject to uncertainty that could affect the timing of somatic mutations. To ensure that purity and copy number estimates are robust, we only used samples where multiple copy number algorithms agreed on local copy number and sample purity (see “Datasets”). We additionally excluded samples where MutationTimeR’s QC metric indicated a possible mismatch between observed mutation allele fractions and fitted purity and copy number profile (see “Sample quality control and curation before applying HRDTimer”).

#### Timing of HRD-attributed indels and structural variants from bulk WGS data

Small insertions and deletions (indels) were classified as early or late based on the variant allele fraction using MutationTimeR and variant allele fractions reported in PCAWG consensus indel files (Figure S7).^27^

To time structural variants in the PCAWG dataset, we used allele-specific copy number profiles from the Purple algorithm that also include information on adjacent structural variants from Gridss.^37,38^ Allele-specific copy number profiles include estimates of total copy number at a position, as well as estimates of the number of copies of the maternally- and paternally-inherited chromosomes. In the absence of sequencing data from both parents, we cannot assign parental origin, but can instead distinguish the major chromosome copy, the one that is present in more copies, and the minor one. Purple and Gridss algorithms cross-check structural and copy number information, resulting in high-resolution copy number profiles around structural variants. The copy number profile and changes allow us to identify structural variants in regions of gain that can be timed with respect to genome doubling. We also verify whether the copy number change at the SV breakpoint is consistent with the SV type.

From the copy number variants found by Purple, we selected those with the following characteristics (Figure S7):

- Supported by a structural variant compatible with the HRD signature, such as a deletion or tandem duplication with a genomic footprint smaller than 100 kb.
- The background copy number around the SV is compatible with clonal gain, to enable timing of the SV in that region:
  ∘ The background major copy number is between 1.7 and 2.3, indicating that the broader region was previously amplified and allowing the timing of smaller variants relative to that gain.
  ∘ The minor background copy number deviates by no more than 0.3 from the nearest integer, and the minor copy number on either side of the structural variant differs by less than 0.3, consistent with a uniform allele-specific copy number profile across the affected region. The background minor copy number for each structural variant was calculated as an average of the minor copy number to the left and right of the structural variant.
- The copy number step at SV breakpoints is consistent with a simple structural variant:
  ∘ The copy number step ratio between the left and right breakpoints (s_l_/s_r_, where s_l_ and s_r_ represent the copy number step on the left and right sides of a structural variant, respectively) ranged within (0.7,1.3), suggesting that both sides of the structural variant experienced copy number changes of similar magnitude and opposite direction.
  ∘ The average absolute value of the copy number step was between 0 and 2.

We considered variants where the rounded absolute value of the copy number step was 2 as early and where the rounded absolute value of the copy number step was 1 as late. We then divided such variants according to the background minor copy number (0, 1, 2 after rounding), and estimated the proportion of late variants for each of the three considered copy number configurations in the region: (total=2, minor=0), (2,1), (2,2) (Figure S7).

### Translating the molecular SBS1 time to human age

#### Normalization of detected mutational burden

A common assumption is that chromosomal gains and genome doubling increase the potential for accumulating somatic mutations. Although the exact timing of chromosomal gains is typically unknown, the distribution of mutation multiplicities across detected mutations can capture information about the relative timing of these gains.

To reflect ‘historical average ploidy’, we define the concept of an ‘effective genome size’, which is estimated based on the multiplicity of mutations.^27^ Because the SBS1 signature is thought to be active continuously throughout life, whereas other mutational processes may be episodic, we estimated the effective genome size specifically using mutations attributed to SBS1 with high probability.

For bulk whole-genome sequencing (WGS) samples, the effective genome size G was computed as:

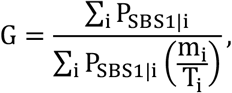

where P_SBS1|i_ is the probability that mutation i was generated by the signature SBS1, m_i_ is the multiplicity of mutation i (i.e., the number of chromosome copies carrying the mutation), and T_i_ is the total copy number at the mutation locus.

The observed total mutation burden n was then rescaled to a diploid-equivalent burden n^′^ using:

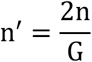

#### Correlates of SBS1 burden in bulk cancer samples

To assess covariates of SBS1 burden, we fitted a multivariable ridge regression of scaled SBS1 on age at diagnosis (categorical, per decade), IHC subtype (ER+, HER2+, TNBC; reference: ER+), tumor grade (G1, G2, G3; reference: G2), and HRD status (HRD/non-HRD; reference: HRD). Ridge regression was used to account for collinearity among covariates. To investigate differences between specific groups, an equivalent OLS model with an identical design matrix was fitted, from which effect sizes and P-values are reported.

#### Functions connecting molecular time to age

SBS1 accumulates approximately linearly with age in normal mammary epithelium but is present in excess in breast cancers (Figure 3). Assuming a constant SBS1 accumulation rate would therefore place HRD onset unrealistically early in life. To account for accelerated SBS1 accumulation during tumor development, we developed two SBS1–age acceleration models, together with a constant-rate linear model as a comparator (full details in Supplementary Note 2).

#### Estimation of SBS1 burden at diagnosis

Bulk whole-genome sequencing measures the SBS1 burden in the most recent common ancestor (MRCA), Y_MRCA_; to estimate the SBS1 burden at diagnosis we added a correction for cell-private SBS1 mutations derived from scNanoSeq data:

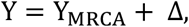

where Δ = 116 mutations (diploid-genome normalized) corresponds to the median excess SBS1 burden observed in single cells relative to the MRCA (Figure 4).

#### Acceleration models

To account for accelerated SBS1 accumulation during tumor development, we considered two SBS1–age acceleration models. Both acceleration models assume an early phase with a normal-cell accumulation rate (s_norm_) and a later phase with a higher tumorspecific rate (s_tumor_). The *two-step model* assumes an abrupt transition between these rates at tumor initiation and provides an upper bound on SBS1 acceleration. The *gradual model*, used as our primary model, replaces this abrupt transition with a sigmoidal increase from s_norm_ to s_tumor_.

#### Early and late SBS1 rates

The early accumulation rate, s_norm_, was estimated from normal breast epithelial cells. Rather than fixing s_norm_ to a single point estimate, we sampled from the range of rates observed across normal cells (see “Uncertainty propagation”). For each tumor, the late accumulation rate, s_tumor_, was chosen such that the cumulative SBS1 burden reached the observed burden at diagnosis (Y) over a subtype-specific tumor growth period d (ER+ ≈16.3 yr; TN ≈10.4 yr), derived from ultrasonography-based tumor doubling times.^48^ Under the *two-step model*:

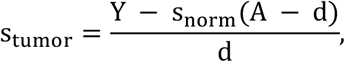

where A is the age at diagnosis. The gradual model uses the same boundary conditions but transitions smoothly between the two rates.

#### Uncertainty propagation

Uncertainty in the normal-cell SBS1 rate, subtype-specific tumor growth period, rate transition parameters, and SBS1-based HRD molecular-time estimate was propagated by sampling model parameters and generating N = 1000 SBS1–age functions per sample.

#### HRD onset age

The molecular time of HRD onset, t_HRD_, represents the fraction of the total SBS1 burden accumulated by the time HRD begins, corresponding to an SBS1 burden of t_HRD_Y. The age of HRD onset, a_c_, is defined as the age at which the SBS1–age function reaches this burden, such that y(a_c_) = t_HRD_Y. Solving this across the N = 1000 sampled functions yields per-sample estimates of a_c_ and the corresponding HRD duration, A − a_c_. Estimates are reported as the median with a 5–95% interval.

#### Estimates of chronological duration of tumor growth in years

The period of tumor growth (b) is estimated from tumor doubling time statistics published by others.^11^

where:

- T_d_ is the tumor doubling time (in days)
- T_s_ is the tumor size at diagnosis (in millimeters)

**Table 1.**
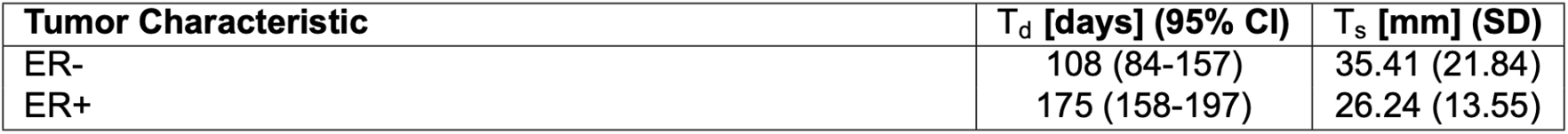
Tumor characteristics with corresponding doubling times (T_d_) and diameters (T_s_), s for size. ^11^ (SD = standard deviation)

| Tumor Characteristic | $T_d$ [days] (95% CI) | $T_s$ [mm] (SD) |
| --- | --- | --- |
| ER- | 108 (84-157) | 35.41 (21.84) |
| ER+ | 175 (158-197) | 26.24 (13.55) |

#### Derivation of tumor growth time from a single cell

We estimate the time d (in days) it takes for a tumor to grow from a single cell to a clinically detectable size, assuming spherical tumor growth from a single cell. The volume of a sphere is:

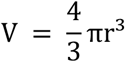

The initial diameter of a single tumor cell is defined as D_0_ = 0.01 mm (corresponding to 10 μm). The tumor diameter at diagnosis is denoted as D_s_ = T_s_ mm. The corresponding radii are 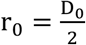 and 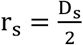. The number of cell doublings, n, required for a single cell to grow into a tumor of diameter D_s_, is given by:

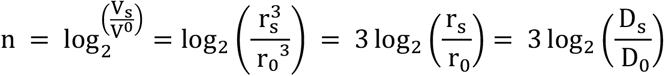

If each volume doubling takes T_d_ days, then the total time of tumor growth is:

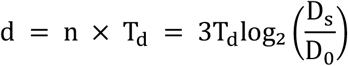

This model leads to an estimated period of approximately **10.4 years of tumor growth for an ER− tumor**, and **16.3 years for an ER+ tumor**.

### Robustness of HRDTimer

HRDTimer inherits its early/late mutation-timing classification from MutationTimeR, whose accuracy was validated in the PCAWG study.^27^ The reliability of the resulting estimates depends primarily on three inputs: sample purity, the copy-number profile, and the assumption of a single, clonal WGD. Uncertainty in the remaining key parameters—mutation multiplicity, mutation-to-signature attribution, and the SBS1-to-age relationship (Supplementary Note 2)—is propagated by HRDTimer’s bootstrapping procedure (see “Confidence intervals for the timing estimates using bootstrapping”). We therefore assessed each of the three primary inputs as follows.

#### Purity

Sample purity was taken from copy-number analysis. To verify this choice, we ran MutationTimeR across a grid of purity values and calculated the post-EM VAF log-likelihood, confirming that it peaked at the purity derived from copy-number analysis and declined as purity deviated from it (Figure S14). Purities far from this value are rejected by MutationTimeR’s quality-control step, which flags samples on the basis of the tail probabilities of the assumed VAF distributions (see “Sample quality control and curation before applying HRDTimer”).

#### Copy number profile

To test sensitivity to copy-number calling error, we randomly perturbed up to 10% of segments by adding or subtracting one copy and re-ran the pipeline. Predicted HRD-onset estimates were essentially unchanged (Figure S15), demonstrating robustness to realistic noise in copy-number calls.

#### Complex CNVs and post-WGD gains

Timing was restricted to regions with major copy number 2 whose timing is consistent with WGD (see “Timing mutations with respect to gains and whole genome duplications”). Complex copy-number patterns, which typically yield copy numbers above 2, are excluded by this constraint. Gains arising after WGD are likewise excluded, both by the major copy number 2 requirement and by the per-gain check that each gain’s timing is consistent with the inferred WGD time.

#### Subsequent losses

Regions retaining major copy number 2 after a single copy loss were retained, with the loss accounted for in the timing formulae for the (2,0) configuration (see “Estimates of molecular time of whole genome duplication”). Regions with biallelic loss were excluded, as their major copy number is no longer 2.

#### Clonal splits and WGD as an early event

Inclusion required a single clonal WGD. In PCAWG, Battenberg,^49^ which can detect subclonal copy-number changes, found no evidence of subclonal WGD; consistent with this, somatic VAF distributions were dominated by late-clonal mutations, whereas parallel subclonal WGDs would instead produce post-WGD mutations at subclonal VAF. A secondary subclonal WGD in a smaller subclone cannot be excluded but would not affect conclusions drawn from the clonal period.

### Single cell (scNanoSeq) re-analysis

#### Classification of cells as tumor or normal

A lineage tree was obtained from cell genotypes using CellPhy.^22,32^ On the lineage tree, we selected a long trunk following the original publication,^22^ and all cells descending from that trunk were considered tumor cells.

#### CNV analysis

We accessed previously published single-cell copy number profiles generated using Gingko.^22^ Based on the cell-specific CNV profiles, we derived consensus truncal CNV profiles by retaining genomic regions where total copy number estimates were consistent across all tumor cells in a given sample—that is, regions where the standard deviation of the estimated copy number was less than 0.5.

#### Evaluating possible germline contamination

True somatic mutations identified via duplex sequencing data can be overwhelmed by germline mutations originating from contaminating DNA fragments. Because germline mutations greatly outnumber somatic mutations, even low-level contamination from another individual can significantly distort the data. To assess contamination, we overlapped somatic mutations with gnomAD variants across decreasing population allele frequencies, following the same procedure used for bulk data. Samples with a high overlap between somatic mutations and gnomAD variants were considered likely to be contaminated by DNA from another individual (Figures S28, S29). We therefore excluded cells with greater than 40% overlap between somatic mutations and gnomAD variants at a population allele frequency threshold of 0.001.

#### Differences in filtering strategy from the original publication

The following filters, which differ from those used in the original publication, ^22^ were applied to the point mutation data:

- Variants overlapping gnomAD v2.1 entries with population allele frequency >0.001 were removed—this differs from the use of dbSNP in the original publication which did not consider population allele fraction;
- Only somatic mutations with ≥20× coverage in matched normal samples were retained, compared to the 10× threshold used in the original study, to improve discrimination between somatic and germline variants.

#### Power calculations

Since scNanoSeq has limited power to detect somatic mutations in individual cells (Figure S30) and the observed mutation counts are confounded by copy number alterations and the variable timing of somatic events, we developed and validated a method to estimate the underlying mutational burden. Full details of the derivation and validation are provided in Supplementary Note 1.

#### Evaluating the sequence coverage of scNanoSeq

Since the genome sequencing coverage in scNanoSeq is sparse and mutations are called when present even in single DNA fragment, any coverage biases might profoundly affect the detected mutational profile and signature. We therefore assessed the uniformity of sequencing coverage across trinucleotide contexts, as compared to the reference human genome. We found that sequencing coverage was approximately uniform, with the exception of the GCA trinucleotide context, which was under-represented (Figure S32). This under-represented context (GCA) was confirmed to be part of the TGCA motif recognized by the restriction enzyme HpyCH4V, explaining the localized depletion in coverage near enzymatic cut sites. Given the relatively even coverage of trinucleotide contexts in scNanoSeq, we performed mutational signature analysis without applying a trinucleotide-based correction.

#### De-novo signature discovery

We extracted mutational signatures from point mutations and indels detected in the scNanoSeq dataset using three distinct approaches.

- by grouping mutations by cell, allowing overlaps between mutation sets from different cells. Indel signatures obtained using this method are shown in (Figure S25).
- by grouping mutations by edge on the phylogenetic tree, based on mutation assignments from CellPhy. In this approach, mutation sets are mutually exclusive. Mutational signatures derived from point mutation data are shown in (Figure S33).
- by combining branch-level mutation sets with somatic mutation data from 800 whole-genome sequenced breast cancer samples.

Across the different extraction methods, we identified a novel mutational signature (“Signature 3” or “SigA – SBS106 like”, Figure S33), which was abundant in one scNanoSeq breast cancer sample (BTE5), and present at low levels in other samples. The mechanism underlying this signature remains unknown. As it has not been observed in large cohorts, we cannot exclude the possibility that it reflects a rare exposure unique to the patient sample during library preparation. We further verified that the inclusion of this novel signature during branch-level signature fitting did not affect the estimated exposures of SBS3, a key signature of interest.

#### Mutational signature fitting on data from single cells

Mutational signature exposures were estimated using MuSiCal, either at the single-cell level for tumor cells or by aggregating nontumor cells, using a likelihood threshold of 0.001 (Figures S21–S23).

The per-cell power to detect mutations with scNanoSeq was typically 0.025 [IQR: 0.019–0.033] (Figure S30), resulting in mutation sets of comparable size to those obtained through exome sequencing. Through experiments described below, we found that although single-cell burden estimates were inherently noisy, the aggregated signature exposures across many cells converged to the true underlying values (Figure S2).

To evaluate the impact of detection sensitivity on signature fitting, we simulated reduced mutation detection by starting from a larger combined mutation set derived from all cells within a single sample. We first fitted breast cancer mutational signatures to the full mutation set. We then downsampled this set to a range of target sizes (x-axis) and measured the deviation of the inferred signature exposures from those estimated using the complete dataset (Figure S2). This procedure was repeated across multiple target mutation counts.

We found that signature exposures for SBS1 and SBS3 in smaller datasets closely matched those from the full dataset, with reduced variability as the number of simulated mutations increased. These findings motivated our approach: performing signature fitting at the single-cell level for tumor cells, while aggregating non-tumor cells—where the number of detected somatic mutations was lower—for joint analysis. For reference, the average number of detected mutations per tumor cell was 225, compared to 54 per non-tumor cell.

#### Estimates of per-cell burden of SBS1 signature

For each cell, our algorithm produces a mutation burden estimate corrected for the power to detect mutations (see Supplementary Note). This correction is applied only in genomic regions where the total copy number ranges from 1 to 5. If the cumulative size of these regions is c Mb, the mutation burden is scaled by a factor of 3000 / c. We further corrected mutation burden estimates for regions where somatic mutations could not be called due to insufficient coverage in matched normal samples (see above). Let d denote the proportion of germline variants with total coverage below 20×, where somatic mutations are not reliably distinguishable. In this case, the mutation burden was scaled by 1/(1 − d). We additionally scaled the mutation burden by effective genome size and multiplied by 2 to report burden per diploid genome.

The effective genome size, G, is defined as:

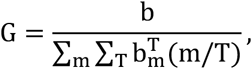

where b is the total mutation burden after power correction, and 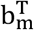 is the estimated burden of mutations with multiplicity m in regions with total copy number T. The power-corrected mutation burden b′ is calculated from the observed number of somatic mutations b as follows:

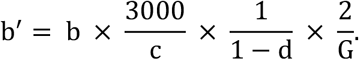

For tumor cells, where the number of mutations per cell is sufficient for reliable signature fitting, we use cell-specific normalized proportions of SBS1. For non-tumor cells, where mutation counts are generally too low for reliable signature fitting, we use SBS1 exposures estimated from pooled mutations across all non-tumor cells.

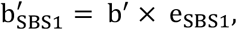

where e_SBS1_ denotes the normalized exposure of signature SBS1.

We also estimated the SBS1 mutation burden at the time of the most recent common ancestor (MRCA). This analysis was limited to samples for which deep bulk whole-genome sequencing data were available from the same tumor. Clonal mutations were identified in bulk samples based on their allele fractions.^19^ For each cell, we intersected the clonal mutations identified in bulk with those detected in single-cell data. The per-cell clonal mutation sets were power-corrected using the same procedure described above. SBS1 burden was then estimated by multiplying the corrected clonal mutation burden by the normalized SBS1 exposure.

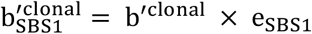

where e_SBS1_ is the normalized exposure of SBS1, obtained by fitting mutational signatures to the set of truncal mutations detected in single cells. The ability to define MRCA mutations depends on both the power to detect mutations and the number of sequenced cells. Therefore, matched bulk whole-genome sequencing was used to obtain more reliable estimates of MRCA burden and mutational signatures that do not depend on these factors.

#### Indel calling

Somatic insertions and deletions (indels) were identified using an analytical pipeline similar to that employed for somatic SNVs, with minor modifications. Briefly, (1) indels were called using samtools mpileup on the filtered a4s2 molecules. (2) Candidate indels were filtered using a custom Python script that required each indel to be supported by at least two reads from each DNA strand and to be located at least 8 base pairs from both ends of all supporting reads. (3) Putative somatic indels were then compared against the matched normal bulk sample using a custom Python script, which required: (i) a minimum coverage of 10 reads at the indel site in the matched normal bulk; (ii) no reads in the matched normal bulk supporting the candidate somatic indel; and (iii) the absence of any indels within 20 base pairs of the somatic indel. (4) Finally, candidate somatic indels were filtered against the dbSNP v20151104 database to exclude potential germline variants or cross-sample contaminations. Only indels meeting all these criteria were retained as high-confidence somatic indels.

### Classification of single cells as HRD (HRDscout)

Somatic point mutations and indels identified in each single cell were aggregated into mutation catalogs and served as input features for a classifier of homologous recombination deficiency (HRD).

#### Training

The random forest classifier was trained on bulk WGS data combining three previously published datasets: the breast cancer 560 cohort,^35^ SCANB,^16^ and the INFORM dataset,^17^ resulting in a combined dataset from 847 breast cancer samples. Binary HRD labels for the bulk samples were derived using HRDetect, applying a threshold of 0.7. HRDetect integrates information from whole-genome-derived point mutations, indels, copy number alterations, and structural variants. Of the 847 samples, 112 were excluded by burden filtering (50 < indel count < 5,000 and SNV count < 20,000), yielding 735 retained samples. All 735 had an assigned HRDetect class label, of which 273 are HRDetect-high (positives) and 462 are HRDetect low/mid (negatives), giving a source-cohort positive prevalence of 0.371. Because HRDscout was designed for singlecell duplex data, we included only the variant types detectable by this assay and excluded structural-variant–based features. To train HRDscout across a range of mutation-detection sensitivities representative of single-cell whole-genome sequencing technologies — including those lower than scNanoSeq’s median of 2.5% — each retained sample was multinomially down-sampled at eight ratios (10×, 20×, 30×, 40×, 50×, 100×, 120×, 150×), with each ratio repeated two to three times, yielding a synthetic training matrix of 15,435 samples × 179 features (96 SBS96 + 83 ID83 channels) with a constant positive (HRD class) prevalence of 0.371 across all regimes. We retained the very-low-sensitivity regimes (120× and 150×) during training to include very low burden cases, although we restrict reported headline statistics to ratios ≤ 100× (simulated detection sensitivity ≥ 1%), comfortably bracketing the scNanoSeq sensitivity range (median 2.5%, IQR 1.9– 3.3%).

#### Imbalance-aware validation metrics

HRDscout was evaluated by stratified 10-fold cross-validation, with folds formed by random partitioning of the training matrix stratified on the HRDetect label; folds were not stratified by simulated detection sensitivity, and each fold’s training and heldout partitions span all down-sampling ratios. Per-sensitivity metrics were obtained by post-hoc stratification of the out-of-fold predictions. Across the six reported regimes (simulated detection sensitivity 1.0–10%), we report cross-validated precision-recall curves (Figures 6a, S20). We tabulate accuracy metrics for tumor data downsampled to scNanoSeq mutation detection sensitivity (assessed for 2.0–3.3% detection power; median 2.5%; compared to scNanoSeq’s median somatic mutation detection power of 2.5%, IQR: 1.9–3.3%). We report precision, recall, F1-score and Matthews correlation coefficient at the FPR ≤ 0.02 operating points used in the main analysis, alongside the threshold-independent area under the precision-recall curve (Supplementary Table S13).

To rule out leakage from downsampled replicates of the same tumor appearing in both the training and validation partitions, we re-ran the evaluation with grouped cross-validation by original tumor (and nested grouped CV for threshold selection); AUC, PR-AUC, F1, and MCC were essentially unchanged at every sensitivity (mean absolute change ≤ 0.007), retaining AUC ≥ 0.97 even at 100× downsampling.

#### Model interpretation

To interpret the model and quantify the relative contribution of each feature to the HRD prediction, we computed SHAP (SHapley Additive exPlanations) values. The SHAP framework enables feature attribution by measuring the marginal contribution of each input variable to the model’s predictions across samples.

The HRDscout classifier was subsequently applied to single-cell mutation profiles to assign HRD probabilities to individual cells that were profiled with scNanoSeq.

### Pooled indel signature analysis of non-tumor cells

To quantify HRD-associated indel patterns below the sensitivity of per-cell HRDscout classification, we estimated the fraction of long indels with microhomology in pooled cell populations. Because individual cells can sometimes contain too few somatic indels for stable estimation, indels were aggregated across predefined cell groups within each sample to obtain sufficient counts.

For HRD tumor samples, cells were stratified into (1) a tumor pool and (2) a non-tumor, non-HRD pool (HRDscout score < 0.7). For non-HRD tumors, (3) all cells were pooled to estimate the background rate of microhomology-associated indels. Within each pool (1-3), we computed the fraction of deletions ≥5 bp exhibiting microhomology, the indel class most strongly associated with HRD, as a per-sample summary of HRD-related indel activity. This metric was then compared across three groups using two-sided Mann–Whitney U tests (Figure S25).

### Annotation of pathogenic germline variants in the scNanoSeq cohort

Germline point mutations and indels were annotated with VEP using the Ensembl reference dataset (v.112, genome build GRCh37) and the ClinVar database (20241001). We identified pathogenic germline mutations in *BRCA1, BRCA2* and *PALB2* genes, and selected variants considered to be pathogenic in ClinVar’s ‘CLNSIG’ field.

### Visualization of lineage trees

Phylogenetic trees were visualized using the Phylo module from the Biopython package (v1.84).

### Subclonal and cell-private SBS3:SBS1 ratio analysis

To test the assumption that the SBS3:SBS1 rate is stable after HRD onset, we compared SBS3:SBS1 activity across successive evolutionary stages using two complementary approaches (Figure S8).

#### Bulk WGS

In whole-genome-duplicated regions of the GEL bulk WGS cohort (n = 56), mutations were partitioned into late-clonal and subclonal according to MutationTimeR maximum a posteriori timing classification. For each set, the SBS3:SBS1 ratio was estimated by multinomial resampling of the 96-context mutation profile followed by signature refitting (100 iterations; median and 95% CI), retaining samples with a median bootstrapped SBS1 exposure > 10. Late-clonal and subclonal ratios were strongly correlated (Spearman *ρ* = 0.72, P = 9.9 × 10^−9^, *n* = 47).

#### Single-cell scNanoSeq

Let n and DP denote, respectively, the number of cells supporting a variant (cells with ≥ 1 supporting duplex UMI) and the total duplex-UMI depth pooled across cells at that site. In 5 HRD samples sequenced with scNanoSeq (median duplex-UMI depth 33 across variant sites), variants were classified by duplex UMI support as private (n = 1, DP > 15), subclonal (2 ≤ n ≤ 10, DP >15), clonal (n > 10, DP > 15), or low-coverage (DP ≤ 15), and cell-private and subclonal SBS3:SBS1 ratios were compared within each sample (1,000 bootstrap iterations). Cell-private and subclonal ratios were concordant across all five samples, falling close to the diagonal and within the range observed in bulk, extending the stability seen between late-clonal and subclonal stages to the most recent, cell-private compartment.

#### Treatment of subclonal mutations

Subclonal mutations were far less abundant than clonal ones (mean per-patient subclonal fraction 14.1% GEL, 5.4% PCAWG, 1.7% SCANB, 8.8% INFORM), yet SBS3 was reliably detected in the GEL subclonal compartment. The core HRDTimer inference deliberately uses only clonal mutations, since subclonal detection in bulk scales with sample purity and would otherwise introduce a bias; the subclonal and cell-private periods are instead accounted for when translating HRD onset to calendar age (see “Estimation of SBS1 burden at diagnosis”).

### Sensitivity of HRD-onset estimates to SBS3-rate variation

To quantify the sensitivity of age-at-HRD estimates to the assumption of a constant post-HRD SBS3 rate, we treated each tumor’s HRDTimer solution (HRD and WGD timings on the SBS1-based molecular-time axis) as ground truth. We then redistributed SBS3 mutations across molecular time using alternative SBS3 rate functions while keeping all other tumor properties fixed, including copy-number state, total mutation burden, Early/Late/NA phase classification, total SBS3 burden, and the per-phase non-SBS3 signature mixture.

We evaluated four SBS3 rate functions: baseline (constant), ramp-up (gradual activation), oscillate (sinusoidal modulation), and oscillate-sat (square-wave bursts). Each function was normalized to the observed SBS3 burden, NSBS3, over the post-HRD interval [t_HRD_, 1] (Eq. 1; Supplementary Note 3). As a result, all scenarios had the same expected total number of SBS3 mutations and differed only in their distribution through molecular time.

For each sample–scenario pair, we generated 200 bootstrap replicates. Each replicate was processed through the full HRDTimer and age-translation pipeline. Signature labels and SBS96 channels were resampled according to the SBS3 probabilities specified by the corresponding rate function. The resulting age-at-HRD estimates were compared with those obtained under the constant-rate model.

We applied this framework to the n = 20 PCAWG breast cancer samples that passed all QC criteria and were included in the main analysis. The baseline, oscillate, and oscillate-sat scenarios produced estimates that were centered near zero bias. The ramp-up scenario was designed as a stress test that concentrates SBS3 mutations late in the post-HRD period, thereby increasing the Late-phase SBS3:SBS1 ratio. This scenario produced the largest systematic shift, with a mean bias of +3.8 years. Even so, this effect remained smaller than the HRDTimer’s age uncertainty for most samples.

Full specification is provided in Supplementary Note 3, “Sensitivity of age-at-HRD estimates to post-HRD SBS3 rate assumptions”.

### Statistics

All reported P-values are two-tailed. Group comparisons of HRD onset age, duration, and related continuous variables were performed using two-sided Mann–Whitney U tests. Where multiple comparisons were made, p-values were adjusted using the Benjamini–Hochberg false discovery rate procedure. Multivariable associations with SBS1 burden were assessed using ordinary least squares (OLS) regression; reported effect sizes and p-values for individual covariates are derived from two-sided t-tests on OLS coefficients (i.e. testing whether each coefficient differs from zero). An equivalent ridge regression model with a cross-validated L2 penalty was fitted in parallel to account for collinearity among covariates; bootstrap 95% confidence intervals were estimated from 500 resampling iterations.

## Data and Code Availability

The somatic mutation calls for whole genome samples with HRD and WGD (PCAWG, SCANB, INFORM) are available as Chromoscope configurations.

- [PCAWG HRD WGD cohort](https://chromoscope.bio/app/?external=https://somatic-browser-test.s3.amazonaws.com/hrdtimer_PCAWG/configs/hrdtimer_PCAWG.all.config.json)
- [SCANB HRD WGD cohort](https://chromoscope.bio/app/?external=https://somatic-browser-test.s3.amazonaws.com/hrdtimer_SCANB/configs/hrdtimer_SCANB.config.json)
- [INFORM HRD WGD cohort](https://chromoscope.bio/app/?external=https://somatic-browser-test.s3.us-east-1.amazonaws.com/inform/INFORM_config_hrdtiming.json)

Data from the National Genomic Research Library (NGRL) used in this research are available within the secure Genomics England Research Environment. Access to NGRL data is restricted to adhere to consent requirements and protect participant privacy. Data used in this research include: point mutation calls and copy number profiles. VCF files of somatic mutations, annotated with allele-specific copy number and MutationTimeR timing, are available in the Research Environment at the following folder:

/re_gecip/cancer_pan/mandreopoulos/2026_Andreopoulos_GEL_Mutation-TimeR_WGD_HRD_Breast_vcfs/

Access to NGRL data is provided to approved researchers who are members of the Genomics England Research Network, subject to institutional access agreements and research project approval under participant-led governance. For more information on data access, visit: https://www.genomicsengland.co.uk/research.

The raw sequencing reads from the scNanoSeq experiments are downloaded from the NCBI Sequence Read Archive (SRA) under the accession codes listed in the original publication (PRJNA958124 and PRJNA962101).^22^ Code for HRDTimer and the analysis of bulk whole genome samples is available through https://github.com/michandreop/HRDTimer/.

## Acknowledgments

- D.G. was awarded and supported by the ‘Developmental Research Project Program’ from Dana Farber/Harvard Cancer Center SPORE in Breast Cancer grant number 1P50CA168504-08.
- M.A. and D.G. are supported by the ‘major grant’ award from the Fund for Innovation in Cancer Informatics awarded to D.G. and P.P.
- NIH SMaHT grant to Data Analysis Center at Harvard Medical School (awarded to P.P.) and Genome Characterization Center at Baylor College of Medicine (C.Z.).
- We thank:
  ∘ Aliah Hawari, Avraam Tapinos, and David C. Wedge for their help with the Battenberg analysis of GEL data.
  ∘ Shannon Ehmsen for her help with figure design.
  ∘ Teng Gao for help with running his tool ‘Numbat’
  ∘ Arezou Ghazani for help with assessing mutation pathogenicity.

*We gratefully acknowledge the participants of the National Genomic Research Library (NGRL), whose contributions made this research possible. Secure access to the NGRL under project ID 1449 was provided by Genomics England, which delivers the NGRL in partnership with NHS England, and is wholly owned by the UK Department of Health and Social Care. The NGRL contains participants’ health data collected by the NHS as part of their care, along with samples and data from their participation in research, for which fully informed consent has been obtained. This includes genomic and clinical data provided through the NHS Genomic Medicine Service, as well as data obtained through research studies, including the 100,000 Genomes Project and the Generation Study, both of which are delivered in partnership with the NHS, and from other research cohorts involving external collaborators*.

## Author Contributions

- Conceptualization: D.G. (Dominik Glodzik), M.A., Doga Gulhan, C.Z.
- Data curation: D.G., M.A., M.N., Y.Z.
- Formal analysis: M.A. and D.G.
- Funding acquisition: D.G. and P.P.
- Software: M.A. and V.V.
- Supervision: D.G., P.P., C.Z.
- Visualization: M.A. and D.G.
- Writing - original draft: D.G. and M.A.
- Writing - review and editing: D.G., M.A., P.P.

## Competing Interests

The authors declare no competing interests.

## Usage of generative AI for editing

During the preparation of this work the authors used ChatGPT in order to streamline phrasing. After using this tool/service, the authors reviewed and edited the content as needed and take full responsibility for the content of the publication.

## Notes

### Competing Interest Statement

The authors have declared no competing interest.

### Summary of Updates

Cohort expanded from 59 to 115 breast cancers by adding 56 Genomics England tumors to PCAWG, SCANB and INFORM. All analyses were rerun on the expanded cohort: HRD and WGD timing, SBS1 burden and acceleration estimates, subtype and cohort comparisons, and the associated statistics. Conclusions remain largely unchanged, with updated values throughout. Abstract, Results, Discussion, figures and supplementary material updated accordingly.

## References

1. Kuchenbaecker, K. B., Hopper, J. L., Barnes, D. R., Phillips, K.-A., Mooij, T. M., Roos-Blom, M.-J., Jervis, S., Van Leeuwen, F. E., Milne, R. L., Andrieu, N., et al. (2017). Risks of breast, ovarian, and contralateral breast cancer for BRCA1 and BRCA2 mutation carriers. Jama 317, 2402–2416.

2. Antoniou, A., Cunningham, A., Peto, J., Evans, D. G., Lalloo, F., Narod, S. A., Risch, H. A., Eyfjord, J. E., Hopper, J. L., Southey, M. C., et al. (2008). The BOADICEA model of genetic susceptibility to breast and ovarian cancers: updates and extensions. British Journal of Cancer 98, 1457–1466.

3. National Comprehensive Cancer Network (2025). Genetic/Familial High-Risk Assessment: Breast, Ovarian, Pancreatic, and Prostate. Version 3.2025.

4. Brose, M. S., Rebbeck, T. R., Calzone, K. A., Stopfer, J. E., Nathanson, K. L., and Weber, B. L. (2002). Cancer risk estimates for BRCA1 mutation carriers identified in a risk evaluation program. Journal of the National Cancer Institute 94, 1365–1372.

5. Davies, H., Glodzik, D., Morganella, S., Yates, L. R., Staaf, J., Zou, X., Ramakrishna, M., Martin, S., Boyault, S., Sieuwerts, A. M., et al. (2017). HRDetect is a predictor of BRCA1 and BRCA2 deficiency based on mutational signatures. Nature medicine 23, 517–525.

6. Pathania, S., Bade, S., Le Guillou, M., Burke, K., Reed, R., Bowman-Colin, C., Su, Y., Ting, D. T., Polyak, K., Richardson, A. L., et al. (2014). BRCA1 haploinsufficiency for replication stress suppression in primary cells. Nature communications 5, 5496.

7. Jonsson, P., Bandlamudi, C., Cheng, M. L., Srinivasan, P., Chavan, S. S., Friedman, N. D., Rosen, E. Y., Richards, A. L., Bouvier, N., Selcuklu, S. D., et al. (2019). Tumour lineage shapes BRCA-mediated phenotypes. Nature 571, 576–579.

8. Karaayvaz-Yildirim, M., Silberman, R. E., Langenbucher, A., Saladi, S. V., Ross, K. N., Zarcaro, E., Desmond, A., Yildirim, M., Vivekanandan, V., Ravichandran, H., et al. (2020). Aneuploidy and a deregulated DNA damage response suggest haploinsufficiency in breast tissues of BRCA2 mutation carriers. Science Advances 6, eaay2611.

9. Kong, L. R., Gupta, K., Wu, A. J., Perera, D., Ivanyi-Nagy, R., Ahmed, S. M., Tan, T. Z., Tan, S. L.-W., Fuddin, A., Sundaramoorthy, E., et al. (2024). A glycolytic metabolite bypasses “two-hit” tumor suppression by BRCA2. Cell 187, 2269–2287.

10. Geyer, F. C., Pareja, F., Weigelt, B., Rakha, E., Ellis, I. O., Schnitt, S. J., and Reis-Filho, J. S. (2017). The spectrum of triple-negative breast disease: high-and low-grade lesions. The American journal of pathology 187, 2139–2151.

11. Maclnnes, E., Duffy, S., Simpson, J., Wallis, M., Turnbull, A., Wilkinson, L., et al. (2020). Radiological audit of interval breast cancers. Estimation of tumour growth rates Breast 51, 114–119.

12. Nee, K., Ma, D., Nguyen, Q. H., Pein, M., Pervolarakis, N., Insua-Rodríguez, J., Gong, Y., Hernandez, G., Alshetaiwi, H., Williams, J., et al. (2023). Preneoplastic stromal cells promote BRCA1-mediated breast tumorigenesis. Nature genetics 55, 595–606.

13. Williams, M. J., Oliphant, M. U., Au, V., Liu, C., Baril, C., O’Flanagan, C., Lai, D., Beatty, S., Van Vliet, M., Yiu, J. C., et al. (2024). Luminal breast epithelial cells of BRCA1 or BRCA2 mutation carriers and noncarriers harbor common breast cancer copy number alterations. Nature Genetics, 1–10.

14. Jolly, C. and Van Loo, P. (2018). Timing somatic events in the evolution of cancer. Genome biology 19, 1–9.

15. Pan-cancer analysis of whole genomes (2020). Nature 578, 82–93.

16. Staaf, J., Glodzik, D., Bosch, A., Vallon-Christersson, J., Reuterswärd, C., Häkkinen, J., Degasperi, A., Amarante, T. D., Saal, L. H., Hegardt, C., et al. (2019). Whole-genome sequencing of triple-negative breast cancers in a population-based clinical study. Nature medicine 25, 1526–1533.

17. Tung, N., Arun, B., Hacker, M. R., Hofstatter, E., Toppmeyer, D. L., Isakoff, S. J., Borges, V., Legare, R. D., Isaacs, C., Wolff, A. C., et al. (2020). TBCRC 031: randomized phase II study of neoadjuvant cisplatin versus doxorubicin-cyclophosphamide in germline BRCA carriers with HER2-negative breast cancer (the INFORM trial). Journal of Clinical Oncology 38, 1539–1548.

18. Genomics England (2024). The National Genomic Research Library. Dataset. 10.6084/m9.figshare.4530893.

19. Black, D., Davies, H. R., Koh, G. C. C., Chmelova, L., Cubric, M., Chan, G. C., Degasperi, A., Czarnecki, J., Toong, P. J., Memari, Y., et al. (2025). Clinical potential of whole-genome data linked to mortality statistics in patients with breast cancer in the UK: a retrospective analysis. The Lancet Oncology 26, 1417–1431.

20. Nishimura, T., Kakiuchi, N., Yoshida, K., Sakurai, T., Kataoka, T. R., Kondoh, E., Chigusa, Y., Kawai, M., Sawada, M., Inoue, T., et al. (2023). Evolutionary histories of breast cancer and related clones. Nature 620, 607–614.

21. Yates, L. R., Knappskog, S., Wedge, D., Farmery, J. H., Gonzalez, S., Martincorena, I., Alexandrov, L. B., Van Loo, P., Haugland, H. K., Lilleng, P. K., et al. (2017). Genomic evolution of breast cancer metastasis and relapse. Cancer cell 32, 169–184.

22. Niu, M., Zhang, Y., Luo, J., Sinson, J. C., Thompson, A. M., and Zong, C. (2023). Characterization of cancer evolution landscape based on accurate detection of somatic mutations in single tumor cells. bioRxiv, 2023–10.

23. Institute, W. S. (2020). COSMIC Mutational Signatures. https://cancer.sanger.ac.uk/signatures/.

24. Alexandrov, L. B., Jones, P. H., Wedge, D. C., Sale, J. E., Campbell, P. J., Nik-Zainal, S., and Stratton, M. R. (2015). Clock-like mutational processes in human somatic cells. Nature genetics 47, 1402–1407.

25. Spisak, N. Manuel, M. de, Milligan, W., Sella, G., and Przeworski, M. (2024). The clock-like accumulation of germline and somatic mutations can arise from the interplay of DNA damage and repair. PLoS biology 22, e3002678.

26. Alexandrov, L. B., Ju, Y. S., Haase, K., Van Loo, P., Martincorena, I., Nik-Zainal, S., Totoki, Y., Fujimoto, A., Nakagawa, H., Shibata, T., et al. (2016). Mutational signatures associated with tobacco smoking in human cancer. Science 354, 618–622.

27. Gerstung, M., Jolly, C., Leshchiner, I., Dentro, S. C., Gonzalez, S., Rosebrock, D., Mitchell, T. J., Rubanova, Y., Anur, P., Yu, K., et al. (2020). The evolutionary history of 2,658 cancers. Nature 578, 122–128.

28. Jin, H., Gulhan, D. C., Geiger, B., Ben-Isvy, D., Geng, D., Ljungström, V., and Park, P. J. (2024). Accurate and sensitive mutational signature analysis with MuSiCal. Nature Genetics 56, 541–552.

29. Martínez-Jiménez, F., Priestley, P., Shale, C., Baber, J., Rozemuller, E., and Cuppen, E. (2023a). Genetic immune escape landscape in primary and metastatic cancer. Nature Genetics 55, 820–831.

30. Telli, M. L., Timms, K. M., Reid, J., Hennessy, B., Mills, G. B., Jensen, K. C., Szallasi, Z., Barry, W. T., Winer, E. P., Tung, N. M., et al. (2016). Homologous recombination deficiency (HRD) score predicts response to platinum-containing neoadjuvant chemotherapy in patients with triple-negative breast cancer. Clinical cancer research 22, 3764–3773.

31. Tomkova, M., McClellan, M. J., Crevel, G., Shahid, A. M., Mozumdar, N., Tomek, J., Shepherd, E., Cotterill, S., Schuster-Böckler, B., and Kriaucionis, S. (2024). Human DNA polymerase ε is a source of C> T mutations at CpG dinucleotides. Nature Genetics, 1–11.

32. Kozlov, A. M., Alves, J. M., Stamatakis, A., and Posada, D. (2022). CellPhy: accurate and fast probabilistic inference of single-cell phylogenies from scDNA-seq data. Genome Biology 23, 37.

33. Coorens, T. H., Oh, J. W., Choi, Y. A., Lim, N. S., Zhao, B., Voshall, A., Abyzov, A., Antonacci-Fulton, L., Aparicio, S., Ardlie, K. G., et al. (2025). The somatic mosaicism across human tissues network. Nature 643, 47–59.

34. Tutt, A. N., Garber, J. E., Kaufman, B., Viale, G., Fumagalli, D., Rastogi, P., Gelber, R. D., De Azambuja, E., Fielding, A., Balmaña, J., et al. (2021). Adjuvant olaparib for patients with BRCA1-or BRCA2-mutated breast cancer. New England Journal of Medicine 384, 2394–2405.

35. Nik-Zainal, S., Davies, H., Staaf, J., Ramakrishna, M., Glodzik, D., Zou, X., Martincorena, I., Alexandrov, L. B., Martin, S., Wedge, D. C., et al. (2016). Landscape of somatic mutations in 560 breast cancer whole-genome sequences. Nature 534, 47–54.

36. ICGC Data Coordination Center (n.d.). ICGC Data Portal. https://dcc.icgc.org/.

37. Martínez-Jiménez, F., Movasati, A., Brunner, S. R., Nguyen, L., Priestley, P., Cuppen, E., and Van Hoeck, A. (2023b). Pan-cancer whole-genome comparison of primary and metastatic solid tumours. Nature 618, 333–341.

38. Shale, C., Cameron, D. L., Baber, J., Wong, M., Cowley, M. J., Papenfuss, A. T., Cuppen, E., and Priestley, P. (2022). Unscrambling cancer genomes via integrated analysis of structural variation and copy number. Cell Genomics 2.

39. Degasperi, A., Amarante, T. D., Czarnecki, J., Shooter, S., Zou, X., Glodzik, D., Morganella, S., Nanda, A. S., Badja, C., Koh, G., et al. (2020). A practical framework and online tool for mutational signature analyses show intertissue variation and driver dependencies. Nature cancer 1, 249–263.

40. Nguyen, L., WM Martens, J., Van Hoeck, A., and Cuppen, E. (2020). Pan-cancer landscape of homologous recombination deficiency. Nature communications 11, 5584.

41. ICGC ARGO Data Portal (n.d.). Accessing ICGC 25K Release Data. https://docs.icgc-argo.org/docs/data-access/daco/applying.

42. Nik-Zainal, S. (2020). Whole-genome-sequencing of triple negative breast cancers: a population study. Version 3. 10.17632/2mn4ctdpxp.3.

43. Freed, D., Pan, R., and Aldana, R. (2018). TNscope: accurate detection of somatic mutations with haplotype-based variant candidate detection and machine learning filtering. biorxiv, 250647.

44. Campbell, P. (2017). Genomic evolution of breast cancer metastasis and relapse - Yates et al. Version V1. Mendeley Data. 10.17632/g7kpzkhz8c.1.

45. Seplyarskiy, V. B., Soldatov, R. A., Koch, E., McGinty, R. J., Goldmann, J. M., Hernandez, R. D., Barnes, K., Correa, A., Burchard, E. G., Ellinor, P. T., et al. (2021). Population sequencing data reveal a compendium of mutational processes in the human germ line. Science 373, 1030–1035.

46. Morganella, S., Alexandrov, L. B., Glodzik, D., Zou, X., Davies, H., Staaf, J., Sieuwerts, A. M., Brinkman, A. B., Martin, S., Ramakrishna, M., et al. (2016). The topography of mutational processes in breast cancer genomes. Nature communications 7, 11383.

47. Gerstung Lab (2024). MutationTimeR. GitHub repository.

48. Nakashima, K., Uematsu, T., Takahashi, K., Nishimura, S., Tadokoro, Y., Hayashi, T., and Sugino, T. (2019). Does breast cancer growth rate really depend on tumor subtype? Measurement of tumor doubling time using serial ultrasonography between diagnosis and surgery. Breast Cancer 26, 206–214.

49. Nik-Zainal, S., Van Loo, P., Wedge, D. C., Alexandrov, L. B., Greenman, C. D., Lau, K. W., Raine, K., Jones, D., Marshall, J., Ramakrishna, M., et al. (2012). The life history of 21 breast cancers. Cell 149, 994–1007.

