## Supplementary material for "Timing the onset of homologous recombination deficiency before breast cancer diagnosis": 00_HRDTimer_Supplementary_Information.pdf

### Supplementary Information
### for "Timing the onset of homologous recombination deficiency before cancer diagnosis"

Michail Andreopoulos[1], Muchun Niu[2,3], Yang Zhang[2,4], Vinayak V. Viswanadham[1], Doga C. Gulhan[1], Hu Jin[1], Felipe Batalini[5], Gerburg Wulf [6], Chenghang Zong[1,7,8,*], Peter J. Park[1,*], and Dominik Glodzik[1*]

### Supplemental Figures

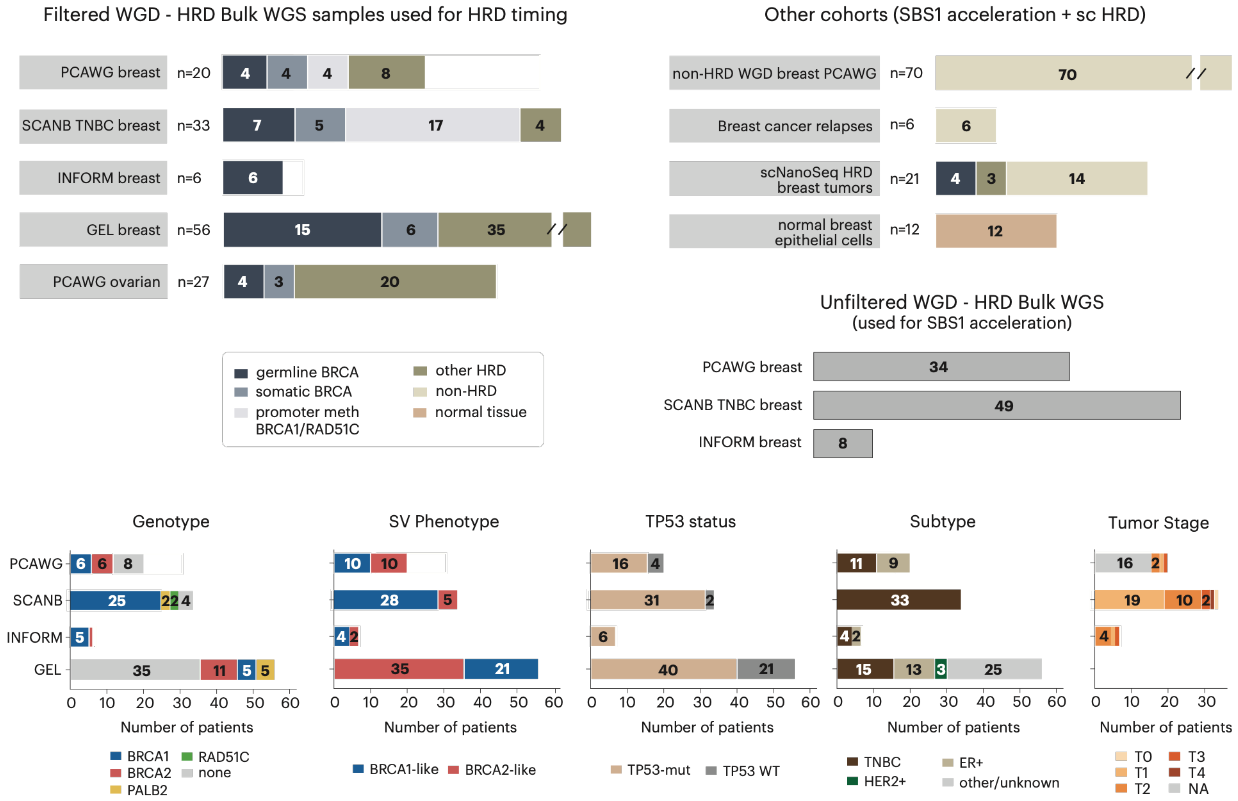


**Figure S1. Cohort diagram.** Sample counts across cohorts profiled by whole-genome sequencing or single-cell Nano-seq (scNanoSeq). HRD+WGD samples are stratified by HRD gene genotype, structural variant (SV) phenotype (see Figure S19), *TP53* mutation status, receptor subtype, and tumor stage.

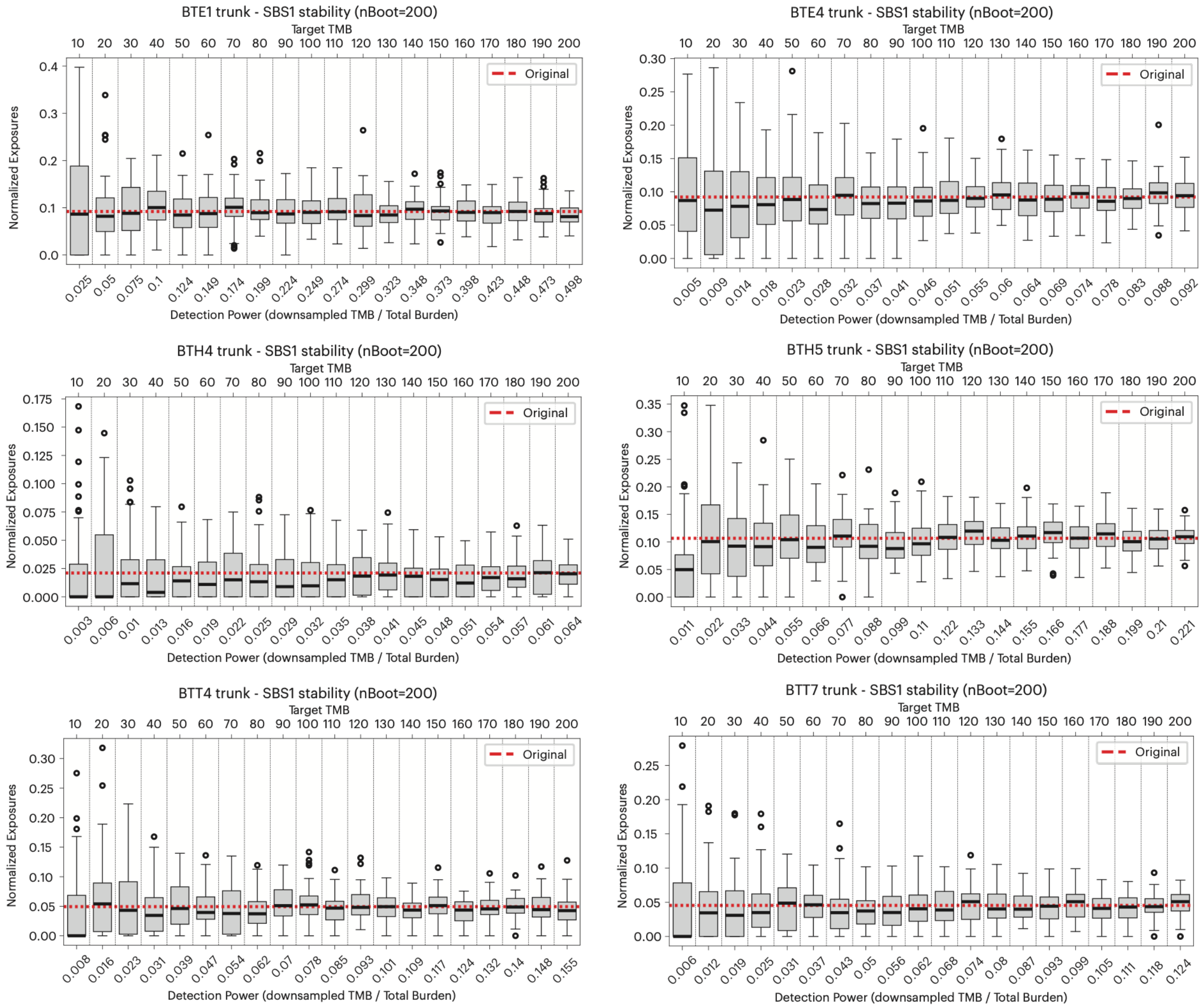


**Figure S2. Accurate recovery of the clock-like SBS1 signature at low mutation burdens common in scNanoSeq cells.** Truncal mutations from six samples (BTE1, BTE4, BTH4, BTH5, BTT4, BTT7) were randomly downsampled to a range of target mutation burdens (TMB = 10–200; top axis), with 200 replicates per target (nBoot = 200), and mutational signatures were re-fitted to each. The bottom axis shows the corresponding detection power, i.e. the target burden as a fraction of that sample's total truncal burden. Gray boxplots show normalized SBS1 exposures across replicates. Estimates remained centered on the original full-burden value (red dashed line) while dispersion decreased with increasing burden, indicating that SBS1 is recovered without systematic bias, and with improving precision, even under sparse single-cell conditions.

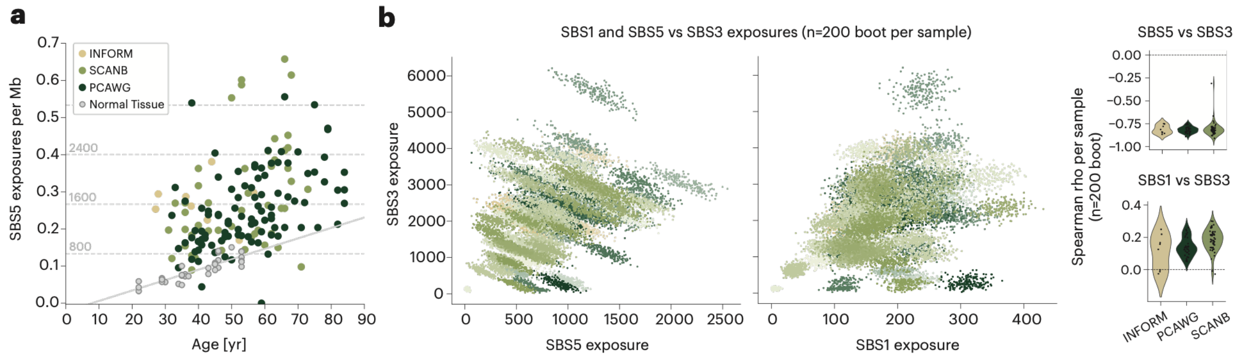


**Figure S3. Evaluation of SBS5 as an alternative molecular clock.** (**a**) SBS5 exposure per Mb versus patient age in WGD-positive breast cancers (INFORM, SCANB, PCAWG) and normal mammary epithelial organoids. Most tumor samples exceed the SBS5 burden expected from age-matched normal tissue (gray line), indicating that SBS5 — like SBS1 — undergoes acceleration during tumor evolution. A subset of samples shows SBS5 burdens comparable to normal tissue. Horizontal dashed lines indicate SBS5 burden rescaled to a diploid genome. (**b**) Joint behavior of SBS3, SBS5 and SBS1 exposures across bootstrap resamples of the mutation catalog (n = 200 bootstraps per sample). Left: SBS3 versus SBS5 exposures, showing strong intra-sample anti-correlation. Middle: SBS3 versus SBS1 exposures, showing largely independent estimation. Right: per-sample distributions of Spearman $\rho$ between SBS3 and SBS5 (top) and between SBS3 and SBS1 (bottom), across bootstraps. SBS3–SBS5 correlations are consistently strongly negative (median $\rho \approx -0.8$ across cohorts), whereas SBS3–SBS1 correlations are weakly positive and close to zero, indicating that SBS5, unlike SBS1, is not a reliable molecular clock for the HRDTimer algorithm.

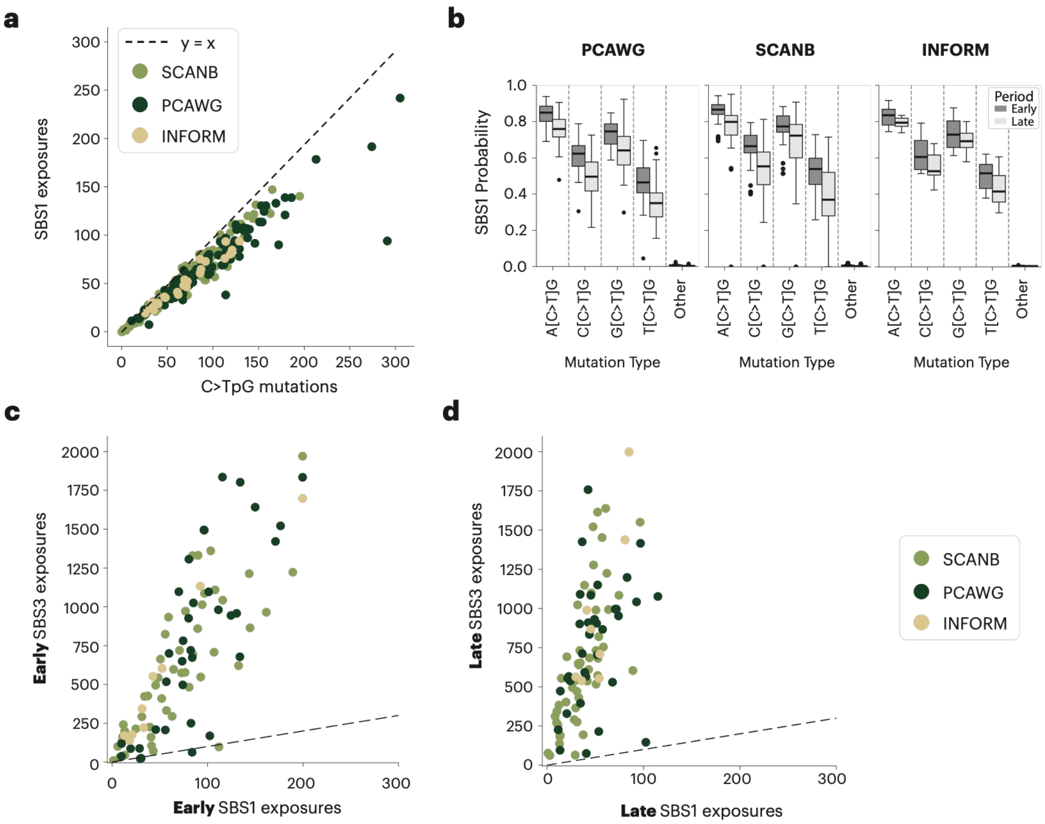


**Figure S4. Comparison of early and late SBS1 and SBS3 exposures.** (**a**) Comparison of [C>T]pG mutation counts and SBS1 exposures (estimated with MuSiCal) in HRD breast cancers. The excess of [C>T]pG relative to those attributed to SBS1 illustrates how non-clock-like processes can contribute [C>T]pG mutations, and motivates the use of signature fitting to estimate and refine the molecular clock. (**b**) Distribution of per-mutation SBS1 probabilities in early and late clonal mutations. Systematic differences between the early and late groups, along with variation across N[C>T]G trinucleotide contexts, motivate using SBS1 and fitting signatures separately for the early and late groups. (**c**) Estimated early clonal SBS1 and SBS3 mutation counts per sample. (**d**) Estimated late clonal SBS1 and SBS3 mutation counts per sample. The correlation between SBS1 and SBS3 counts within each group is comparable pre- and post-WGD across the cohort, suggesting a stable relative activity of these two processes

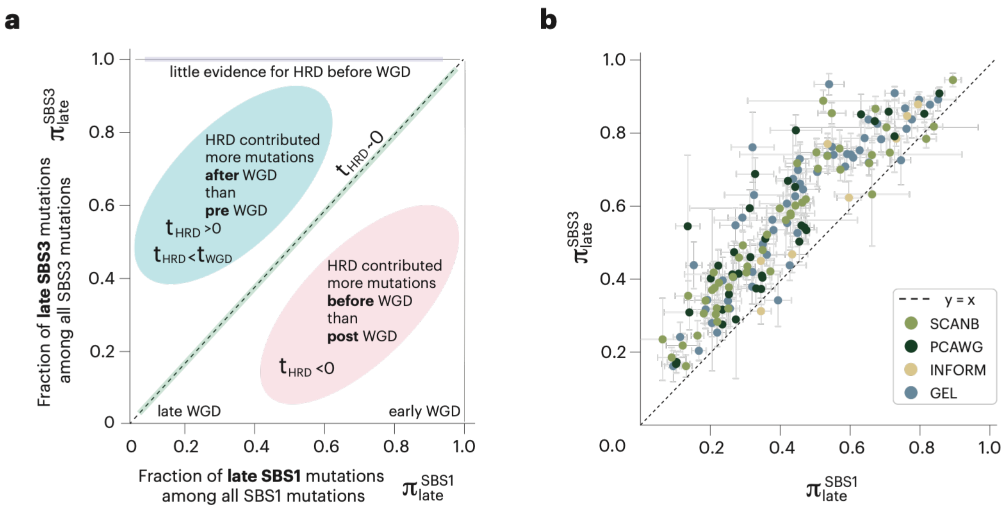


**Figure S5. Diagram explaining the space of signature-centered (SBS1, SBS3) fractions of late mutations.** Fraction of late mutations, attributed to SBS1 and SBS3, denoted by $\pi_{late}^{SBS1}$ and $\pi_{late}^{SBS3}$ across the cohort. Left: interpretation of points in areas of the plot. Right: observed data.

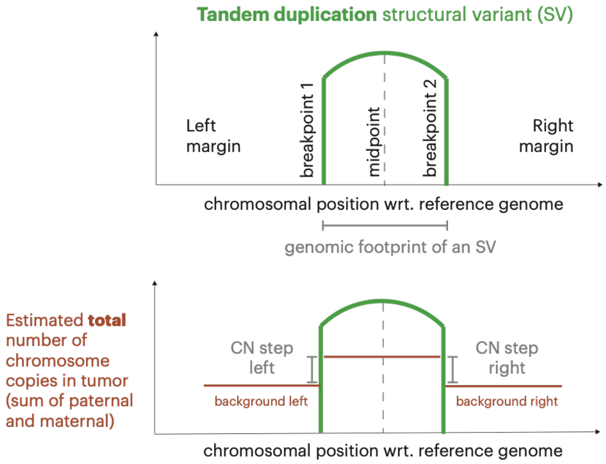


**Figure S6. Diagram of the local features used to time structural variants (SVs) with respect to genome duplication.** The “copy number (CN) step” at the breakpoints of an SV is quantified from the total copy number profile, using results from the PURPLE algorithm. SVs preceding genome duplication will have a CN step of 2, whereas late SVs will have CN step of 1. For this analysis, we used SVs where the wider region was consistent with a chromosomal gain, and where the copy number profile around the structural variant was symmetric and thus consistent with a singleton SV changing the local copy number in line with its type (deletion or tandem duplication, see Methods).

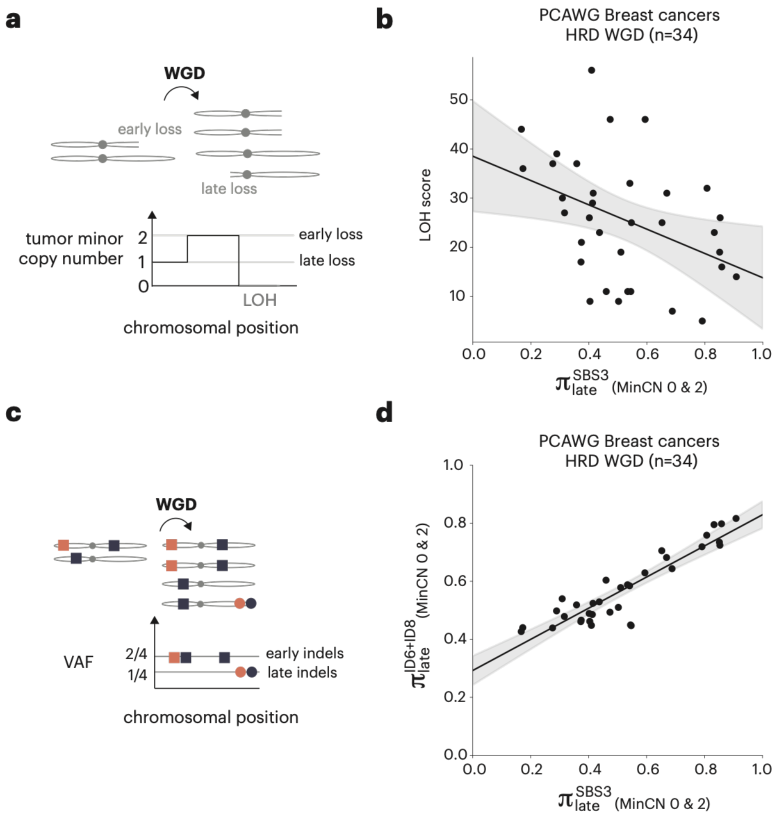


**Figure S7. Timing of other types of variants associated with HRD.** Correlations between the proportion of late SBS3 point mutations and HRD-associated LOH events and indels. (**a**) Early (pre-WGD) and late (post-WGD) point mutations were distinguished based on their allele fractions. Similarly, arm-level chromosomal losses were timed based on the magnitude of copy number change across breakpoints. (**b**) Scatterplot showing the relationship between $\pi_{late}^{SBS3}$ and HRD score in PCAWG breast cancer samples with both HRD and WGD (n=34). A negative correlation is expected and observed: early chromosomal arm losses typically result in loss of heterozygosity (LOH) because only a single copy of each parental chromosome is present, whereas arm losses post genome duplication are less likely to lead to LOH because each parental chromosome is present in two copies. (**c**) Schematic diagram illustrating how HRD-attributed indels were timed according to variant allele fraction. (**d**) Scatterplot showing the relationship between $\pi_{late}^{SBS3}$ and $\pi_{late}^{ID6+ID8}$ in PCAWG breast cancer samples with both HRD and WGD (n=34). $\pi_{late}$ values represent the proportion of late mutations, calculated in genomic regions where the major copy number is 2 and the minor copy number is 0 or 2—copy number states that allow for retrospective discrimination of variant timing relative to WGD.

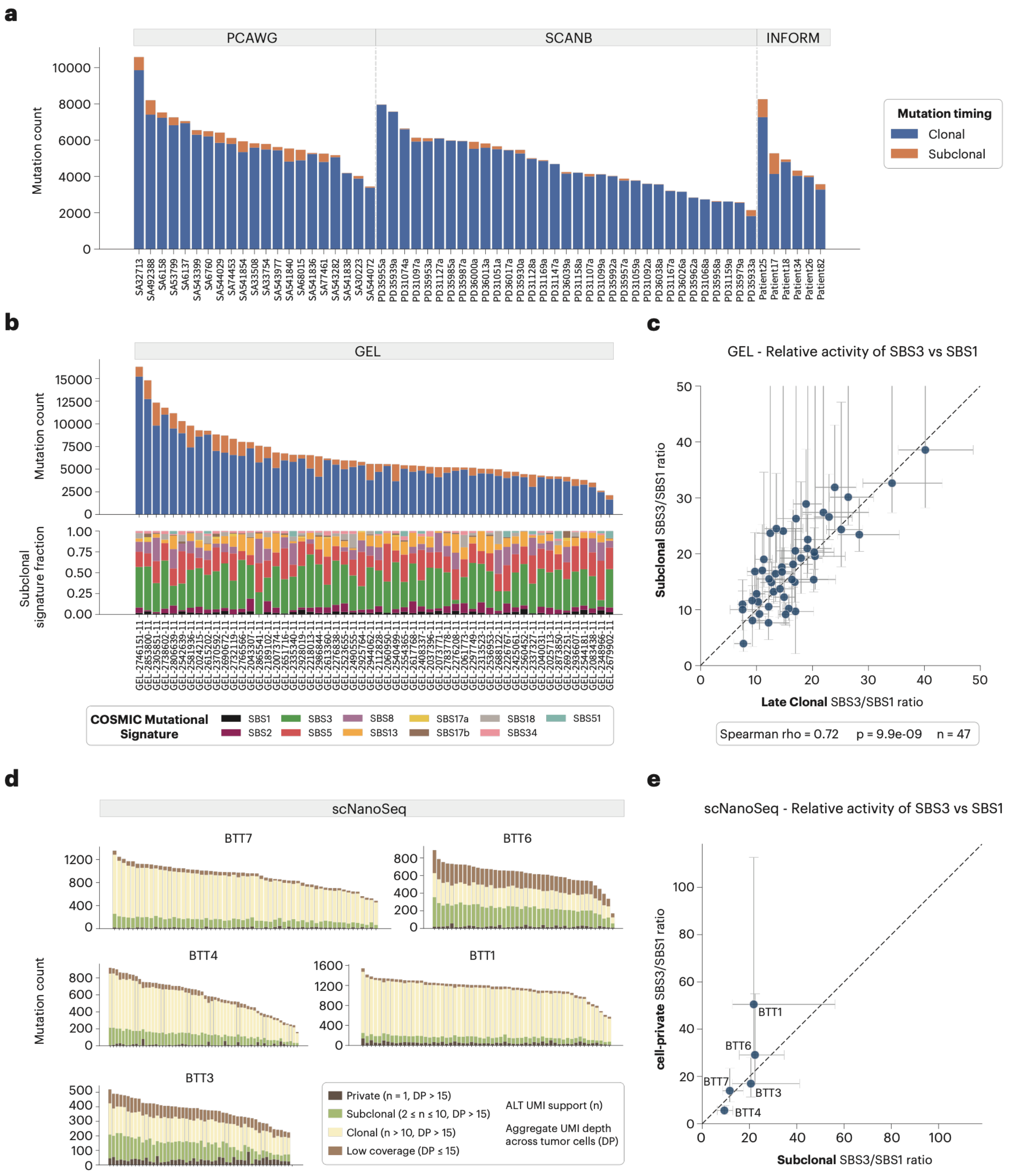


**Figure S8. Comparison of clonal/late and subclonal SBS3/SBS1 activity across bulk and single-cell cohorts.** (**a**) Per-tumor SNV burden in WGD regions stratified by mutation timing (clonal, blue; subclonal, orange) across PCAWG (n = 21), SCANB (n = 33), and INFORM (n = 6) HRD breast tumors, ordered by total count. One PCAWG sample has no recorded age and is excluded from the age-based timing analysis. Timing assigned from variant allele frequency and copy number state using MutationTimeR (MAP estimate). (**b**) Top: per-tumor clonal/subclonal SNV burden in WGD regions for GEL WGD-HRD breast tumors (n = 56). Bottom: COSMIC SBS signature contributions to the subclonal mutations (colors as defined in the panel). (**c**) Subclonal versus late clonal SBS3/SBS1 activity ratios for GEL Breast WGD-HRD tumors; points and bars, median and 95% bootstrap CI obtained by multinomial resampling of the SBS96 mutation profile followed by signature refitting (n = 100 bootstrap iterations); dashed line, $y = x$. Only samples with median SBS1 exposure count $> 10$ across bootstraps were retained. Spearman $\rho = 0.72$, $\mathrm{P} = 9.9 \times 10^{-9}, n = 47$. Large CIs are a result of low SBS1 counts. (**d**) Per-cell SNV counts in five scNanoSeq HRD breast tumors (BTT1, BTT3, BTT4, BTT6, BTT7), stratified by clonality. Variants were classified using duplex UMI counts by the number of tumor cells carrying $\geq 1$ ALT duplex UMI ($n$) and aggregate duplex UMI depth across tumor cells ($\mathrm{DP}$): Private ($n = 1$, $\mathrm{DP} > 15$), Subclonal ($2 \leq n \leq 10$, $\mathrm{DP} > 15$), Clonal ($n > 10$, $\mathrm{DP} > 15$), Low coverage ($\mathrm{DP} \leq 15$). Cells ordered by total count. (**e**) Cell-private versus subclonal SBS3/SBS1 ratios per scNanoSeq sample using the private and subclonal classes defined in (d); points and bars, point estimate and 95% bootstrap CI from multinomial resampling of the SBS96 spectrum (1,000 iterations); dashed line, $y = x$.

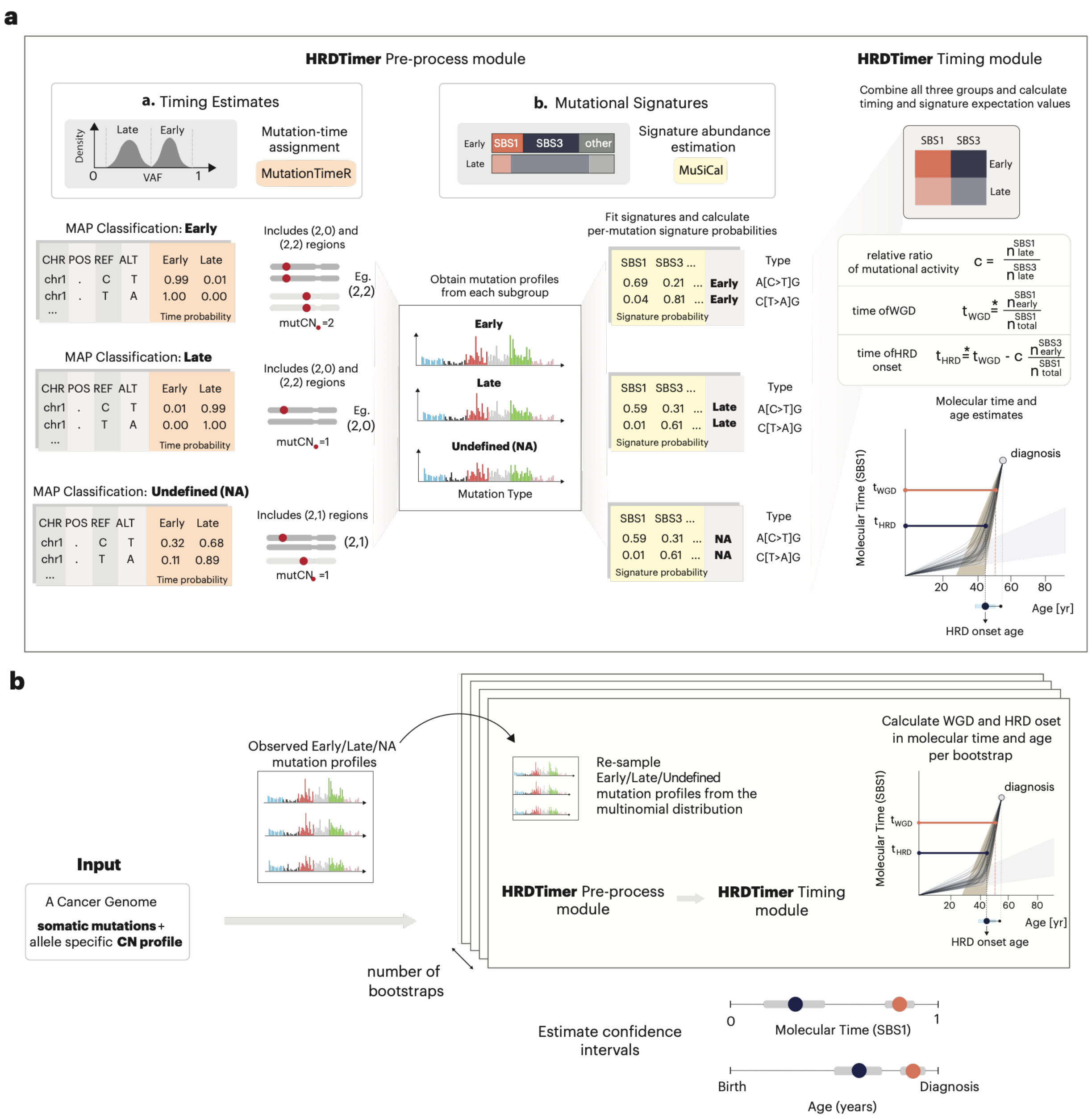


**Figure S9. Schematic representation of the key steps in the HRDTimer workflow.** (**a**) The main steps of a single iteration of the HRDTimer workflow for one sample. First, in the pre-processing module, individual mutations are assigned timing (Early, Late, or Undefined/NA) via MAP classification using the MutationTimeR algorithm and signature-of-origin probabilities using MuSiCal. These are then used collectively in the timing module to estimate the time points of WGD and HRD onset in SBS1-based molecular time. Finally, the calculated molecular time is converted to patient age using the relevant SBS1–age model. (**b**) The sampling scheme used to estimate confidence intervals for WGD and HRD onset, expressed in SBS1-based molecular time or patient age. Each iteration of the workflow steps shown in (a)—excluding the per-mutation timing estimates, which are computed once (See Methods)—is repeated across multiple bootstraps using mutational profiles resampled from a multinomial distribution parametrized by the observed mutation spectrum. Uncertainty in timing estimates is propagated from the uncertainty in the mutational signature exposures.

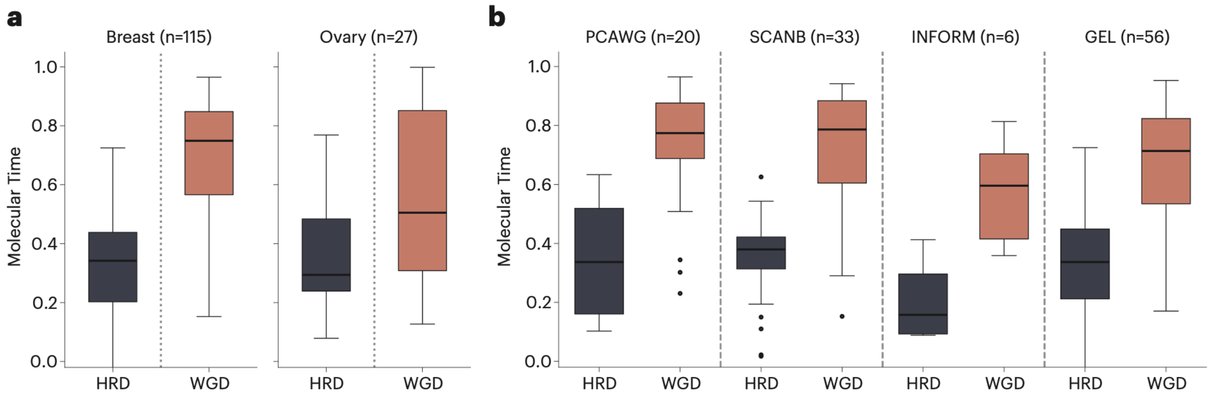


**Figure S10. Summary of molecular time estimates for HRD onset and whole-genome duplication (WGD).** Distributions of estimated HRD onset and WGD timing, expressed in SBS1-based molecular time, across samples. Results are stratified by tissue (**a**; breast, n = 115; ovary, n = 27) and by breast cancer cohort (**b**; PCAWG, n = 20; SCANB, n = 33; INFORM, n = 6; GEL, n = 56). In each group, HRD onset precedes WGD. Boxplots show the median, interquartile range, and whiskers; points denote outliers.

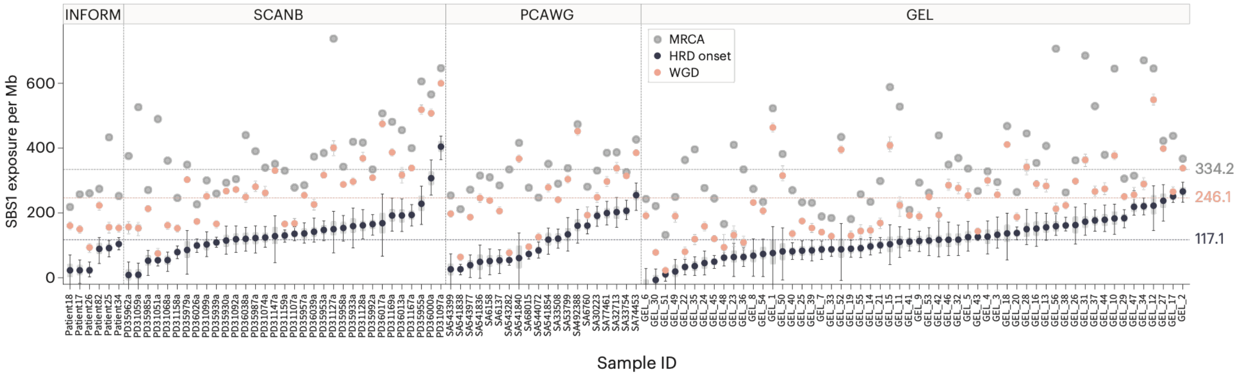


**Figure S11. SBS1 exposures per Mb at whole-genome duplication and HRD onset relative to the MRCA in breast cancer cohorts.** SBS1 exposures per Mb were calculated as molecular time multiplied by genome-corrected SBS1 counts, normalized to the effective genome size (G). Samples are shown individually, grouped by cohort (INFORM, SCANB, PCAWG, GEL). Gray dots represent total SBS1 exposure at the most recent common ancestor (MRCA); colored points represent estimated exposure at WGD (peach) and HRD onset (navy). Error bars indicate 95% confidence intervals, and shaded gray bands show interquartile ranges across samples. Dashed lines denote the median exposure across samples for each event type (labeled at right: MRCA, WGD, and HRD onset).

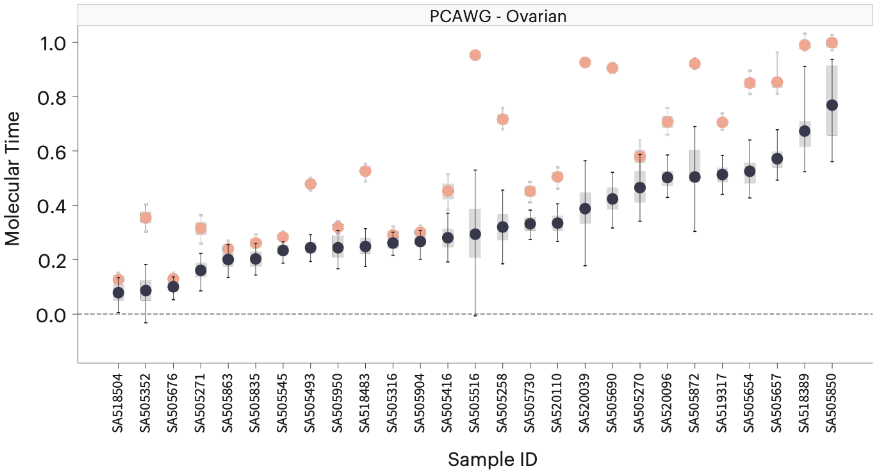


**Figure S12. Estimates of molecular timing for HRD onset and whole-genome duplication in PCAWG ovarian cancers.** Per-sample estimates of HRD onset (navy) and WGD (peach) timing in SBS1-based molecular time (n = 27). Error bars denote 95% confidence intervals and shaded gray bands denote the interquartile range, both estimated by bootstrapping. Samples are sorted by the estimated SBS1 molecular time of HRD onset.

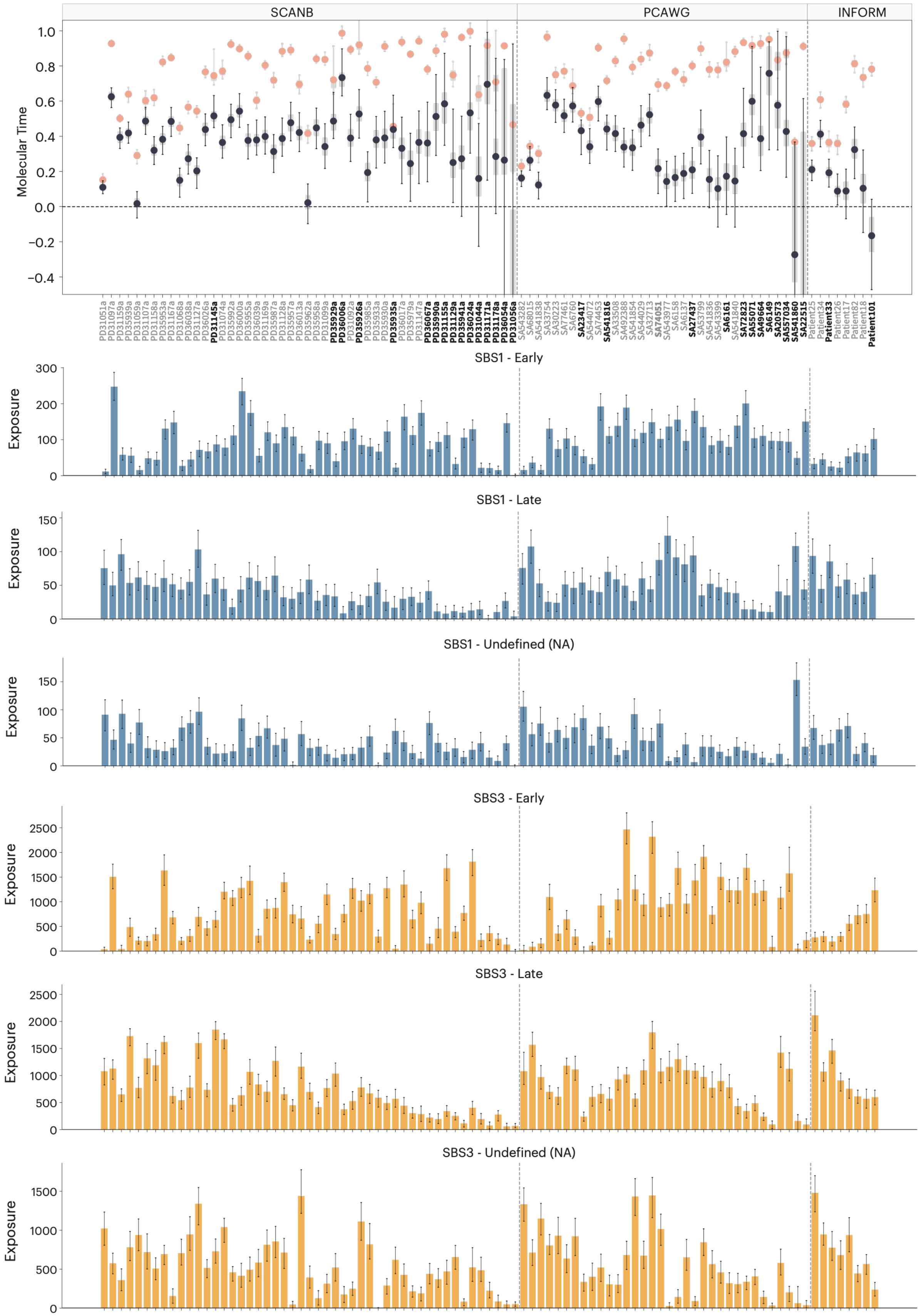
SCANB
PCAWG
INFORM
Molecular Time
SBS1 - Early
SBS1 - Late
SBS1 - Undefined (NA)
SBS3 - Early
SBS3 - Late
SBS3 - Undefined (NA)
Exposure

**Figure S13. Quantification of confidence in molecular (SBS1-based) timing estimates of HRD onset and whole-genome duplication in breast cancers, and sample filtering.** The top panel shows inferred molecular times for HRD onset (navy) and WGD (peach) across samples from three breast cohorts (SCANB, PCAWG, INFORM); error bars denote 95% confidence intervals and gray bars denote the inter-quartile range, both estimated by bootstrapping. The bottom tracks show the corresponding SBS1 and SBS3 exposures, stratified by timing class (Early, Late, Undefined/NA). Samples are sorted by the width of the HRD-onset confidence interval, and samples excluded from the main analysis are shown in bold (see Methods).

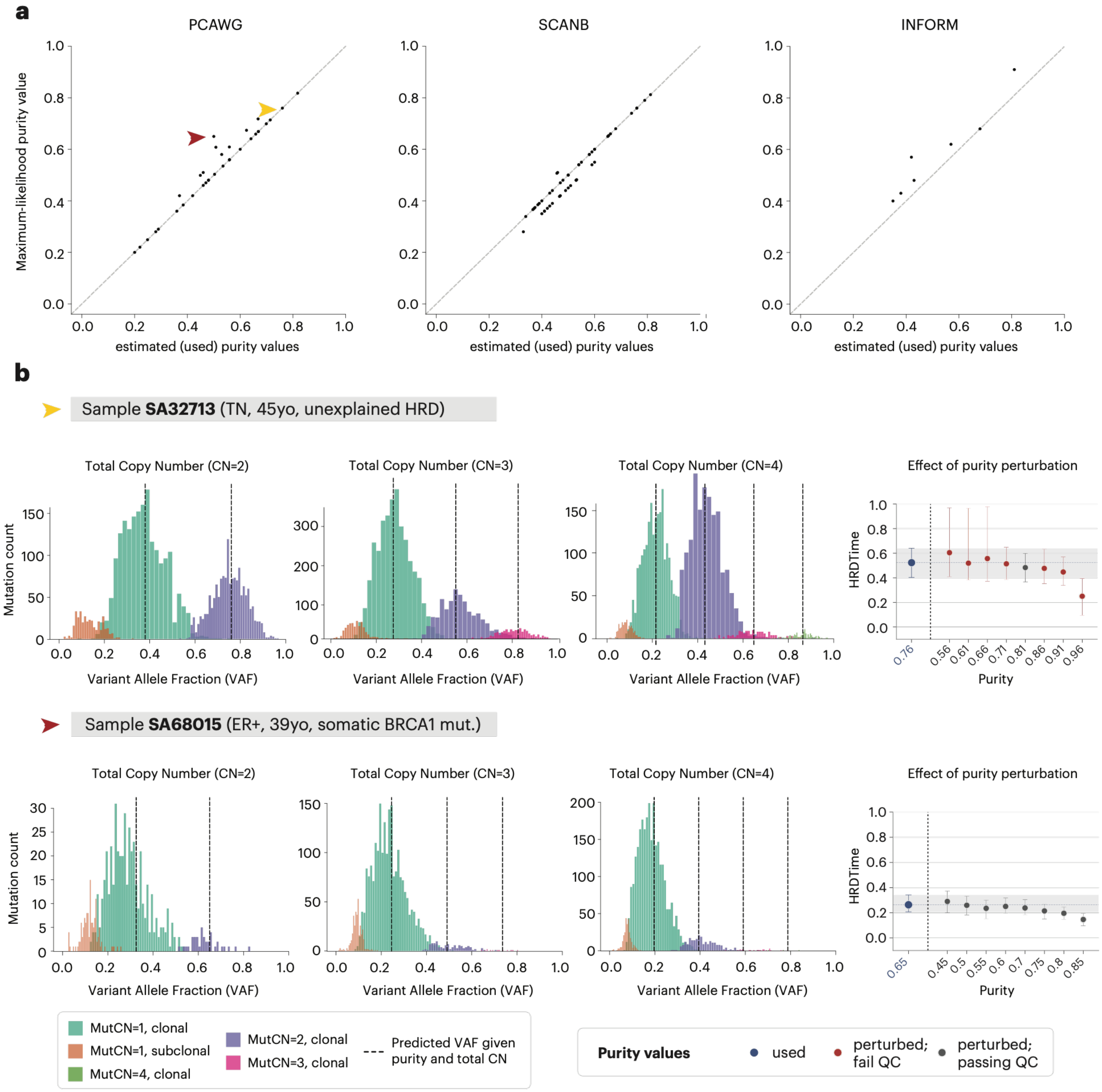


**Figure S14. Validation of purity estimates.** (**a**) Purity values used for timing versus purity values that maximize the likelihood of the observed somatic variant allele fractions (VAFs). Maximum-likelihood purities were obtained by running MutationTimeR across a grid of purity values and recording the post-EM VAF likelihood. Arrows indicate the two samples shown in (b). (**b**) Two example samples with clinical annotations. Top: Sample SA32713 (TN, 45 yr, unexplained HRD), where the HRDTimer purity matched the MutationTimeR maximum-likelihood purity; VAF distributions of somatic mutations, stratified by local total copy number, align with the predicted peaks for each mutation multiplicity. Purities further from the maximum-likelihood value are correctly rejected by MutationTimeR QC. Bottom: SA68015 (ER+, 39 yr, somatic BRCA1 mutation), where the used purity was farthest from the maximum-likelihood value. HRDTimer nonetheless returned consistent HRD onset estimates across purities from 0.45 to 0.7.

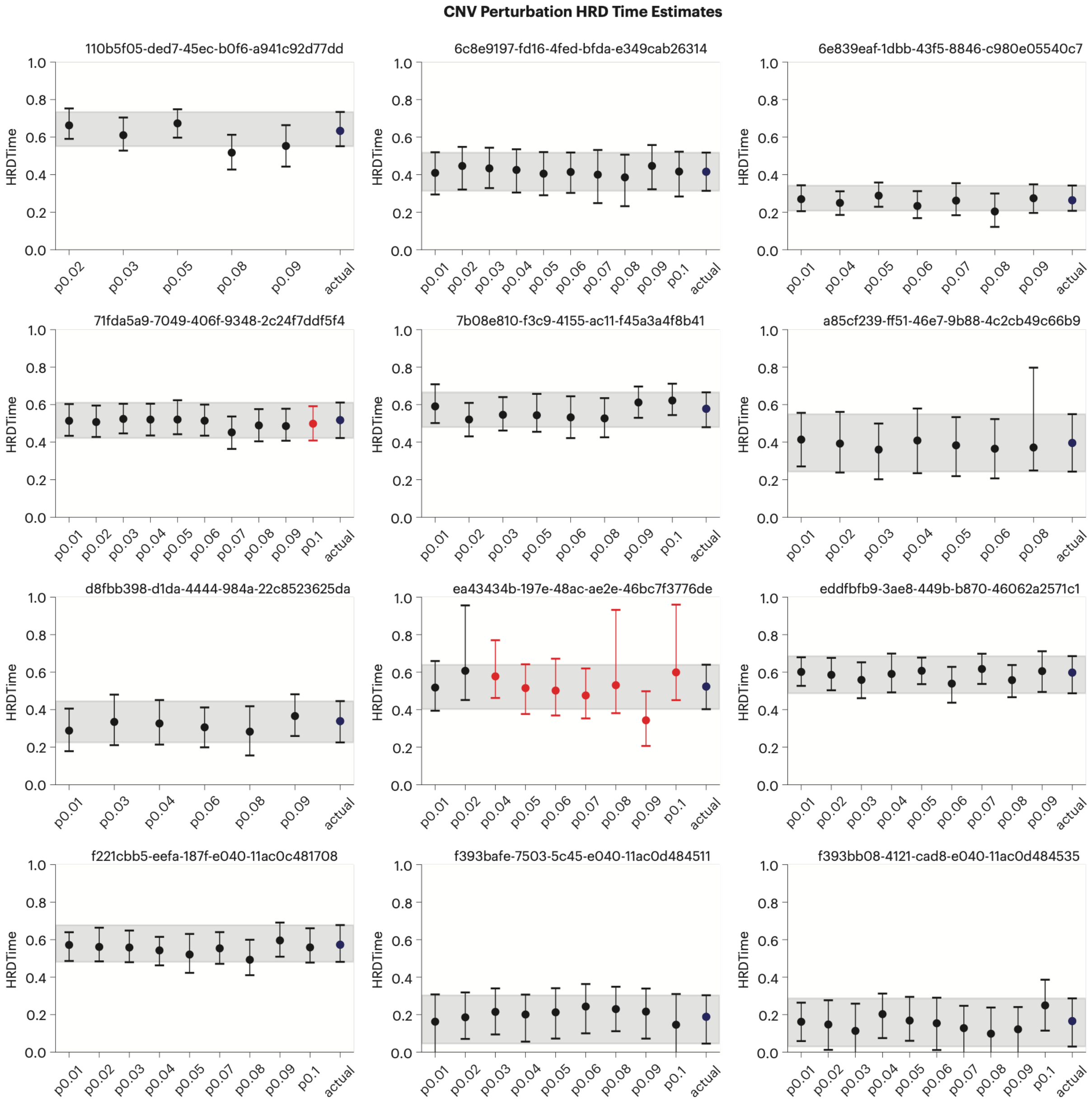


**Figure S15. Perturbing up to 10% of CNV segments does not affect predicted HRD onset.** Twelve randomly selected PCAWG samples, one per panel. For each sample, pX denotes the fraction of CNV segments randomly perturbed by adding or subtracting one chromosome copy (e.g., p0.01 = 1%). The blue dot ("actual", rightmost point) is the HRDTimer estimate from the best-estimate copy number profile used in the manuscript, and the shaded gray band is its 95% confidence interval. Perturbed estimates remain within this band across all perturbation levels. Black dots, analyses passing MutationTimeR QC; red dots, analyses failing QC.

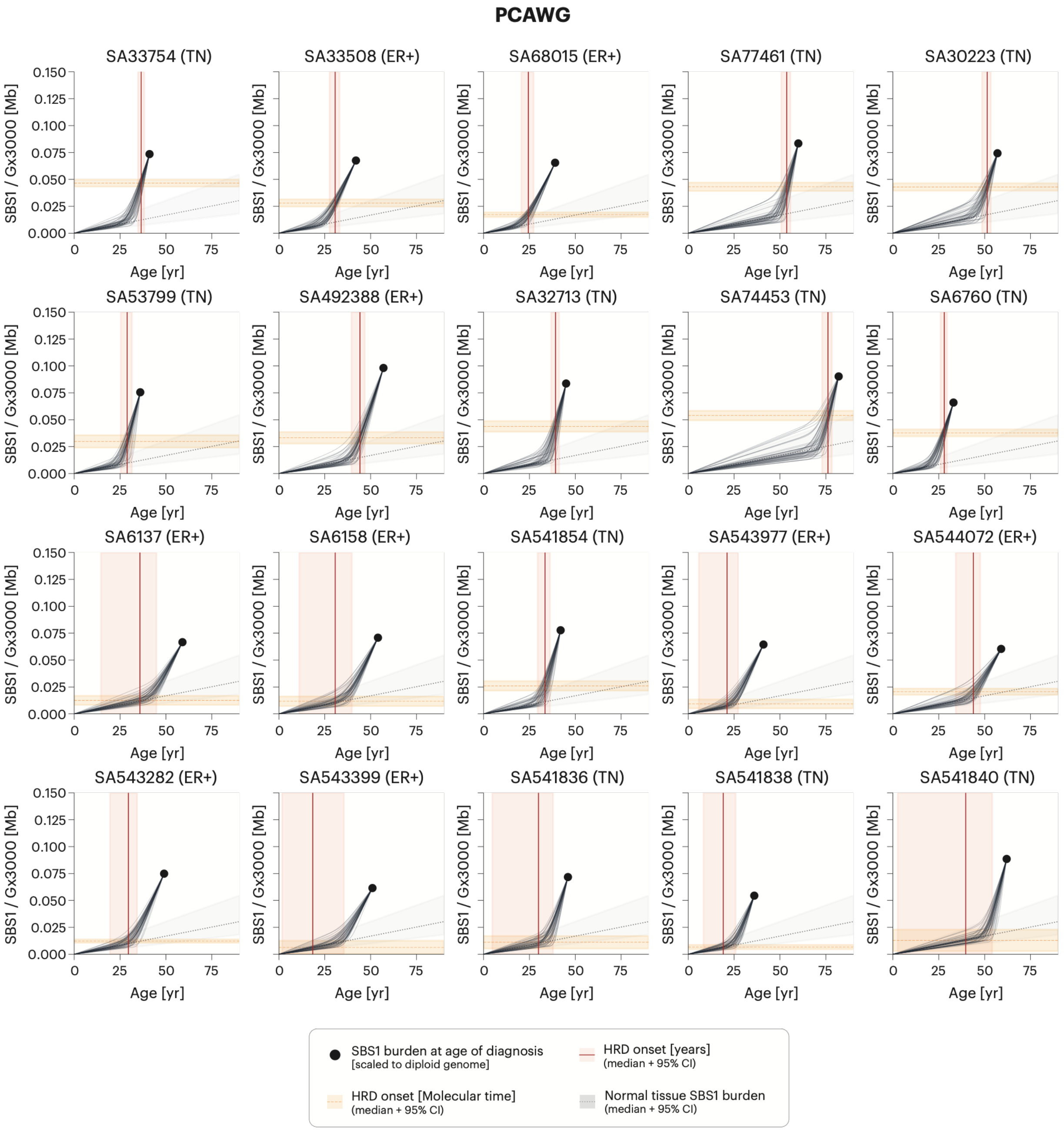


**Figure S16. Example fits of the SBS1–age function in the PCAWG dataset.** Each panel shows the gradual SBS1–age model fitted to one sample, relating SBS1 burden (scaled to a diploid genome, SBS1/G × 3000 Mb) to age. Dark navy curves show the bootstrap ensemble of fitted trajectories, capturing uncertainty in the model parameters and the SBS1-based molecular-time estimate. The black dot marks the SBS1 burden at the age of diagnosis (scaled to diploid genome), which incorporates a fixed estimate of post-MRCA SBS1 mutations (116 per diploid genome) derived from scNanoSeq data to account for cell-private mutations undetectable in bulk sequencing. The orange dashed horizontal band indicates HRD onset in molecular (SBS1-based) time (median and 95% CI); the red vertical line and pink shaded band indicate HRD onset in years (median and 95% CI). The gray shaded band shows the normal-tissue SBS1 burden (median and 95% CI). G denotes effective genome size (see Methods). For the conceptual basis of the SBS1–age models, see Figure 5a.

#### SCANB

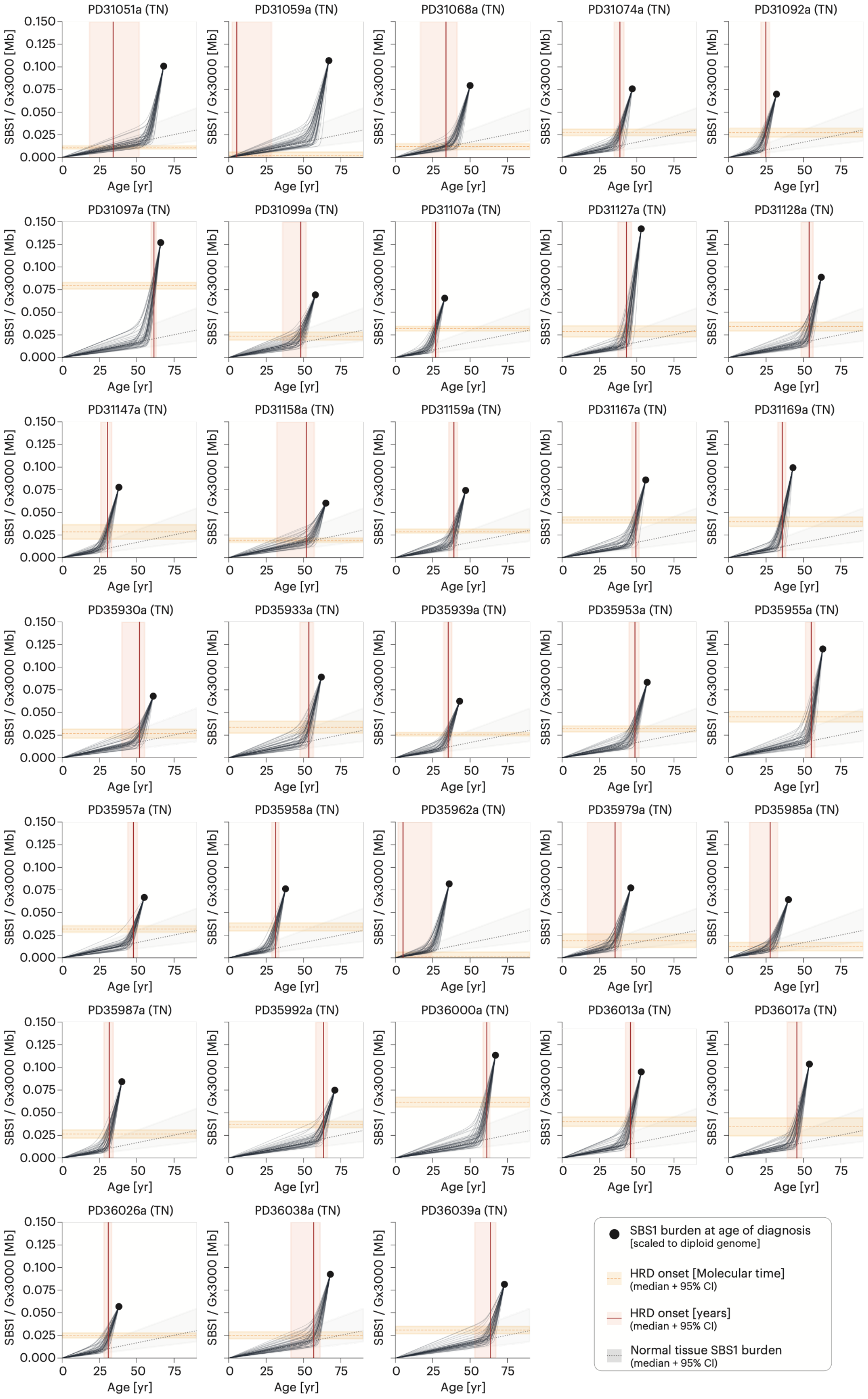
PD31051a (TN)
PD31059a (TN)
PD31068a (TN)
PD31074a (TN)
PD31092a (TN)
PD31097a (TN)
PD31099a (TN)
PD31107a (TN)
PD31127a (TN)
PD31128a (TN)
PD31147a (TN)
PD31158a (TN)
PD31159a (TN)
PD31167a (TN)
PD31169a (TN)
PD35930a (TN)
PD35933a (TN)
PD35939a (TN)
PD35953a (TN)
PD35955a (TN)
PD35957a (TN)
PD35958a (TN)
PD35962a (TN)
PD35979a (TN)
PD35985a (TN)
PD35987a (TN)
PD35992a (TN)
PD36000a (TN)
PD36013a (TN)
PD36017a (TN)
PD36026a (TN)
PD36038a (TN)
PD36039a (TN)
SBS1 / Gx3000 [Mb]
0.150
0.125
0.100
0.075
0.050
0.025
0.000
Age [yr]
0
25
50
75
SBS1 burden at age of diagnosis
[scaled to diploid genome]
HRD onset [Molecular time]
(median + 95% CI)
HRD onset [years]
(median + 95% CI)
Normal tissue SBS1 burden
(median + 95% CI)

**Figure S17. Example fits of the SBS1–age function in the SCANB dataset.** See Figure S16 for the explanation and legend.

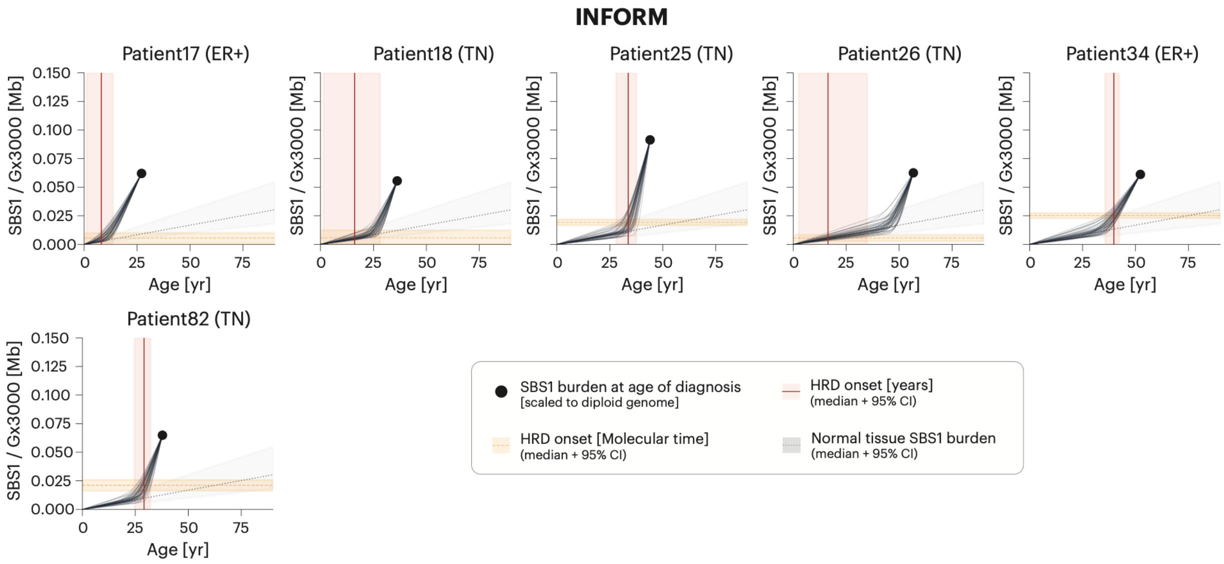


**Figure S18. Example fits of the SBS1–age function in the INFORM dataset.** See Figure S16 for the explanation and legend.

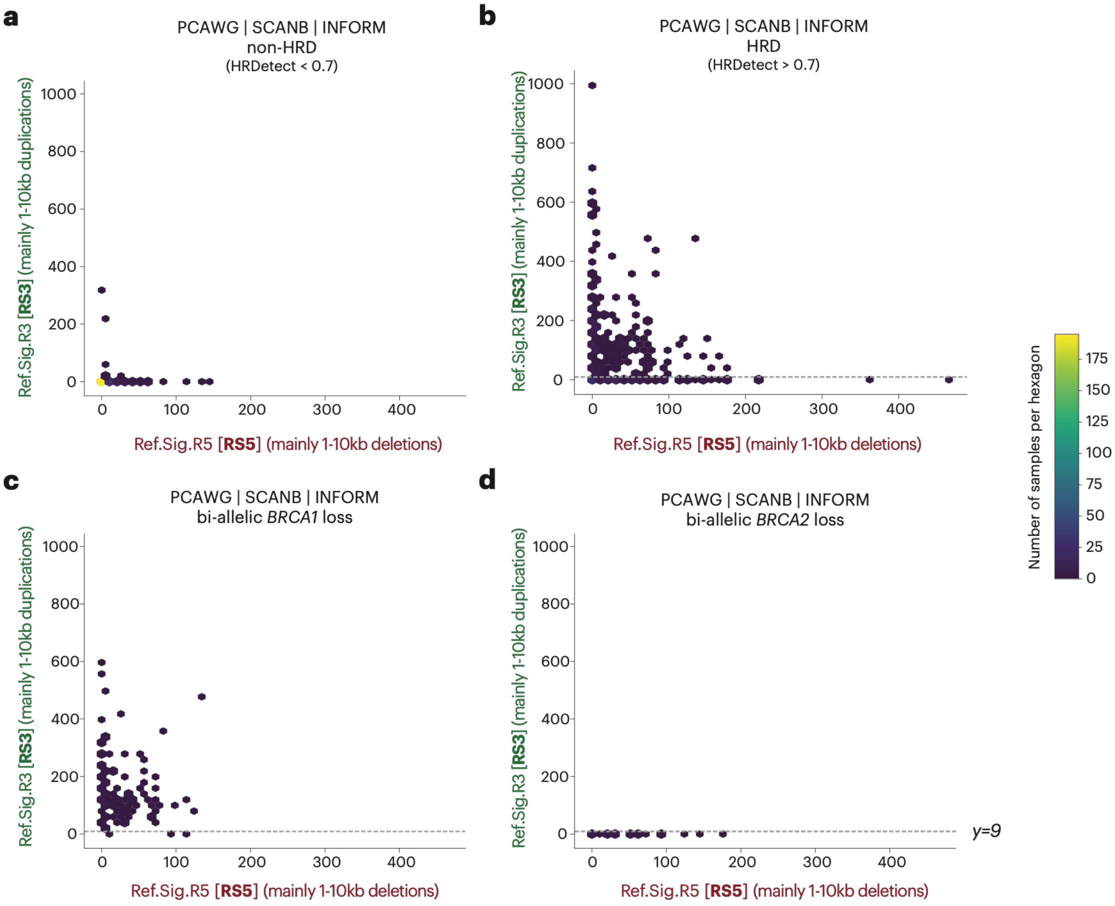


**Figure S19. Classification of HRD samples into *BRCA1*-like and *BRCA2*-like subgroups based on somatic structural variant (SV) patterns.** Axes show the abundance of RS5- (x) and RS3-attributed (y) SVs (rearrangement signatures) in each panel; the same axes apply to all panels. (**a**) Non-HRD samples. (**b**) HRD samples. (**c**) Tumors with biallelic *BRCA1* loss, which show higher RS3 abundance. (**d**) Tumors with biallelic *BRCA2* loss. Panels (c) and (d) motivated a threshold of $\geq 9$ RS3-attributed SVs to separate *BRCA1*-like from *BRCA2*-like phenotype.

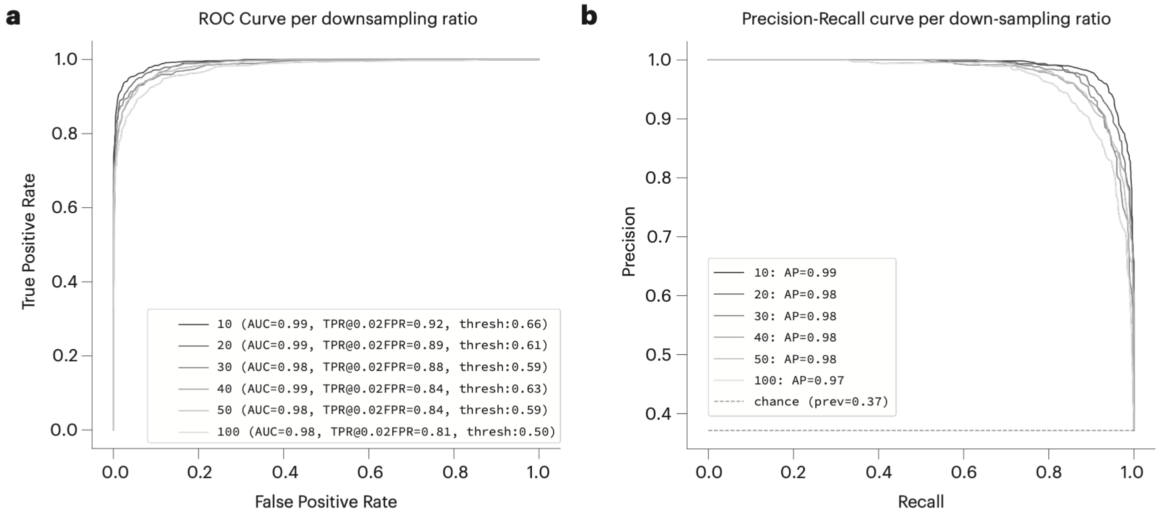


**Figure S20. Cross-validation performance of HRDscout.** HRDscout was trained on bulk whole-genome sequencing (WGS) data combining three previously published breast-cancer cohorts: the 560-breast cancer cohort, SCANB, and INFORM. Of the 847 samples in the combined catalog, 112 were excluded by mutation-burden filtering (50 < indel count < 5,000 and SNV count < 20,000), yielding 735 retained samples. All 735 had an assigned HRDetect class label: 273 HRDetect-high positives and 462 HRDetect-low/intermediate negatives (source-cohort positive prevalence 0.371). To model mutation-detection sensitivities representative of single-cell WGS technologies, including levels below scNanoSeq's median sensitivity of 2.5%, each sample was downsampled at eight ratios (10×, 20×, 30×, 40×, 50×, 100×, 120×, and 150×), each repeated two to three times. This yielded a synthetic training matrix of 15,435×179 (samples × features), comprising 96 SBS (SBS96) and 83 indel (ID83) channels. HRDscout was evaluated by stratified 10-fold cross-validation, with folds randomly partitioned by HRDetect label. Folds were not stratified by simulated detection sensitivity, so each training and held-out partition spanned all down-sampling ratios; per-sensitivity metrics were obtained by post hoc stratification of out-of-fold predictions. (**a**) Receiver operating characteristic (ROC) curve. (**b**) Precision-recall curve; the dashed line indicates the chance level (precision = prevalence = 0.37). Both are stratified by down-sampling ratio.

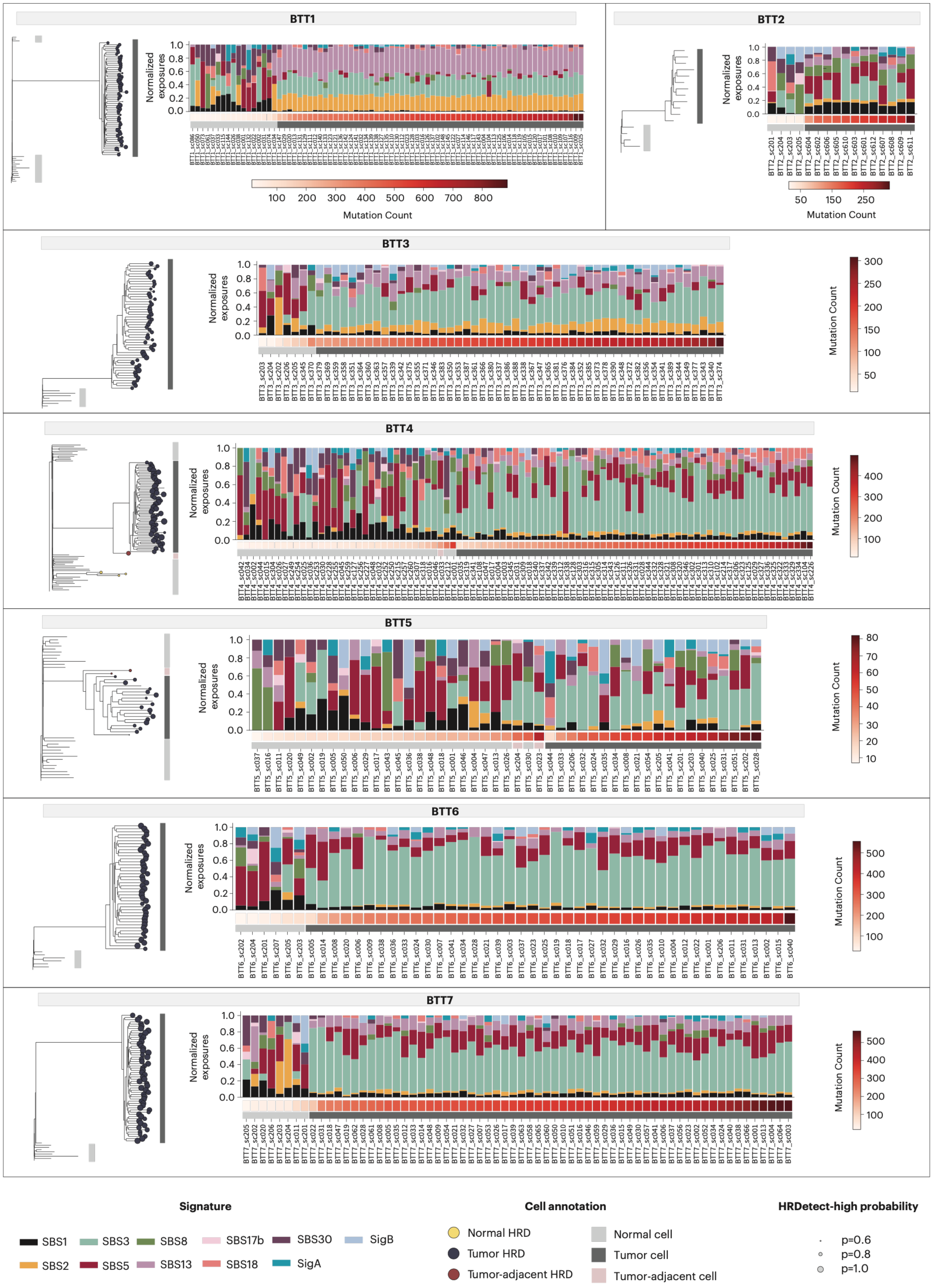
BTT1
BTT2
BTT3
BTT4
BTT5
BTT6
BTT7
Normalized exposures
Mutation Count
Signature
SBS1
SBS2
SBS3
SBS5
SBS8
SBS13
SBS17b
SBS18
SBS30
SigA
SigB
Cell annotation
Normal HRD
Tumor HRD
Tumor-adjacent HRD
Normal cell
Tumor cell
Tumor-adjacent cell
HRDetect-high probability
p=0.6
p=0.8
p=1.0

**Figure S21. Per-cell mutational signature fits for BTT (triple-negative) scNanoSeq samples.** For each of the seven BTT samples, every bar shows the normalized fitted signature exposures for an individual cell, across the signature panel indicated in the legend. Cells were sorted first by tumor/normal classification, then by total detected mutation burden. The colored annotation bar beneath each plot gives the cell classification (normal, tumor, or tumor-adjacent, with HRD status indicated), and the adjacent color scale shows per-cell mutation count. On the accompanying phylogenetic trees, dots mark cells classified as HRD-high by HRDscout, with dot size scaled by HRDetect-high probability.

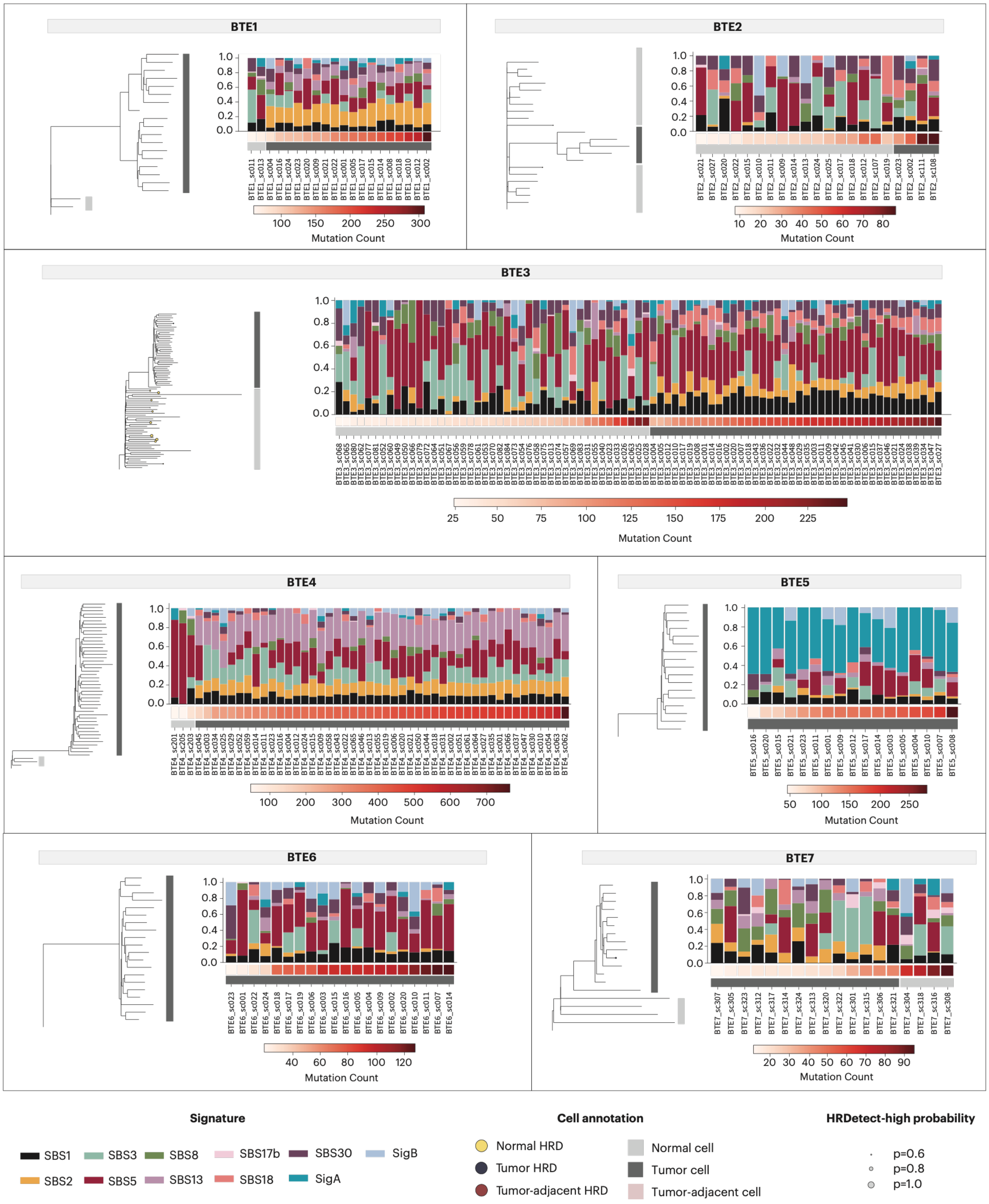


**Figure S22. Per-cell mutational signature fits for BTE (ER+) scNanoSeq samples.** See Figure S21 for details.

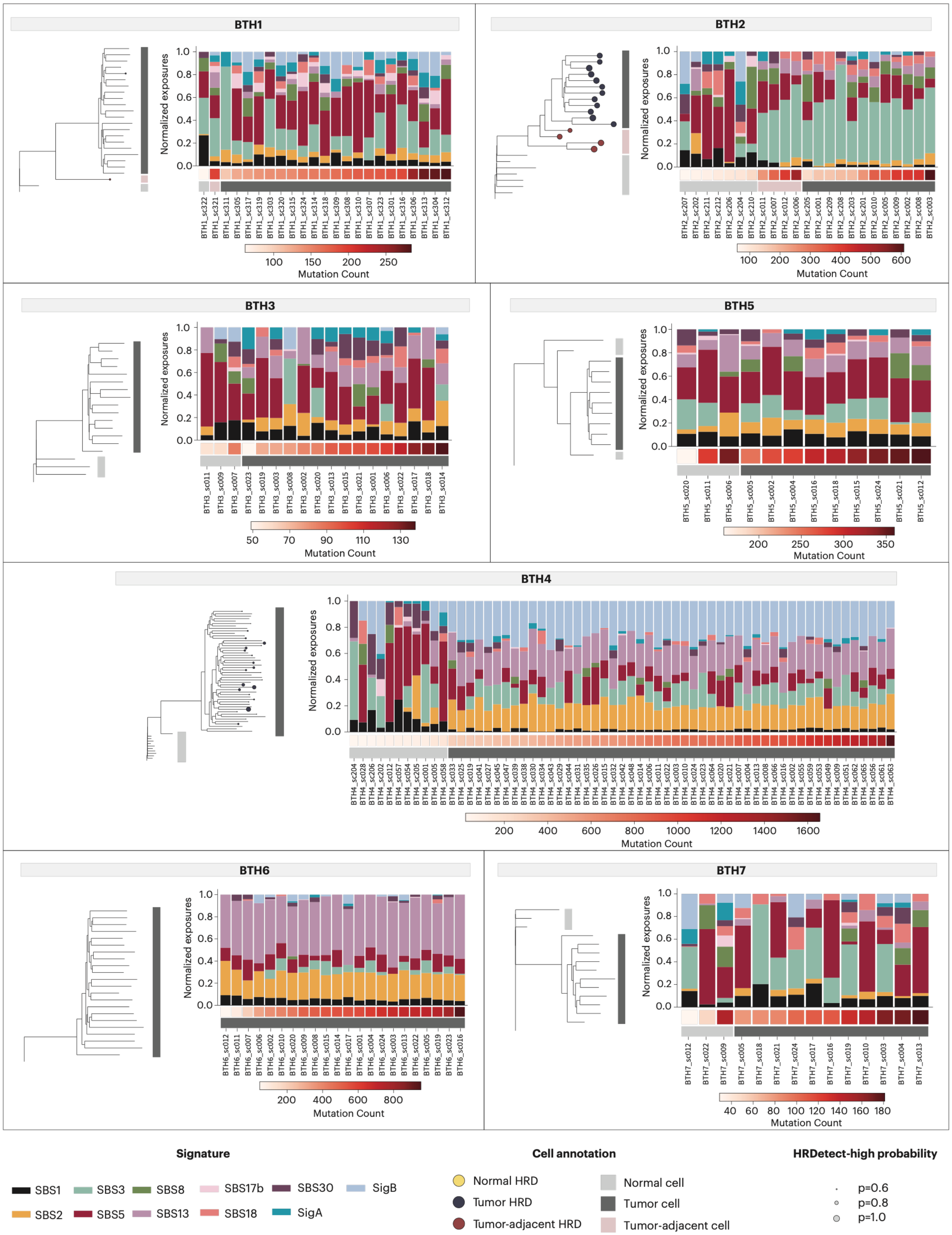


**Figure S23. Per-cell mutational signature fits for BTH (HER2+) scNanoSeq samples.** See Figure S21 for details.

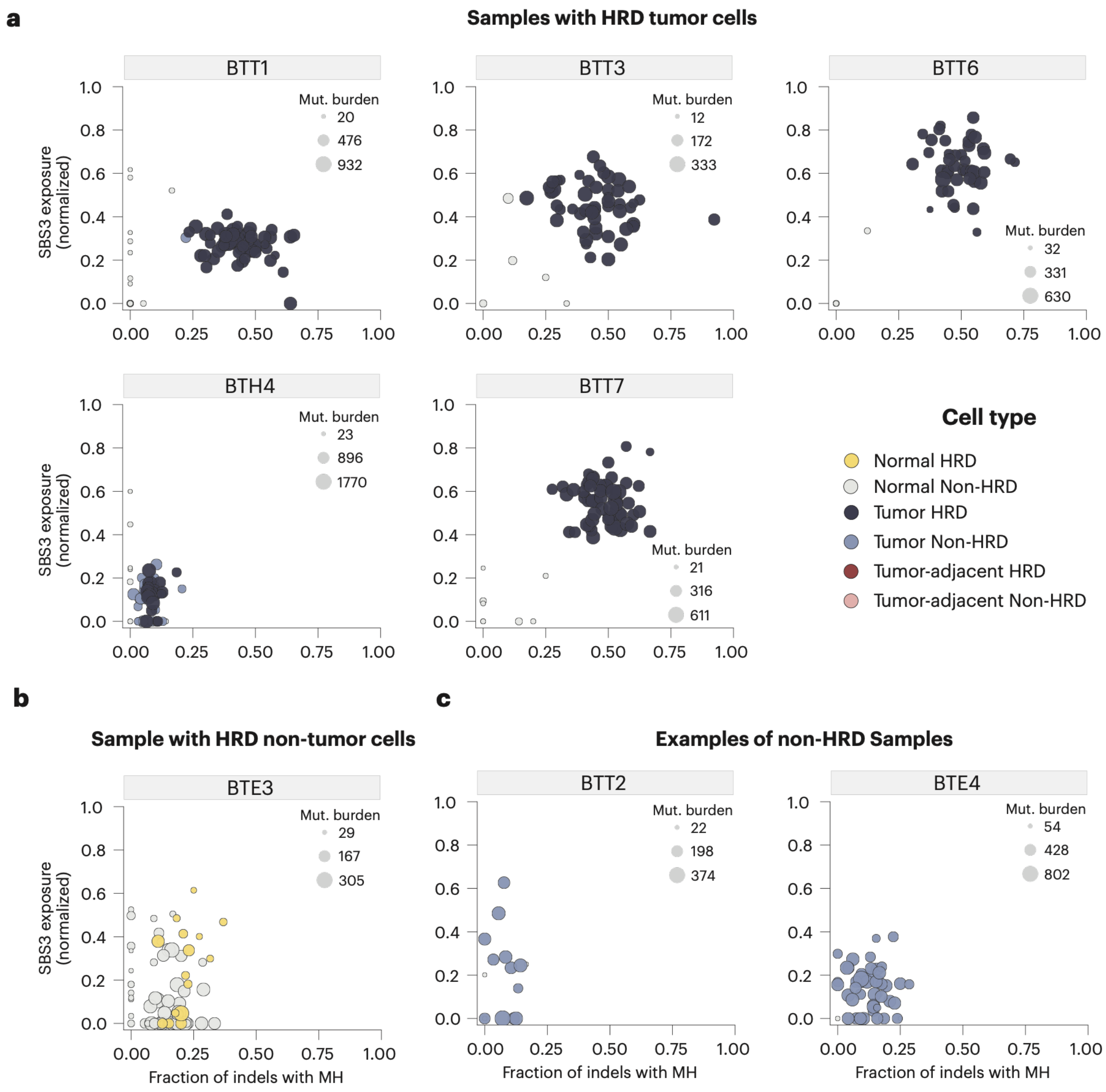


**Figure S24. HRDscout predictions across samples and cell classes.** Cells were classified as tumor, tumor-adjacent, or normal based on the lineage tree, and HRD status was assigned using HRDscout. Scatter plots show normalized SBS3 exposure (y-axis) versus the fraction of indels with microhomology (MH; x-axis) — the two hallmarks of HRD — with points colored by combined cell class and HRD status (see legend) and sized by per-cell mutation burden. (**a**) Samples in which tumor cells were predominantly classified as HRD (BTT1, BTT3, BTT6, BTH4, BTT7). (**b**) Sample BTE3, in which some non-tumor cells were classified as HRD. (**c**) Non-HRD samples (BTT2, BTE4).

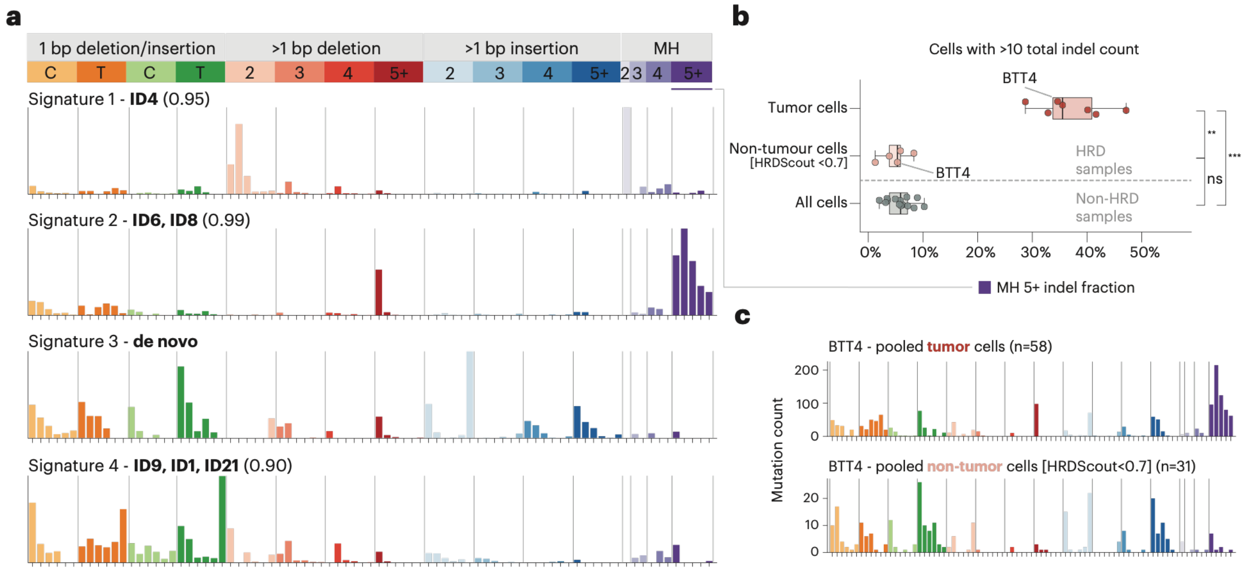


**Figure S25. Indel mutational signatures in scNanoSeq data of HRD breast tumors.** (**a**) De novo indel (ID83) signatures extracted from somatic indels across single cells, called from duplex sequencing reads. For each signature the best-matching COSMIC v3.2 reference signature(s) and cosine similarity are shown: Signature 1 (ID4, 0.95), Signature 2 (ID6/ID8, 0.99) and Signature 4 (ID9/ID1/ID21, 0.90); Signature 3 is de novo with no confident COSMIC match. Signature 2 (ID6/ID8), dominated by ≥5 bp microhomology (MH)-flanked deletions, reflecting HR deficiency. (**b**) Fraction of MH-flanked deletions ≥5 bp (MH 5+ indel fraction) among cells with > 10 total indels, compared across pooled tumor cells and pooled non-tumor cells (HRDscout < 0.7) from seven HRD samples, and all cells from non-HRD samples; BTT4 is highlighted. Boxplots show median and IQR, whiskers extend to 1.5× IQR, and points denote individual samples. Two-sided Mann–Whitney U: tumor vs non-tumor (HRD) cells, $P < 0.01$ (**); tumor (HRD) vs non-HRD cells, $P < 0.001$ (***); non-tumor (HRD) vs non-HRD cells, not significant (ns). (**c**) Pooled ID83 spectra for BTT4 tumor (n = 58) and non-tumor (n = 31, HRDscout < 0.7) cells; note the differing y-axis scales.

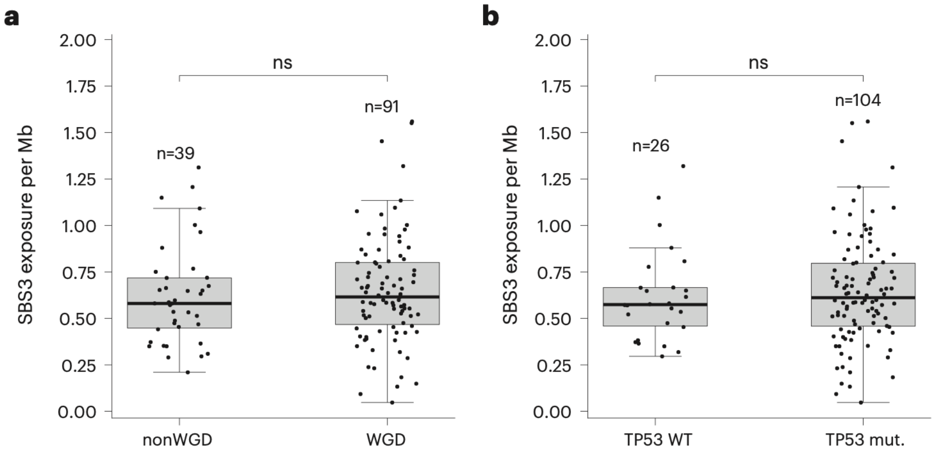


**Figure S26. SBS3 exposure per Mb does not change across major genomic transitions.** (**a**) SBS3 exposure per Mb (corrected for effective genome size) in HRD tumors stratified by WGD status (non-WGD, n = 39 [PCAWG = 28, INFORM = 11]; WGD, n = 91 [SCANB = 49, PCAWG = 34, INFORM = 8]). (**b**) SBS3 exposure per Mb in HRD tumors stratified by TP53 mutation status (TP53-wild-type, n = 26 [PCAWG = 16, INFORM = 7, SCANB = 3]; TP53-mutant, n = 104 [PCAWG = 46, SCANB = 46, INFORM = 12]). Box plots show median and interquartile range; whiskers extend to 1.5× IQR. Individual tumors are overlaid as points. SBS3 burden is not significantly different between groups in either comparison (two-sided Wilcoxon rank-sum test; ns, P > 0.05), arguing against changes in SBS3 mutational activity at WGD or TP53 loss

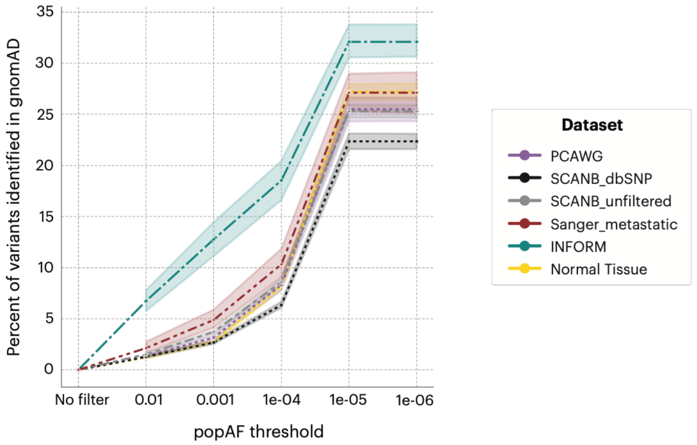


**Figure S27. Assessment of potential germline contamination in bulk datasets through overlap between somatic mutation sets and a germline variant database.** For each dataset, the overlap rate between its somatic mutation calls and population variants in the gnomAD database was computed as a proxy for germline contamination. Elevated overlap rates indicated potential germline contamination in the INFORM somatic mutation dataset. To address this, variants present in gnomAD at a population allele frequency ≥ 0.1% ((≥1 in 1,000 individuals) were excluded from the INFORM dataset. The unfiltered SCANB call set (SCANB_unfiltered) showed no evidence of germline contamination and was retained for the main analysis.

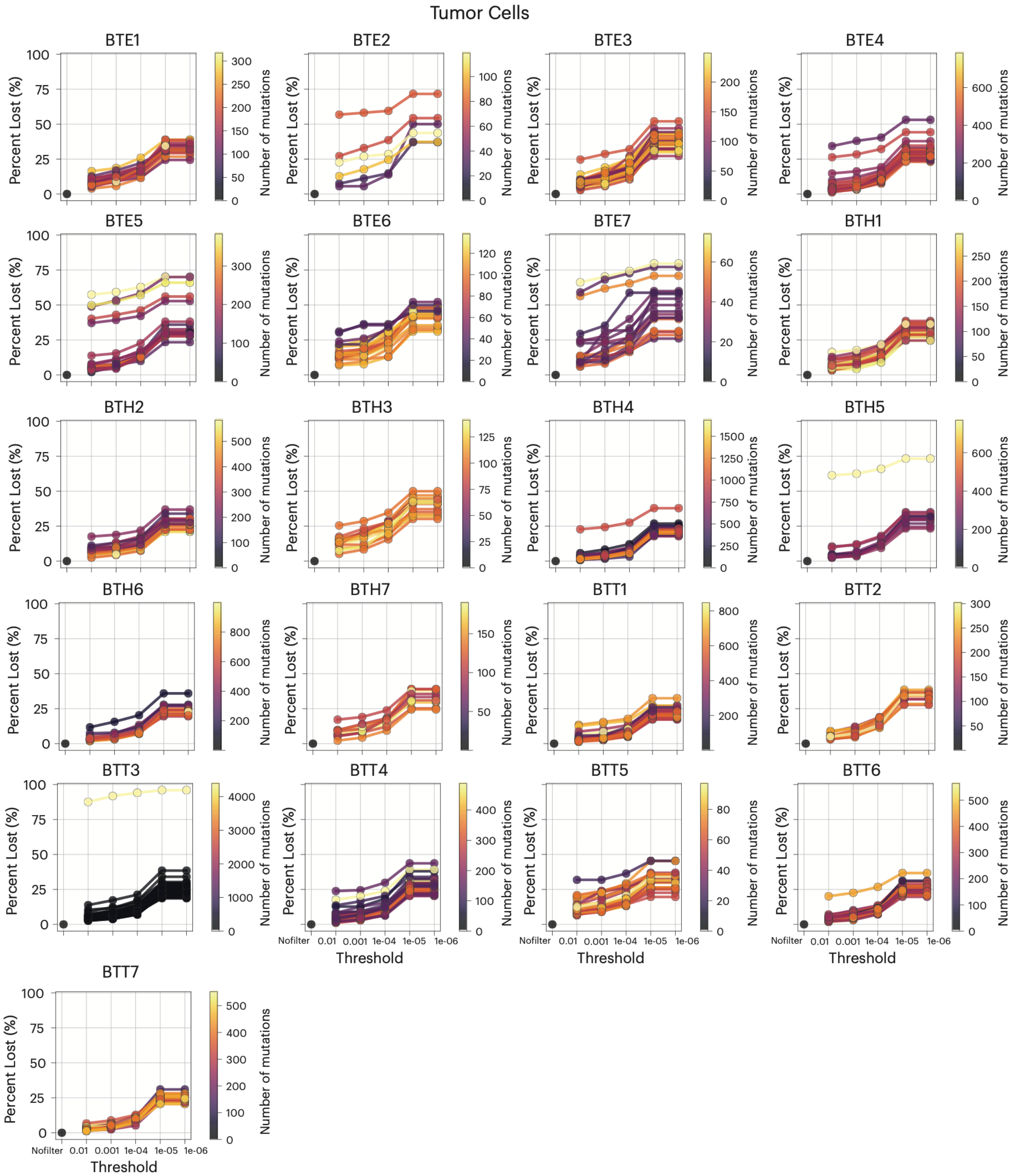


**Figure S28. Assessment of potential germline contamination of somatic mutation calls in tumor cells profiled with scNanoSeq.** The x-axis shows the population allele frequency threshold used for filtering (including a no-filter baseline); the y-axis ("Percent Lost") indicates the percentage of a cell's somatic mutations that overlap gnomAD germline variants above the given threshold. Each line represents one cell, colored by its total mutation count (color scale). Two filters were applied: (i) cells with > 40% overlap at the 0.001 threshold were excluded entirely, and (ii) within retained cells, individual mutations overlapping common gnomAD variants (population allele frequency ≥ 0.1%, ≥ 1 in 1,000) were removed. This filtering was performed to ensure the accuracy of downstream mutational signature analysis on the retained set of somatic mutations.

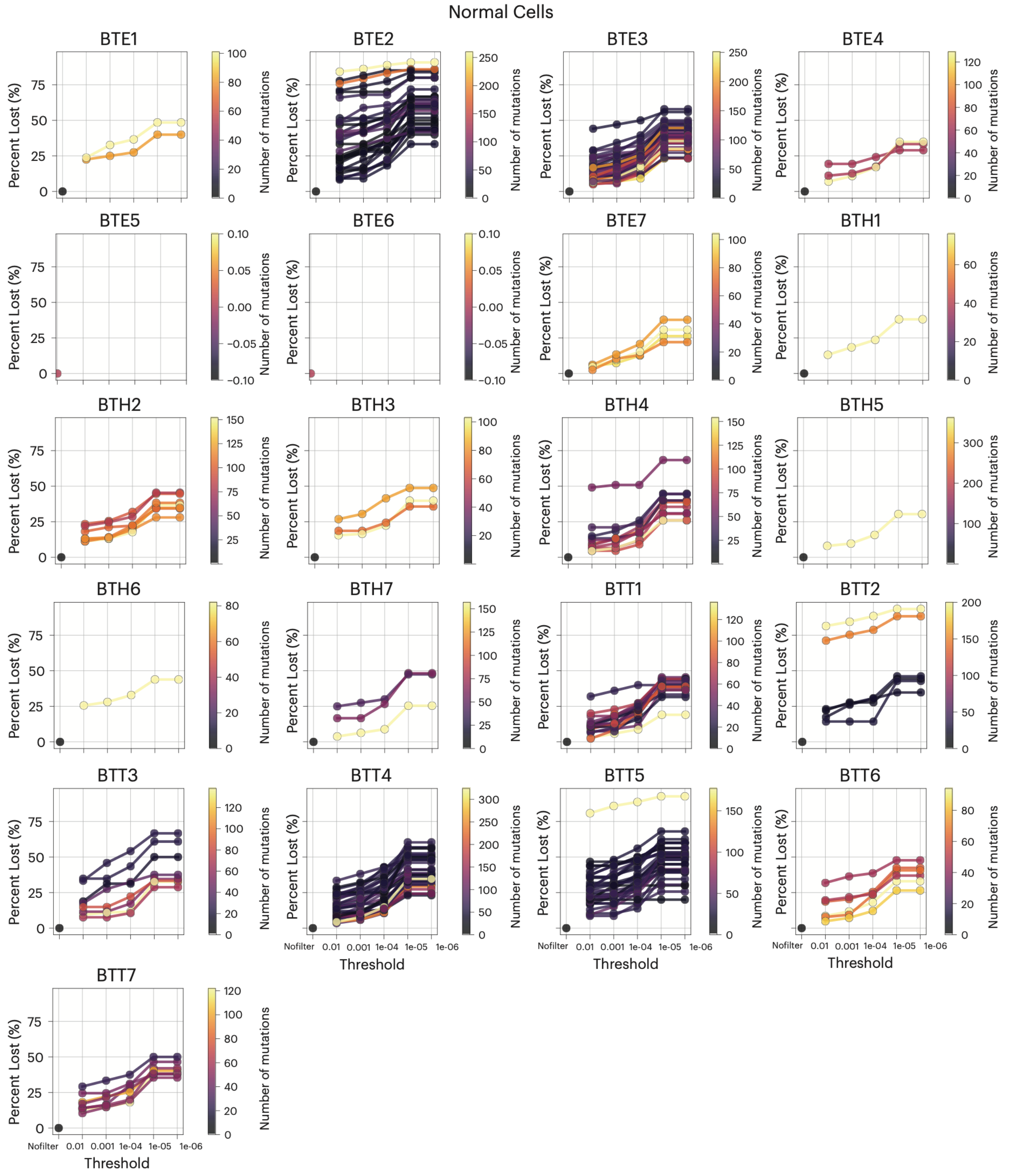


**Figure S29. Assessment of potential germline contamination of somatic mutation calls in normal cells profiled with scNanoSeq.** Same process as explained in Figure S28 caption.

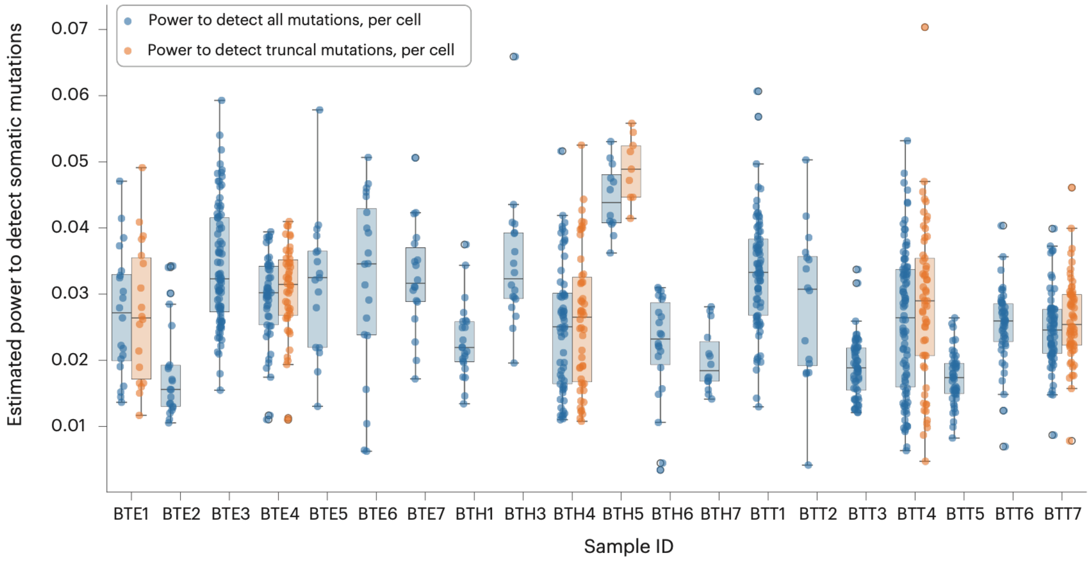


**Figure S30. Power for detecting mutations in cells using scNanoSeq.** For each sample, per-cell estimated power is shown for detecting all somatic mutations (blue) and truncal mutations (orange). Truncal mutations are those detected in a cell that also appear in the matched bulk WGS calls; these values were used to correct the detected mutation burden per cell. Boxplots show median and interquartile range, whiskers extend to 1.5× IQR, and points represent individual cells. Sample BTH2 is not included because power could not be estimated, owing to copy number exceeding 5 across almost all genomic regions.

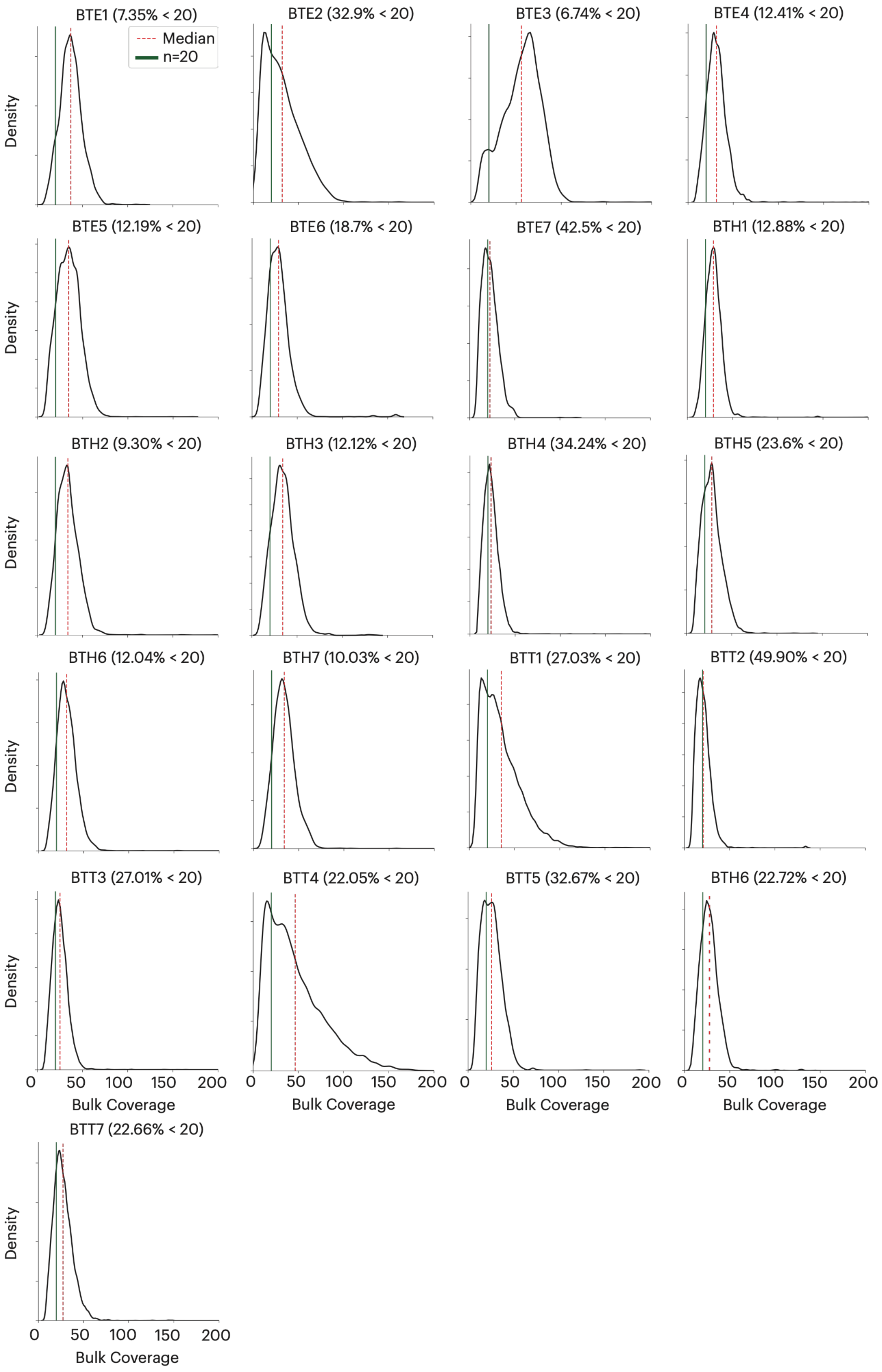
BTE1 (7.35% < 20)
Median
n=20
BTE2 (32.9% < 20)
BTE3 (6.74% < 20)
BTE4 (12.41% < 20)
BTE5 (12.19% < 20)
BTE6 (18.7% < 20)
BTE7 (42.5% < 20)
BTH1 (12.88% < 20)
BTH2 (9.30% < 20)
BTH3 (12.12% < 20)
BTH4 (34.24% < 20)
BTH5 (23.6% < 20)
BTH6 (12.04% < 20)
BTH7 (10.03% < 20)
BTT1 (27.03% < 20)
BTT2 (49.90% < 20)
BTT3 (27.01% < 20)
BTT4 (22.05% < 20)
BTT5 (32.67% < 20)
BTH6 (22.72% < 20)
BTT7 (22.66% < 20)
Density
0
50
100
150
200
Bulk Coverage

**Figure S31. Estimation of genome-wide coverage distribution in matched-normal sc-NanoSeq samples.** The single-cell somatic mutation dataset was restricted to mutations with a matched germline sample sequencing coverage of at least 20×, ensuring high-confidence variant calls. The distribution of sequencing coverage at germline mutation loci allows us to approximate a genome-wide distribution. The 20× threshold is indicated by the vertical green line, and the percentage of mutations with coverage below 20× is given in each panel title. Regions with lower coverage in the matched-normal sample are unsuitable for reliable mutation calling; their extent is therefore accounted for when calculating the corrected mutation burden. The median coverage in the matched-normal sample is indicated by the dashed red line.

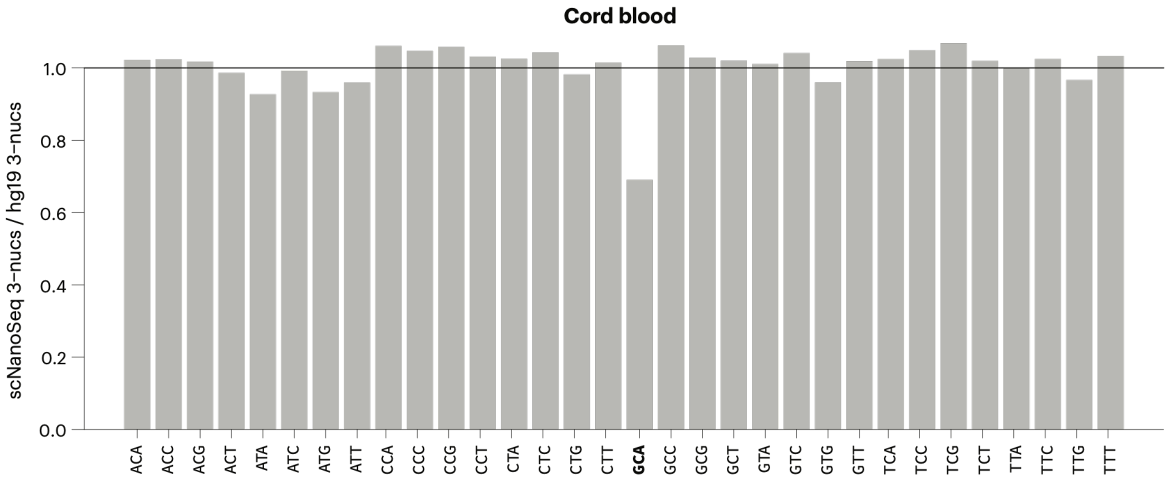


**Figure S32. Tri-nucleotide sequence content of duplex reads in a cord blood sample characterized with scNanoseq.** We used cord blood, a scNanoseq benchmarking sample, to assess coverage bias of this technology. The frequency of each trinucleotide in duplex reads amenable to variant calling was compared to its frequency in the reference genome (hg19). Most sequence contexts are represented proportionally; however, the GCA context is underrepresented in scNanoSeq reads. This motif is part of the recognition site for the restriction enzyme used during library preparation. Other than this trinucleotide, based on relatively flat coverage of trinucleotides, we did not apply a trinucleotide correction before signature analysis.

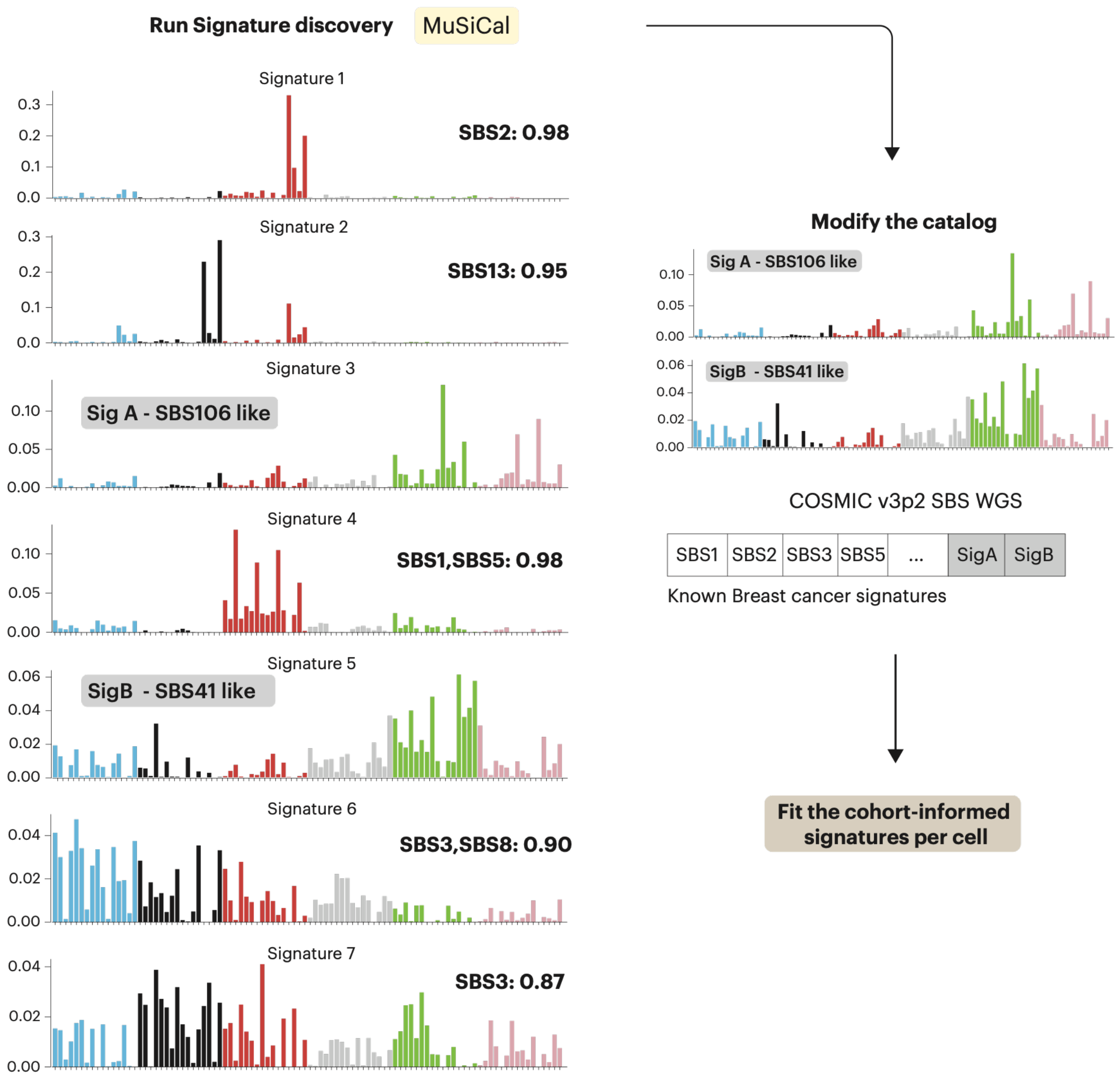


**Figure S33. Mutational signatures identified from point mutations using per-edge analysis of single-cell data profiled with scNanoSeq.** Signatures were extracted with MuSiCal from mutations assigned to individual edges of the phylogenetic tree, ensuring mutually exclusive mutation sets per branch. The discovered signatures were compared to the COSMIC v3.2 reference set using cosine similarity, and we also attempted to explain them as combinations of common signatures. The cosine similarity between each discovered signature and its closest COSMIC match (singleton or pair) is indicated above the corresponding spectrum. Most discovered signatures matched a COSMIC signature directly or could be explained by a linear combination of two common signatures. Two signatures had no close COSMIC match and were treated as de novo (SigA, SBS106-like; SigB, SBS41-like); both were also observed in the original publication. For per-cell signature fitting, we used the COSMIC signatures closest to our rediscovered signatures together with these two de novo signatures.

### Supplementary Tables

1. Metadata for bulk whole-genome sequencing (WGS) samples with HRD and/or WGD status. (**SupTable1_WGS_metadata.xlsx**)
2. Robustness of mutational-signature fitting to low mutation burdens: SBS1/SBS3 exposures across downsampled truncal mutation sets, matching burdens common in scNanoSeq cells. (**SupTable2_trunk_signature_stability_downsampling.xlsx**)
3. Evaluation of SBS5 as an alternative molecular clock: SBS5 exposures versus age and the joint SBS1/SBS5/SBS3 exposure behaviour used to justify adopting SBS1. (**SupTable3_SBS5_age_and_SBS1_SBS5_vs_SBS3.xlsx**)
4. Timing of HRD-attributed structural variants relative to WGD (proportion of late SVs, pi_late). (**SupTable4_SV_pi_late.xlsx**)
5. Timing of HRD-attributed indels (ID6/ID8) relative to WGD (proportion of late indels, pi_late). (**SupTable5_ID6_ID8_pi_late.xlsx**)
6. Late clonal versus subclonal SBS3:SBS1 activity ratios in bulk WGS (GEL cohort), testing stability of the relative rate after HRD onset. (**SupTable6_subclonal_bulk.xlsx**)
7. Subclonal versus cell-private SBS3:SBS1 activity ratios in scNanoSeq single cells, extending the SBS3:SBS1 stability test to the cell-private compartment. (**SupTable7_scNanoSeq_subclonal.xlsx**)
8. HRDTimer estimates (SBS1-based molecular time of HRD onset and WGD, and derived ages) in breast cancer WGS samples with HRD & WGD. (**SupTable8_HRDTimer_estimates.xlsx**)
9. HRDTimer molecular time estimates of HRD onset and WGD in PCAWG ovarian cancers. (**SupTable9_Ovary_HRDTime_WGDTime.xlsx**)
10. SBS1 mutation-burden estimates in breast cancers, including WGD samples with or without HRD, with covariates used for the burden regression. (**SupTable10_SBS1_burden.xlsx**)
11. SBS1 burden in matched primary-relapse breast cancer pairs (scaled per diploid genome) with inter-sample accumulation rates. (**SupTable11_scaledSBS1_Age_Primary_Metastasis_SangerStudy.xlsx**)
12. Per-cell mutation burden, mutational-signature exposures, and mutation-detection power across 21 breast cancers profiled by scNanoSeq. (**SupTable12_scNanoseq_allResults.xlsx**)
13. Performance metrics for HRDscout predictions across mutation-detection-power strata (sensitivity, specificity, AUC); estimates from cross-validation. (**SupTable13_HRDScout_performance.xlsx**)
14. De novo indel (ID83) signatures discovered in scNanoSeq data, with best-matching COSMIC references and cosine similarities. (**SupTable14_Discovered_deNovo_ID_Signatures.csv**)
15. Per-cell HRDscout predictions for cells profiled by scNanoSeq. (**SupTable15_PerCell_HRDScout_predictions.xlsx**)
16. Pooled fraction of long (>=5 bp) microhomology-flanked deletions in scNanoSeq cell groups (HRD tumor, non-tumor, and non-HRD control cells). (**SupTable16_scNanoSeq_MHindel_fractions.xlsx**)
17. SBS3 exposure per Mb stratified by WGD status and by TP53 mutation status, showing SBS3 burden does not change across these genomic transitions. (**SupTable17_SBS3_burden_WGD_TP53.xlsx**)
