## Supplementary material for "Timing the onset of homologous recombination deficiency before breast cancer diagnosis": Supplementary Note 1 - A mathematical framework for power correction of a single cell duplex sequencing assay (scNanoSeq).pdf

Michail Andreopoulos

### 1 Define notation

The following notation will be used throughout this section.

- Mutation: A detected alteration in a DNA fragment that **meets** the a4s2 criterion; at least 4 reads cover the locus, with at least one read per allele.
- Variant: A detected alteration in a DNA fragment, regardless of whether it meets the a4s2 criterion.
- UMI: Unique Molecular Identifier used to tag a single DNA fragment originating from a chromosomal copy. Each chromosomal copy can be covered by at most one UMI at a given locus. For simplicity, the term UMI count will be used to refer to the number of unique DNA fragments (chromosomal copies) in which a *variant* is present, at a given locus.
- $T$: Total copy number of a given chromosomal region, $T \in \mathbb{N}_{\neq 0}$
- $M_1$: Number of chromosome copies inherited from parent 1 that subsequently might have undergone gains or was lost $M_1 \in \mathbb{N}$
- $M_2$: Number of chromosome copies inherited from parent 2 that subsequently might have undergone gains or was lost $M_2 \in \mathbb{N}$
- $M$: Minor copy number $M = \min(M_1, M_2)$. The copy number of a chromosomal region will be characterized as $(T, M)$. (The case of $M_1 = M_2 = 0$ means homozygous deletion, observed only for smaller chromosomal regions)
- $m$: Multiplicity of a mutation, i.e. how many chromosomal copies carry the mutation at a specific locus ($m \leq T$)
- $n$: Total UMI counts at a given locus. In a single cell, $n \leq T$.

### 2 Motivation

Single-cell duplex sequencing technologies tag each sequenced DNA fragment with a unique molecular identifier (UMI). Therefore, no more than one UMI can be linked to a single chromosomal copy at a specific locus in a single cell. This inherent property allows the UMI count to capture information about the copy number of a locus.

The power to detect a mutation depends, among other factors, on its multiplicity. In chromosomal regions covered by sequencing reads, if a mutation occurs in multiple chromosomal copies, several DNA fragments—each tagged with a unique UMI—will carry the mutation, increasing the statistical likelihood of detection (i.e., mutation detected on both strands passing the a4s2 criterion on at least one UMI-tagged DNA fragment). To accurately estimate the total mutational burden using duplex technologies, it is essential to account for how detection power varies with multiplicity, which is limited by the total copy number in a region. Therefore, regions with different copy numbers should be corrected separately.

A DNA fragment tagged with a single UMI can carry a number of variants. For each variant at a given locus, we can count the total number of times it is present across different chromosomal copies by counting how many UMIs report that variant at the corresponding position. For a variant to be called from scNanoSeq data, the following conditions must hold:

- Variant detected in both strands

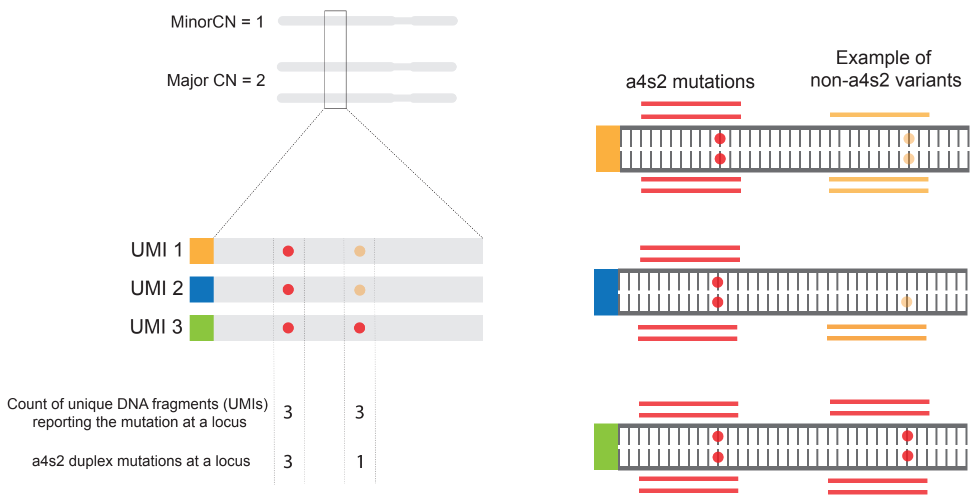


**Figure 1.** Schematic diagram illustrating the a4s2 criterion for identifying a duplex mutation in a genomic region with total copy number (CN) of 3. Red and orange horizontal segments denote sequencing reads. Both left and right mutations satisfy the a4s2 criteria because each has at least one DNA fragment (tagged by a UMI) with a4s2 coverage. For the right mutation, coverage for only the bottom UMI satisfies the a4s2 criterion, but the remaining DNA fragments where the variant is also present capture the information about multiplicity of the mutation in a cell.

- Variant supported by at least four reads, each with sequencing quality no less than 30
- All of the q$\geq$ 30 bases support the variant base.

Duplex sequencing technologies provide high-specificity mutation calls, at the expense of reduced sensitivity. Reduced sensitivity is a result of incomplete sequencing coverage, so that many variants do not have a4s2 (or similar) coverage. While non-duplex UMI-tagged DNA fragments do not have the duplex accuracy, they remain informative—particularly when estimating mutation multiplicity for confident variants—a4s2 duplex called on another DNA fragment, potentially in a different cell. For example, such non-duplex UMIs can be used to assess variant allele fraction (VAF) of mutations in data from many cells ("pseudobulk")

To quantify the prevalence of duplex-supported mutations within a given sample—defined as variant sites that pass the a4s2 criterion—we compute the ratio of duplex-supported mutations to all detected variants. This metric reflects the fraction of variant calls that meet stringent duplex criteria and can be leveraged for accurate estimates of somatic mutation burdens.

Below we describe two frameworks for correcting somatic burden by power to detect mutations in each cell. The "empirical" framework uses germline mutations and estimates their recall per cell and per copy number state. The "analytical" approach uses the observed UMI coverage per cell and the duplex ratio to calculate the power to detect somatic mutations in each cell and copy number state.

### 3 Empirical Framework - Estimating Mutation Detection Sensitivity Using Germline Mutations

Germline heterozygous mutations, identified from matched normal tissue, provide a useful reference for assessing copy number changes in allele-specific representation. Unlike somatic mutations, the multiplicity of germline heterozygotes is solely determined by local chromosomal copy number and, in the case of gains, by whether the paternal or maternal allele was amplified.

One approach to calculating power per multiplicity is by generating a pseudobulk using the UMI counts of germline heterozygous variants. These germline heterozgotes are called independently from bulk sequencing data. Since the initial copy number configuration of these variants is known—(T, M) = (2, 1)— and since these variants are numerous, we can accurately calculate their variant allele frequencies (VAFs), enabling assignment of allele-specific copy

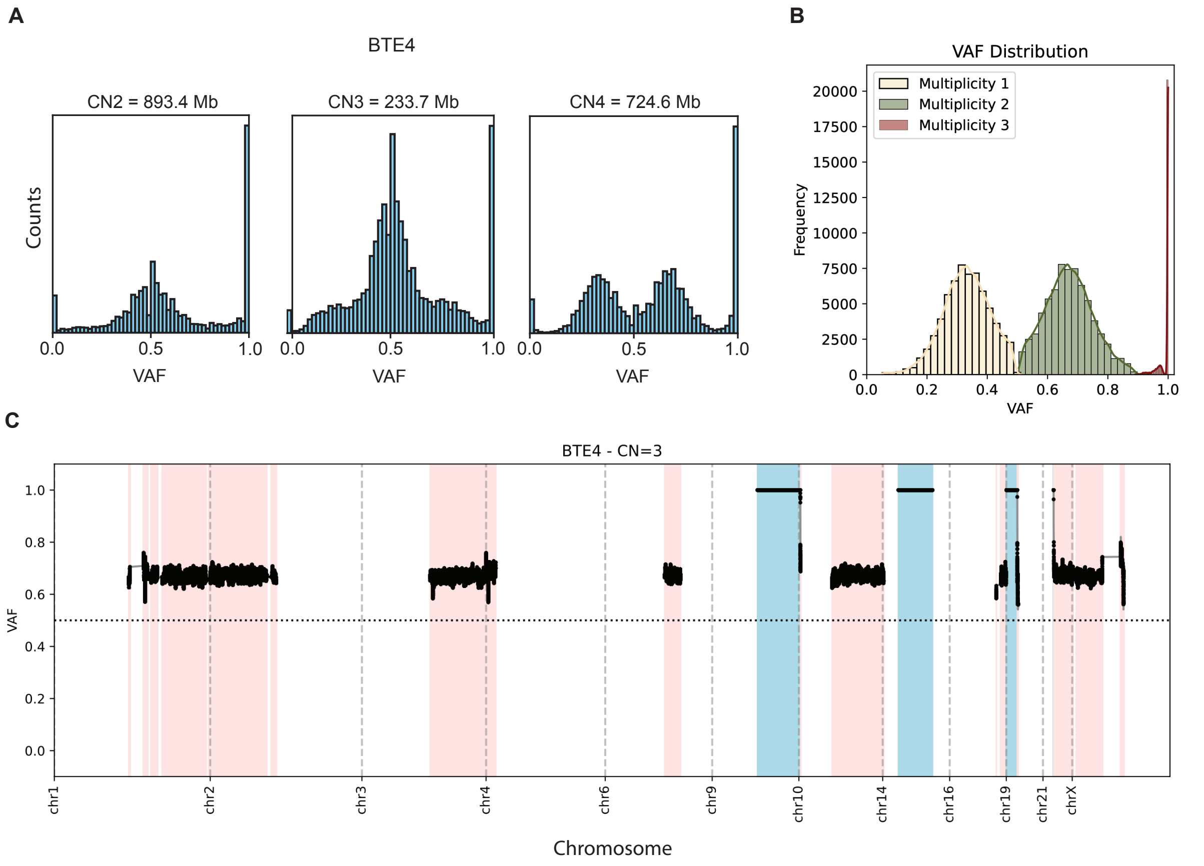


**Figure 2.** (a) Pseudobulk distribution of all variants—including non-duplex UMI-supported—stratified by total copy number (CN). The length of each corresponding genomic region is indicated in the panel title (in megabases, Mb). (b) Example VAF distribution for BTE4, CN=3 regions. Mutations assigned to different multiplicities are shown with different colours. (c) Allele-specific copy number ($M$) inference for regions with total copy number $T$, based on VAF profiles transformed to the range $[0.5, 1]$.

number states to genomic regions and classification of germline variants according to their most likely multiplicities.

Assuming that the power to detect somatic mutations can be approximated using germline heterozygous mutations within the same genomic regions, we can infer detection power per multiplicity by comparing the number of a4s2-called germline heterozygous mutations to the total number of germline mutations found from bulk whole-genome data in those regions. This provides an empirical measure of detection sensitivity under allele-specific copy number contexts. However, this approach has inherent limitations, which are discussed below.

To construct the pseudobulk, we restrict analysis to genomic regions where the total copy number ($T$) is consistent across cells. Specifically, we retain only regions in which the standard deviation of $T$ across cells is less than 0.5. Variants are then stratified according to total copy number, and their variant allele frequencies (VAFs) are computed and visualized, as shown for a representative sample in Fig. 2a.

The next step involves assigning allele-specific copy numbers and determining the minor allele copy number ($M$) for individual regions with a given $T$, based on the VAF profiles of germline mutations in these regions. To facilitate this, VAF values are first transformed to the range $[0.5, 1]$ and then smoothed to reduce noise. Genomic segments exhibiting consistent VAF behavior are grouped together, and the corresponding $M$ value is inferred. This procedure is illustrated in Fig. 2c, where regions with $(T, M) = (3, 0)$ are marked in turquoise and those with $(3, 1)$ in light pink.

The final step involves determining the multiplicity of each germline variant. To this end, we applied an Expectation-Maximization (EM) algorithm to fit a mixture of negative binomial models to the VAF distribution, as estimated from all UMIs. Each mutation was then assigned to a multiplicity class based on its VAF and the expected VAFs associated with different multiplicities. As shown in Fig. 2b, we illustrate an example for regions with total copy number $T = 3$, where mutations were classified into multiplicities $m = 1, 2, 3$, corresponding to theoretical VAFs of $1/3$, $2/3$, and 1, respectively.

Using this framework, we assigned multiplicities to duplex-supported germline heterozygous mutations (a4s2) by intersecting the set of variants present in the UMI-count pseudobulk (Fig. 2c) with the true detected variants. This enabled us to quantify, for each cell $i$, the number of detected germline heterozygous mutations in regions defined by total copy number $T$, minor allele copy number $M$, and assigned multiplicity $m$, denoted as $N_{T,M,m}(i)$. The final empirical power per multiplicity was then calculated as:

$$P^{\text{emp}}_{T,M,m}(i) = \frac{N_{T,M,m}(i)}{\alpha B_{T,M}}, \tag{1}$$

where $B_{T,M}$ denotes the total number of germline heterozygous mutations in chromosomal regions with total copy number T and minor M, detected in a bulk sample from the matched normal tissue, irrespectively of whether the mutations were also detected in the pseudobulk. Given that the matched normal genome has an initial copy number configuration of $(T,M) = (2,1)$—with mutations present on both chromosomal copies—the inclusion of the $\alpha$ term serves to normalize the expected size of the matched normal mutation set. This correction ensures comparability with the germline heterozygous mutation sets detected in tumor cells, where copy number alterations may shift the underlying multiplicity distributions.

To illustrate the role of the $\alpha$ term, consider a genomic region with total copy number $T = 3$. In this context, the possible minor allele copy numbers are $M = 0,1$:

- For $M = 1$, the possible multiplicities for a mutation are $m = 1,2$. Assuming that half of the mutations in $(T,M) = (3,1)$ have $m = 1$ and the other half have $m = 2$, $\alpha$ is set to 0.5 to ensure a fair comparison with the equivalent matched normal bulk set in such regions.
- For $M = 0$, there is only one possible minor CN ($M = 0$) with a multiplicity of $m = 3$. In this case, to obtain a correct empirical power estimate, $\alpha$ is still set to 0.5. This accounts for the fact that one parental copy (and thus half of the germline mutations in the matched normal bulk) is entirely lost.

Equivalent corrections can be performed for any total CN region.

#### 3.1 Limitations

The empirical framework described above relies on several assumptions that can significantly influence the results and limit its applicability in certain scenarios. It is primarily suited for cases where the total copy number (CN) profile is relatively homogeneous across cells within the same sample. When CN profiles vary significantly between cells, it becomes difficult to confidently define a pseudobulk and defined regions with fixed total CN, which can lead to inaccurate VAF calculations and affect downstream analyses.

In addition, the presence of variable CN states between cells reduces the number of genomic regions that can be reliably assigned a consistent total CN value across cells. As a result, only a small fraction of germline mutations is retained, making the correction applicable to a limited number of regions. Even in cases where the VAF spectrum is reasonably well defined, such as in Fig. 2a, there can still be variability in identifying the VAF peaks. Outlier cells may introduce further noise, making it challenging to accurately assign multiplicities to mutations.

### 4 Analytical Framework

Applying our empirical framework to samples with relatively homogeneous total copy number (CN) profiles across cells revealed that the distribution of total variant counts (UMI counts) across multiplicity states encodes informative structure about mutation multiplicity. For example, in regions with total CN = 3 containing germline heterozygous variants, we were able to decompose the VAF distribution into distinct multiplicity components, owing to the framework's ability to reliably assign multiplicity to mutations. UMI support for germline variants with multiplicity estimates for a representative cell is shown in Fig. 3a. The zero bin reflects variants confidently identified and assigned multiplicity in the pseudobulk but lacking detectable support in that specific cell. Although this approach cannot be directly applied to somatic mutations (Fig. 3a) due to the lack of prior multiplicity estimates, it motivated the development of an analytical model to infer the true number of somatic mutations per multiplicity and CN state by exploiting the structure of the observed UMI count distributions.

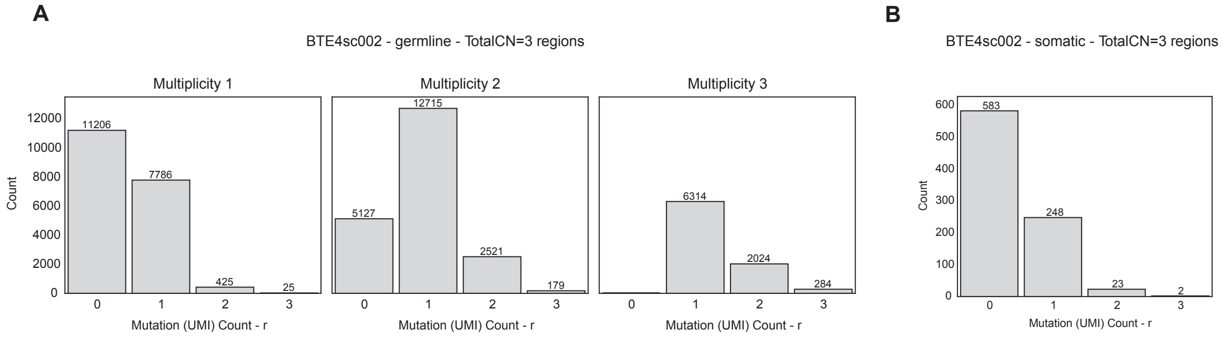


**Figure 3.** A. UMI support barplot for germline variants in a representative cell. The germline variants are stratified by mutation multiplicity estimated from pseudobulk across all cells. B. UMI support barplot for somatic mutations from the same cell. Our analytical method decomposes the observed distribution of UMI support for somatic mutations into a mixture of distributions, one per multiplicity.

The characteristic property of Duplex sequencing assays that allows for such a framework is that each DNA fragment, originating from a chromosomal copy is assigned to a unique molecular identifier (UMI). This implies that per cell, in a region of total CN, $T$, the number of UMI counts (variants) reported at a locus, which from now on will be denoted as $r$, can not exceed the number of available copies:

$$r \leq T. \tag{2}$$

Unlike other sequencing techniques (not using UMIs and not single cell) where the data we get is the raw number of reads covering a given locus, and hence we have no control over how many reads report a chromosomal copy, in single-cell duplex sequencing data we know that the number of UMIs reporting a variant is limited by the number of chromosome copies in a cell. Sampling DNA fragments at a locus should be modeled without replacement—the same DNA fragment cannot be sequenced twice. At a given locus with total CN $T$, assuming that a mutation is present with multiplicity $m$ (with $m \leq T$) and that the total number of available UMIs at that locus is $n$, the probability that $r$ UMIs will report a **variant** is given by the Hypergeometric distribution—a distribution that models sampling without replacement:

$$P(r|T,n,m) = \frac{\binom{m}{r}\binom{T-m}{n-r}}{\binom{T}{n}} \tag{3}$$

To call a mutation in scNanoSeq data, a variant must meet the **a4s2** criterion. We model the probability of a variant being duplex-called as a mutation using a binomial process. The probability of success ($p_d$) in this process is estimated empirically as a fraction of the number of duplex-called germline mutations (A; duplicated, if a mutation is duplex-detected on multiple UMIs) over all total UMI support of germline mutations (B; total UMI support for all germline variants; a UMI is counted twice if it supports two variants) available UMIs reporting germline variant in a given sample/cell.

$$p_d = \frac{A}{B} \tag{4}$$

The probability that a mutation would be duplex-called is calculated as a probability that at least one of these $r$ DNA fragments tagged by UMIs will meet the a4s2 criteria ($r_d \geq 0$) and is hence given by:

$$P(r_d \geq 0) = \sum_{r_d=1}^{r} \text{Bin}(r_d, p_d) = 1 - \text{Bin}(r_d = 0, r, p_d) \tag{5}$$

In the presented formula in Eq. 3, $n$ is a discrete variable that can take integer values from 1 to $T$. It becomes apparent that in order to apply the proposed analytical framework to observed data, we need to have a measure of

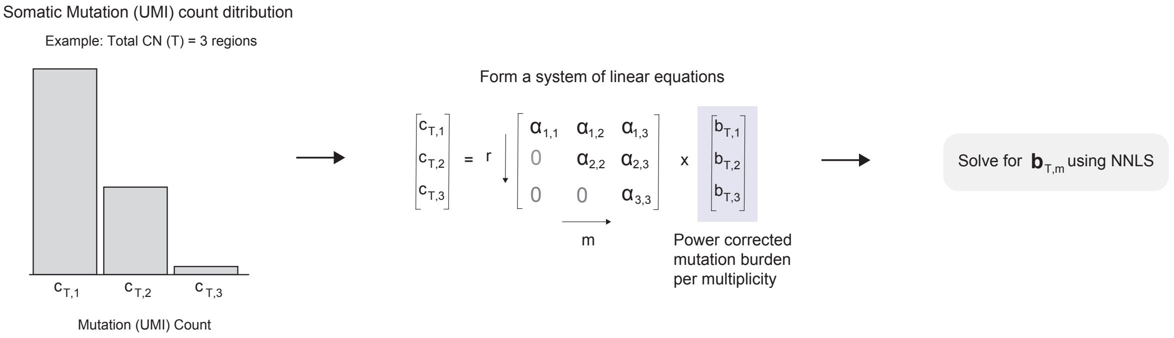


**Figure 4.** Schematic showing how the observed UMI support of somatic mutations can be used to estimate the power-corrected burden of somatic mutations.

the distribution of total UMI coverage, $n$, for a cell, in a total CN region, $T$. To be able to use this term we should therefore weigh each $n$ component with the observed UMI coverage.

$$\langle P(r \mid T,n,m)\rangle_{n_{obs}} = w_0 \times 0 + w_1 P(r|T,n=1,m) + \cdots + w_T P(r|T,n=T,m), \tag{6}$$

where each weight is calculated by the observed UMI count distribution in each cell as follows: $w_i = \frac{w_i}{\sum_i w_i}$ Hence, at a region with total CN $T$, UMI count distribution $n_{obs}$, having a mutation with true multiplicity $m$, the probability that $r$ UMIs will report a variant ($r \leq m$) and at least one of these variants will be classified as a duplex mutation is given by:

$$\langle P(r \mid T,n,m)\rangle_{n_{obs}} \times (1 - \mathrm{Bin}(r_d = 0, p_d)) \tag{7}$$

The above formula allows us to analytically estimate the sensitivity for duplex mutation calls given the total copy number and mutation mutation multiplicity. We estimate this distribution ($n_{obs}$) using UMI coverage at known germline mutations, and the fraction of UMIs satisfying the duplex criteria, which can be readily estimated for each cell.

#### 4.1 Estimating $n_{obs}$ distribution

To estimate the weights $w_i$ described above, including the fraction of mutations missed due to incomplete coverage, we leveraged germline heterozygous mutations and the matched normal bulk sequencing data. By stratifying germline variants detected in the bulk normal by total CN (as inferred from the corresponding tumor sample), restricting to confidently assigned CN regions, and merging with the single-cell UMI counts in those same regions, we derived a per-cell empirical distribution of UMI support across multiplicity states. This includes an estimate of $w_0$, reflecting the fraction of mutations not covered in scNanoSeq. Assuming that the coverage profile of germline heterozygous variants is representative, we used this distribution to extrapolate to somatic mutations.

#### 4.2 Defining power per multiplicity

From the above equation, it is evident that mutations with higher multiplicity are more likely to be called using duplex criteria, as they may be present and sequenced from more DNA fragments. Given the constraints on the total UMI count $n$ and true multiplicity $m$, we can construct a triangular matrix based on the value of T. The coefficients of this matrix are given by:

$$\alpha_{rm}(T, p_d, n_{obs}) = \langle P(r \mid T,n,m)\rangle_{n_{obs}} \times (1 - \mathrm{Bin}(r_d = 0, r, p_d)), \quad n,m \leq T \quad r \leq m \tag{8}$$

The goal of the proposed analytical power framework is to estimate the true number of somatic mutations per cell, power corrected and stratified by multiplicity. To do so, the framework uses the observed somatic UMI counts with $r \geq 1$ (as illustrated in Fig. 3b), together with the inferred coefficients of the power matrix $\alpha_{rm}$, to solve a system of linear equations (schematized in Fig. 4). This computation is performed on a per-cell basis, and the system is solved using non-negative least squares (NNLS).

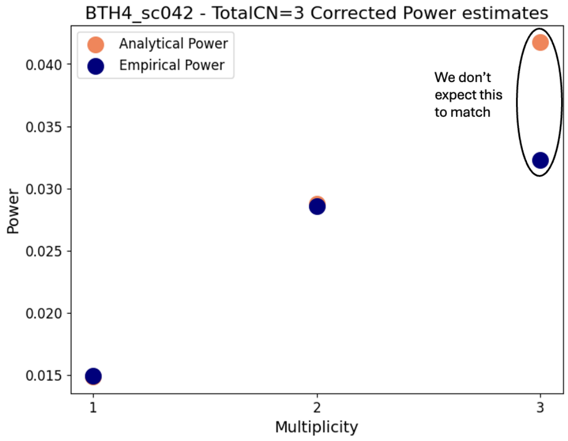


**Figure 5.** The analytical and emprical estimates of mutation detection sensitivity (power) in an example cell for regions with $T = 3$. The analytical and empirical estimates match for the mutations with multiplicity of 1 and 2, but mismatch for multiplicity of 3. We believe the discrepancy can be attributed to the assumptions used for the empirical estimates.

### 5 Comparison of Analytical and Empirical Estimates

To validate the analytical estimate of power per multiplicity for a given cell $i$ within a sample, we compare it against confident empirical estimates from germline mutations. The analytical power per multiplicity is formally defined as:

$$\mathrm{Pow}^{\mathrm{an}}_{T,m}(i, n_{\mathrm{obs}}, p_d) = \sum_r \alpha_{rm}(T, p_d, n_{\mathrm{obs}}),$$

where $\mathrm{Pow}^{\mathrm{an}}_{T,m}(i, n_{\mathrm{obs}}, p_d)$ denotes the estimated power to detect mutations for total copy number $T$ and multiplicity $m$ in cell $i$. We validated our analytical estimates using a sample characterized by a high number of mutations in regions with retained total CN = 3, specifically regions with CN configuration $(T, M) = (3, 1)$. As demonstrated for this sample and consistently observed across all samples, the power estimates derived empirically closely agree with those from our analytical method when sufficient germline mutation counts are present in the retained regions, underscoring the accuracy of the proposed framework.

Notably, for the configuration $(T, M) = (3, 0)$, the empirical estimate of mutation detection power does not match the analytical one. We attribute the mismatch to the breakdown of the assumption underlying the $\alpha$ parameter (Eq. 1) when one parental chromosomal copy is completely lost in the single cells. If chromosomal copy one from one of the parents carries more or fewer germline variants, $\alpha = 0.5$ is not appropriate. Since phasing of germline variants is not possible, there is no way to a priori estimate $\alpha$ suitable for a cell and a given segment of loss-of-heterozygosity (LOH). The problem may be more acute when there is only a handful of LOH segments, as in the discussed case.

Because the parameters needed for the empirical model are not evaluable, we opted for using the analytical model for correction of mutation detection sensitivity in the manuscript. Furthermore, in other samples and cells, the calculations for the empirical model are not feasible because of the copy number being too variable between cells.
