## Supplementary material for "Timing the onset of homologous recombination deficiency before breast cancer diagnosis": Supplementary Note 2 - Gradual SBS1 Acceleration Model.pdf

### Gradual SBS1 Acceleration Model

Michail Andreopoulos

SBS1 mutations arise from the spontaneous deamination of methylated cytosines and are widely regarded as accumulating approximately linearly with chronological age in normal tissues. In tumor samples, proliferative dynamics are expected to perturb this baseline process, leading to an acceleration of the SBS1 mutation rate relative to normal tissue, as reflected in the elevated SBS1 burden observed at diagnosis.

Here, we develop a model for SBS1 accumulation as a function of age that explicitly accounts for sample-specific characteristics, including cancer subtype and age at diagnosis. The aim is to infer the temporal dynamics of SBS1 rate changes within tumors by jointly modeling baseline accumulation and subsequent acceleration from the observed mutational burden.

#### 1 Baseline SBS1 Accumulation in Normal Tissue

##### 1.1 Data

The baseline SBS1 accumulation rate is estimated from EpCAM-positive cells isolated from histologically normal mammary tissue in donors lacking identifiable driver mutations. For each normal-tissue sample of age $x$ with observed SBS1 burden $y$ (mutations per megabase, normalized to a 3000 Mb diploid genome), the cell-specific rate is $y/x$.

##### 1.2 Model

Assuming linear SBS1 accumulation with age in normal tissue, the baseline trajectory is:

$$y_{\text{norm}}(x) = s_{\text{norm}} \cdot x, \tag{1}$$

where $s_{\text{norm}}$ is sampled from the empirical distribution of $y/x$ across all normal-tissue samples (see Section 8).

##### 1.3 Sampling $s_{\text{norm}}$

A kernel density estimate (KDE) is constructed using $y/x$ ratios observed in normal cells, using a Gaussian kernel with bandwidth $h = 0.5$, giving the density $\hat{f}_{\text{KDE}}$. In each iteration, $s_{\text{norm}}$ is drawn from this distribution by inverse-CDF sampling. The 5th and 95th percentiles $s_{\text{low}}$, $s_{\text{high}}$ are displayed as a grey band in the cumulative burden plots. The normal-SBS1 slope, $s_{\text{norm}}$, may vary as a function of the cell type from which the tumor-initiating precursor arose, as well as the presence of early driver events that elevate the total SBS1 burden during initial stages. Sampling from the kernel density estimate (KDE) explicitly propagates this variability into the final estimate of onset age, accounting for a range of such scenarios given the observed distribution of SBS1 burden in normal cells.

#### 2 Observed Tumor Data

For each tumor sample, the inputs are:

- $A$: age at diagnosis (years);
- $Y$: total SBS1 burden per diploid genome at diagnosis;
- Tumor subtype: ER+ or triple-negative (TN);
- $f_{\text{HRD}} \in [0,1]$: HRDTime: the fraction of the total SBS1 burden (scaled to the diploid genome) that had accumulated by the time the HRD event occurred, reported with a 95% confidence interval $[\ell, u]$.

The HRDTime ($f_{\text{HRD}}$) is relevant only in the final step (Section 7), where it is used to estimate the age of HRD onset using the inferred SBS1-to-age model. The SBS1 burden at the HRD event is:

$$Y^* = f_{\text{HRD}} \cdot Y. \tag{2}$$

Uncertainty in $f_{\text{HRD}}$ is propagated by sampling $f_{\text{HRD}} \sim \mathcal{N}(\mu_{\text{HRD}}, \sigma_{\text{HRD}})$, where $\sigma_{\text{HRD}} = (u - \ell)/(2 \times 1.96)$.

#### 3 Subtype-Specific Tumor Growth Period Priors

The time between tumor initiation and clinical diagnosis is not directly observable and cannot be inferred in the absence of sample-specific longitudinal data. In our model, we estimate the tumor growth period using subtype-specific data on tumor growth rates established from consecutive mammography scans[1]. ER+ tumors are generally slower-growing, whereas triple-negative (TN) tumors are more aggressive and develop faster. These differences are reflected in longer predicted tumor growth period for ER+ versus TN tumors.

##### 3.1 Prior distributions

Tumor growth period, $d$ (years), is sampled each iteration as:

$$d \sim \mathcal{N}(\mu_d, \sigma_d), \tag{3}$$

where $\sigma_d = (d_{\text{hi}} - d_{\text{lo}})/(2 \times 1.96)$ from the published[1] 95% confidence interval $[d_{\text{lo}}, d_{\text{hi}}]$:

| Subtype | $\mu_d$ (yr) | 95% CI (yr) | $\sigma_d$ (yr) |
|---|---|---|---|
| ER+ | 16.33 | [13.55, 19.10] | 1.42 |
| TN | 10.36 | [6.60, 14.32] | 1.97 |

##### 3.2 Late-phase SBS1 rate

Given $s_{\text{norm}}$ and $d$, the SBS1 rate during the tumor phase —the interval from tumor initiation to diagnosis—must yield the observed total burden $Y$. Let $x_d = A - d$ denote the inferred age at tumor initiation, at which the normal-tissue trajectory has reached a burden $s_{\text{norm}} x_d$. If the SBS1 rate during the tumor phase is constant on average, the tumor-phase slope is the slope of the line joining initiation, $(x_d,\ s_{\text{norm}}x_d)$, to diagnosis, $(A,\ Y)$:

$$s_{\text{tumor}} = \frac{Y - s_{\text{norm}} \cdot x_d}{d}, \qquad x_d = A - d. \tag{4}$$

The fold-change $s_{\text{tumor}}/s_{\text{norm}}$ quantifies the increased relative rate of SBS1 accumulation during the tumor phase, with values of 5–30-fold typical across the samples analyzed; the upper bound of this range is set by the analysis of metastatic breast cancers (Figure 3C).

#### 4 Gradual SBS1 Acceleration Model

A step-change model (two-step model) — in which the SBS1 rate increases instantaneously from $s_{\text{norm}}$ to $s_{\text{tumor}}$ at tumor initiation — is biologically unrealistic. Nevertheless, it provides a useful boundary case corresponding to maximal acceleration, and thus defines a lower bound on the inferred tumor growth period. In reality, cell fitness increases progressively as multiple oncogenic mutations are acquired. We therefore model the transition in rate using a smooth sigmoid function, which more realistically captures the gradual nature of this process.

##### 4.1 Instantaneous rate

We model the instantaneous SBS1 accumulation rate at age $x$ is:

$$\frac{dy}{dx}(x) = s_{\text{norm}} + r_d \cdot \sigma(x; m, k), \qquad \sigma(x; m, k) = \frac{1}{1 + e^{-k(x-m)}}, \tag{5}$$

where:

- $r_d = s_{\text{tumor}} - s_{\text{norm}} > 0$ is the excess SBS1 rate attributable to the tumor-phase acceleration;
- $m$ (years) is the sigmoid midpoint — the age at which the instantaneous rate is halfway between $s_{\text{norm}}$ and $s_{\text{tumor}}$;
- $k > 0$ (yr$^{-1}$) controls the steepness of the transition.

At early ages ($x \ll m$), $\sigma \approx 0$ and the rate is close to $s_{\text{norm}}$: the sample follows the SBS1-accumulation rate defined by the normal cells and behaves like normal tissue. At late ages ($x \gg m$), $\sigma \approx 1$ and the rate approaches $s_{\text{tumor}}$: the tumor-phase acceleration is fully established.

##### 4.2 Cumulative burden

The cumulative SBS1 burden at age $x$ is obtained by integrating the instantaneous rate ((5)) from the time of fertilization:

$$y(x) = \int_0^x \left[s_{\text{norm}} + r_d \cdot \sigma(v; m, k)\right] dv. \tag{6}$$

The antiderivative of the sigmoid is the softplus function:

$$\int \sigma(v; m, k)\, dv = \frac{1}{k} \ln\left(1 + e^{k(v-m)}\right) + C, \tag{7}$$

so evaluating from 0 to $x$ gives:

$$\int_0^x \sigma(v; m, k)\, dv = \frac{1}{k} \ln\left(1 + e^{k(x-m)}\right) - \frac{1}{k} \ln\left(1 + e^{-km}\right). \tag{8}$$

The second term is the antiderivative evaluated at $v = 0$, which ensures $y(0) = 0$. Combining both terms:

$$y(x) = s_{\text{norm}} \cdot x \;+\; r_d \left[\frac{1}{k} \ln\left(1 + e^{k(x-m)}\right) - \frac{1}{k} \ln\left(1 + e^{-km}\right)\right]. \tag{9}$$

The $r_d$ term is zero at $x = 0$ and grows smoothly from baseline, producing the characteristic S-shaped rise in cumulative SBS1 burden visible in the trajectory plots.

#### 5 Probabilistic Sampling of the Steepness $k$

Rather than treating $k$ as fixed, we allow it to vary across iterations to capture a spectrum of biologically plausible SBS1 acceleration regimes. Specifically, $k$ is sampled at each iteration from a distribution determined by model-informed quantities—namely, the tumor growth period and the relative SBS1 excess (defined as the SBS1 burden at diagnosis relative to that expected under baseline SBS1 accumulation). This formulation reflects uncertainty in the rate at which tumors transition from baseline to accelerated SBS1 accumulation.

##### 5.1 The logit function and the tumor growth period constraint on $k$

To relate the steepness $k$ of the sigmoid to a biologically interpretable *timescale*, we make use of the inverse relationship between the sigmoid and the logit function. The *logit* of a quantity $f \in (0,1)$ is defined as

$$\text{logit}(f) = \ln\left(\frac{f}{1-f}\right), \tag{10}$$

where $f$ represents the *fraction of transition* from the early (baseline) SBS1 rate to the late (accelerated) rate.

For the sigmoid $\sigma(x;m,k)$, the value at a displacement $\Delta$ from the midpoint $m$ corresponds to the fraction of the transition that has been completed:

$$\sigma(m+\Delta;m,k) = \frac{1}{1+e^{-k\Delta}}. \tag{11}$$

Solving $\sigma(m+\Delta;m,k) = f$ gives

$$\Delta = \frac{\text{logit}(f)}{k}. \tag{12}$$

This quantity $\Delta$ represents the time required to progress from the midpoint (50% completion) to a higher completion level $f$ (e.g. 90%).

Because the sigmoid is symmetric about $m$, the corresponding point on the left satisfies $\sigma(m-\Delta) = 1-f$. Therefore, the full duration required to transition from $(1-f)$ to $f$—for example, from 10% to 90% completion—is

$$\Delta t_{(1-f)\to f} = \frac{2\,\text{logit}(f)}{k}. \tag{13}$$

This provides a direct mapping between the steepness $k$ and the total transition time.

###### 5.1.1 Choice of boundaries on $k$

In the absence of longitudinal data, we define a conservative, biologically grounded range $[k_{\min}, k_{\max}]$ for the steepness $k$, using the mapping between $k$ and the transition timescale established above (with $f = 0.90$).

**Lower bound.** The full $10\% \to 90\%$ transition cannot take longer than the tumor growth period $d$, which we treat as an upper bound on the transition window:

$$\frac{2\,\text{logit}(f)}{k} \le d \qquad \Longrightarrow \qquad k_{\min} = \frac{2\,\text{logit}(f)}{d}. \tag{14}$$

Intuitively, slower-growing tumors can transition more gradually (longer $d$ permits smaller $k$), whereas shorter durations force steeper transitions to complete the shift in the time available.

**Upper bound.** We impose a minimum plausible transition duration $\Delta t_{\min} = 3$ years, representing the most rapid biologically reasonable acceleration. This caps the steepness at

$$k_{\max} = \frac{2\,\text{logit}(f)}{\Delta t_{\min}}. \tag{15}$$

Together these define the admissible range $k \in [k_{\min}, k_{\max}]$.

##### 5.2 Sampling of $k$ per iteration

In each iteration, the baseline rate $s_{\text{norm}}$ and tumor growth period $d$ are sampled first. Together with $\Delta t_{\min}$, the sampled $d$ fixes the admissible range $k \in [k_{\min}, k_{\max}]$ for that iteration via Eq. (14).

Where $k$ falls within this range is not chosen uniformly but guided by two data-derived quantities, defined below: a duration component $p_{\text{dur}}$ and a relative-excess component $p_{\text{excess}}$. These are not direct predictors of $k$; rather, they modulate its position within the admissible range. The two are combined into a single score that interpolates between $k_{\min}$ and $k_{\max}$.

###### 5.2.1 Duration component: $p_{\mathrm{dur}}$

The sampled tumor growth period $d$ is mapped to its percentile under the subtype-specific prior:

$$p_{\mathrm{dur}} = \Phi\left(\frac{d-\mu_d}{\sigma_d}\right), \tag{16}$$

where $\Phi(\cdot)$ is the standard normal CDF. Thus $p_{\mathrm{dur}} \approx 0.5$ for a typical period and $p_{\mathrm{dur}} \ll 0.5$ for an unusually short one. We use $(1-p_{\mathrm{dur}})$ as the weight, which is large only when the sampled period is atypically short.

###### 5.2.2 Relative excess component: $p_{\mathrm{excess}}$

We quantify the strength of SBS1 acceleration as the fold-change between the tumor and baseline rates, capped and normalized to the unit interval:

$$p_{\mathrm{excess}} = \frac{1}{R}\min\left(\frac{s_{\mathrm{tumor}}-s_{\mathrm{norm}}}{s_{\mathrm{norm}}},\, R\right), \qquad R = 30. \tag{17}$$

When $s_{\mathrm{tumor}} \approx s_{\mathrm{norm}}$ the increase is small and $p_{\mathrm{excess}} \approx 0$; large fold-changes drive $p_{\mathrm{excess}} \to 1$ and favor steeper transitions. The cap $R$, inferred from metastasis data, prevents extreme fold-changes from dominating.

###### 5.2.3 Combined steepness prior

The two components are combined multiplicatively:

$$p_{\mathrm{combined}} = (1-p_{\mathrm{dur}})\cdot p_{\mathrm{excess}} \;\in\; [0,1]. \tag{18}$$

The multiplicative form reflects a biological assumption: a steep transition is favored only when the tumor growth period is unusually short *and* the SBS1 excess is large. If either signal is weak, $p_{\mathrm{combined}}$ stays small and more gradual transitions are preferred.

###### 5.2.4 Sampling of $k$

The combined score interpolates within the admissible range:

$$k_{\mathrm{center}} = k_{\mathrm{min}} + p_{\mathrm{combined}}\left(k_{\mathrm{max}} - k_{\mathrm{min}}\right), \tag{19}$$

so $p_{\mathrm{combined}} = 0$ gives the shallowest transition ($k_{\mathrm{min}}$) and $p_{\mathrm{combined}} = 1$ the steepest ($k_{\mathrm{max}}$). Rather than fixing $k$ at this central value, we sample around it to avoid over-committing to a single point in the admissible range. We take a modest spread, set to a small fraction of the range width,

$$\sigma_k = 0.1\left(k_{\mathrm{max}} - k_{\mathrm{min}}\right), \tag{20}$$

so that sampled values stay well within $[k_{\mathrm{min}}, k_{\mathrm{max}}]$, and draw

$$k \sim \mathcal{N}(k_{\mathrm{center}},\, \sigma_k). \tag{21}$$

This lets $k$ vary smoothly across plausible regimes while staying anchored to both the temporal constraints and the observed magnitude of SBS1 acceleration.

#### 6 Constrained Optimization of the Transition Midpoint

For each sampled parameter set $(s_{\mathrm{norm}}, d, k)$, the sigmoid midpoint $m$ is inferred rather than fixed. It is the only free parameter of the fit: the excess rate is set in advance from the sampled slopes, $r_d = s_{\mathrm{tumor}} - s_{\mathrm{norm}}$ (Eq. (5)), so $m$ alone positions the transition in time such that both the total SBS1 burden and the late-phase rate are consistent with the observed data.

##### 6.1 Constraint formulation

The midpoint $\hat{m}$ is obtained by solving a least-squares system defined by two constraints:

$$y(A;\, \hat{m}) = Y, \tag{22}$$

$$\left.\frac{dy}{dx}\right|_{x=0.95A,\, \hat{m}} = s_{\mathrm{tumor}}. \tag{23}$$

**Burden consistency (Eq. (22)).** The cumulative SBS1 burden at diagnosis, evaluated from Eq. (9), must match the observed value $Y$.

**Late-rate constraint (Eq. (23)).** The instantaneous SBS1 rate near diagnosis (at $0.95A$), evaluated from Eq. (5), must match $s_{\text{tumor}}$. Because the late asymptote $s_{\text{norm}} + r_d = s_{\text{tumor}}$ holds by construction, this constraint forces the sigmoid to be near saturation by $0.95A$, i.e. the accelerated regime must be essentially complete by diagnosis.

##### 6.2 Solution

The two constraints are assembled into a residual vector

$$\mathbf{R}(m) = \begin{pmatrix} y(A;m) - Y \\ \frac{dy}{dx}\Big|_{x=0.95A,\, m} - s_{\text{tumor}} \end{pmatrix}, \tag{24}$$

and $\hat{m}$ is obtained by least-squares minimization of $\|\mathbf{R}(m)\|^2$, constrained to $m \in [0, A]$ and initialized at $m_0 = A/2$. A fit is accepted only if the residual cost falls below $10^{-3}$.

Steeper transitions (larger $k$) place $\hat{m}$ closer to diagnosis, while shallower ones shift it earlier. Because $k$ is sampled within a biologically constrained range (Section 5), the optimization converges reliably and yields no unrealistic midpoints in practice.

#### 7 Locating the HRD Event on the SBS1 Trajectory

Given a fully specified SBS1 accumulation trajectory $y(x)$ determined by $(s_{\text{norm}}, d, k, \hat{m})$, the HRD event is inferred as the time at which the trajectory accumulates a fraction of the total observed SBS1 burden consistent with molecular clock estimates.

We define a target burden threshold:

$$Y^* = f_{\text{HRD}} \cdot Y, \tag{25}$$

and solve for the crossing time:

$$a_c = \{x \in [0, A] \; : \; y(x; \hat{m}, k, s_{\text{norm}}) = Y^*\}. \tag{26}$$

This equation is solved numerically for each sampled trajectory. Lower and upper uncertainty bounds, $a_l$ and $a_h$, are computed analogously by perturbing $f_{\text{HRD}}$ within its uncertainty interval.

The resulting estimate $a_c$ represents the inferred absolute age at which the tumor accumulated the SBS1 burden attributable to HRD onset under the fitted trajectory. The implied HRD duration is $A - a_c$. Importantly, the transformation to calendar years is applied post-hoc: the SBS1-based estimate of HRD onset is projected onto the inferred SBS1-to-age function without influencing the estimation of its parameters: $m$ or $k$.

#### 8 Probabilistic Uncertainty Propagation

All model inputs carry uncertainty, including the baseline slope $s_{\text{norm}}$, tumor growth period $d$, HRD fraction $f_{\text{HRD}}$, and the sigmoid steepness $k$. These uncertainties are propagated jointly through the nonlinear SBS1 accumulation model to obtain a distribution over HRD onset times.

##### 8.1 Algorithm

For each tumor sample, $N = 1000$ valid iterations are generated:

1. **Sample baseline slope.** $s_{\text{norm}} \sim \hat{f}_{\text{KDE}}$ via inverse-CDF sampling.
2. **Sample tumor growth period.** $d \sim \mathcal{N}(\mu_d, \sigma_d)$ from the subtype-specific prior.
3. **Compute late-phase slope.** $s_{\text{tumor}}$ is computed via Eq. (4).
4. **Sample HRD fraction.** $f_{\text{HRD}} \sim \mathcal{N}(\mu_{\text{HRD}}, \sigma_{\text{HRD}})$, defining $Y^* = f_{\text{HRD}} \cdot Y$.
5. **Compute $p_{\textbf{combined}}$ and sample $k$.** Using $(d, s_{\text{tumor}}, s_{\text{norm}})$, compute $p_{\text{dur}}$, $p_{\text{excess}}$, and $p_{\text{combined}}$, and sample $k \sim \mathcal{N}(k_{\text{center}}, \sigma_k)$. (Section 5).

6. **Fit midpoint.** Solve the constrained system (Section 6) for $\hat{m}$.

7. **Infer HRD crossing time.** Solve $y(a_c) = Y^*$.

##### 8.2 Output summaries

For each tumor, posterior summaries are computed over all accepted iterations:

- median and 5–95% quantiles of $a_c$ (HRD onset age),
- median and 5–95% quantiles of $A - a_c$ (HRD duration).

We additionally report the empirical distributions of $k$, $p_{\text{combined}}$, $d$, and $s_{\text{tumor}}$ as diagnostic summaries of model behavior.

#### 9 Parameter Summary

| Symbol | Description | Value / Distribution | Origin |
|---|---|---|---|
| $s_{\text{norm}}$ | Baseline SBS1 slope | $\hat{f}_{\text{KDE}}$ | Normal tissue data |
| $d$ | Tumor growth period | $\mathcal{N}(\mu_d, \sigma_d)$ | Subtype prior |
| $f_{\text{HRD}}$ | HRD molecular fraction | $\mathcal{N}(\mu_{\text{HRD}}, \sigma_{\text{HRD}})$ | Sample-specific |
| $k$ | Sigmoid steepness | $\mathcal{N}(k_{\text{center}}, \sigma_k)$, truncated | Derived (Section 5) |
| $m$ | Sigmoid midpoint | Optimized latent variable | Constraint system |
| $f$ | Transition threshold | 0.90 | Fixed |
| $\Delta t_{\text{min}}$ | Minimum transition time | 3 yr | Biological prior |
| $R$ | Fold-change saturation | 30 | Fixed |
| $N$ | Iterations per sample | 1000 | Fixed |

#### References

[1] Emma G MacInnes et al. “Radiological audit of interval breast cancers: Estimation of tumour growth rates”. In: *The Breast* 51 (2020), pp. 114–119.
