## Supplementary material for "Timing the onset of homologous recombination deficiency before breast cancer diagnosis": Supplementary Note 3 - Sensitivity of age-at-HRD estimates to post-HRD SBS3 rate assumptions.pdf

#### Overview

HRDTimer estimates the timing of homologous recombination deficiency (HRD) onset as a fraction $t_{\text{HRD}} \in [0,1]$ on the SBS1-based molecular-time axis. The framework assumes that, following HRD onset, SBS3 acts as a secondary molecular clock whose rate is sample-specific but, on average, constant over time—an assumption supported by the stability of SBS3 accumulation we observe across the late-clonal, subclonal, and cell-private periods (**Supp. Fig. X**).

To probe alternative modes of SBS3 accumulation and quantify the sensitivity of our estimates to this assumption, we developed a simulation framework that randomly draws each tumour's mutation catalogue under four biologically motivated SBS3-rate scenarios while preserving all other properties of the real tumour: its copy-number structure, total mutation burden, and the phase-specific (early, late, undefined) mixture of non-SBS3 signatures. For each scenario and sample, 200 bootstrap replicates were passed through the complete HRDTimer–age-translation algorithm, and the resulting age-at-HRD estimates were compared with a reference age derived directly from the sample's HRDTimer posterior. We applied the simulation framework to the $n = 20$ PCAWG breast cancer samples that passed all QC checks and were included in the main analysis (see Main text).

#### Notation

| Symbol | Meaning |
|---|---|
| **Indices & sizes** | |
| $i$ | Index over somatic mutations (catalogue rows) |
| $N$ | Number of somatic mutations |
| $N_{\text{SBS3}}$ | SBS3 mutation count, $N_{\text{SBS3}} = \sum_i p_i^{\text{SBS3, real}}$ |
| **Fixed tumour inputs** | |
| $\text{MajCN}_i$, $\text{MinCN}_i$ | Major/minor copy number at the locus of mutation $i$ |
| $\phi(i)$ | Temporal phase of mutation $i$ relative to WGD: **Early** (pre-WGD), **Late** (post-WGD), or **NA** (undefined; in (2,1) regions) |
| $p_i^G$, $p_i^S$ | Soft pre-WGD (Gain) / post-WGD (Single) probabilities for NA mutations |
| $p_i^{\text{SBS3, real}}$ | MuSiCal posterior probability that mutation $i$ is SBS3 (real-tumour fit; unperturbed data) |
| $\mathbf{m}_\phi$ | Real-tumour exposure vector for phase $\phi$ (non-SBS3 signatures, unit sum) |
| $t_{\text{HRD}}$ | HRD onset in SBS1-based molecular time calculated from unperturbed data; SBS3 active on $[t_{\text{HRD}}, 1]$ |
| $\omega$ | Post-HRD window width, $\omega = 1 - t_{\text{HRD}}$ |
| $\mathbf{w}_\sigma$ | COSMIC SBS96 spectrum of signature $\sigma$ |
| **Scenario-dependent (vary with $s$)** | |
| $\lambda_3^{(s)}(t)$ | SBS3 rate function for scenario $s$ on $[t_{\text{HRD}}, 1]$ |
| $w_i$ | Molecular time drawn for mutation $i$ in $[t_{\text{HRD}}, 1]$ |
| $r_i^{(s)}$ | Scenario scaling factor at $w_i$: scenario rate / baseline rate |
| $p_i^{\text{SBS3},(s)}$ | Imposed SBS3 probability for mutation $i$ under scenario $s$ |
| **Per-replicate draws** | |
| $S_i$ | Signature label assigned to mutation $i$ in a given replicate, e.g. $S_i \in \{\text{SBS1}, \text{SBS3}, \dots\}$ |
| $c_i$ | SBS96 channel of mutation $i$ in one replicate |

### 1 Simulation framework

#### 1.1 Motivation and interpretation of the simulation framework

This simulation framework was designed to assess the robustness of HRDTimer molecular timing and age estimates to departures from the constant SBS3 accumulation rate assumed by HRDTimer. Although this assumption is supported by the observed stability of SBS3 activity following HRD initiation, particularly during late-stage tumour evolution (Main text; Supplementary Fig. X), evaluating alternative SBS3 temporal-rate functions can provide a formal assessment of the sensitivity of timing estimates to potential changes in SBS3 mutation rates over time.

For each tumour, the original HRDTimer solution, including HRD and WGD timings in SBS1-based molecular time, was treated as the ground truth. These timings define a tumour-specific evolutionary timeline comprising successive temporal phases ($\phi$), as well as the intervals during which SBS3-associated mutations can accumulate. Alternative SBS3 rate functions were then simulated on this fixed timeline while keeping all other tumour-specific features unchanged, including copy-number state, assignment of mutations to temporal phases, and background mutational signature composition. This isolates the effect of temporal variation in SBS3 accumulation and enables direct comparison of timing estimates obtained under different SBS3 rate models.

#### 1.2 Simulation Framework – Diagram

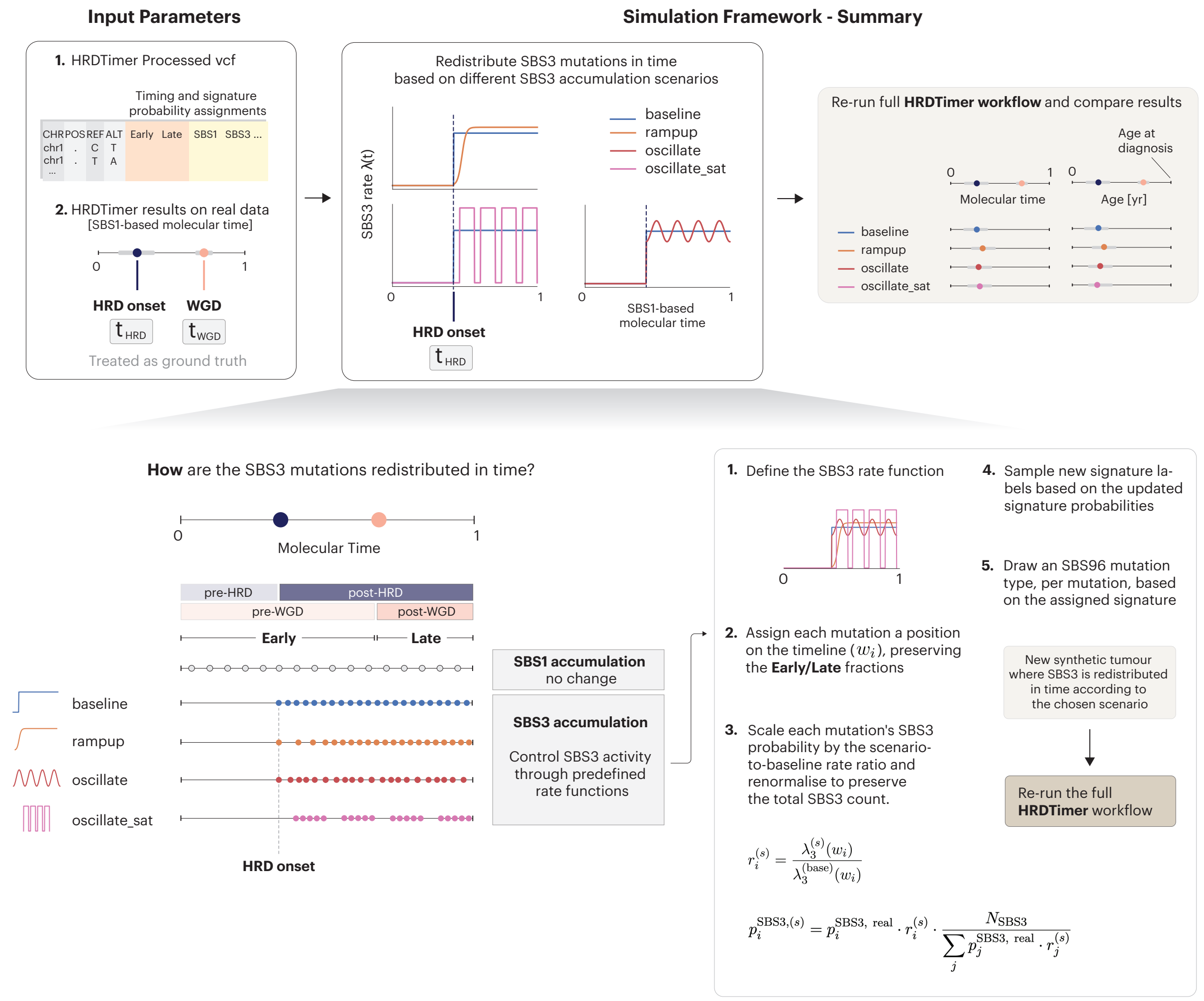


**Figure 1. Simulation framework for redistributing SBS3 mutations across molecular time.** HRDTimer-processed VCFs and real-data timing estimates for HRD onset ($t_{\mathrm{HRD}}$) and WGD ($t_{\mathrm{WGD}}$) are used as ground truth (top left). SBS3 mutations are then redistributed in molecular time under four rate functions $\lambda(t)$—baseline, rampup, oscillate and oscillate_sat—while the SBS1 profile is held fixed. For each scenario, mutations are assigned timeline positions preserving Early/Late fractions, their SBS3 probabilities rescaled by the scenario-to-baseline rate ratio and renormalised to conserve the total SBS3 count, and signature labels and SBS96 types resampled accordingly (bottom right). The full HRDTimer workflow is re-run on each synthetic tumour and the inferred timings compared across scenarios in molecular time and age (top right).

#### 1.3 Simulation Framework – Description

##### Input Parameters

The simulator is designed to preserve the biological characteristics of each tumour while enabling fair comparisons across alternative SBS3 temporal-rate scenarios and bootstrap replicates.

**Fixed inputs from the real tumour.** The following quantities are taken directly from the real tumour sample and are held fixed across all scenarios and replicates: the total mutation count $N$; per-mutation copy number $(\mathrm{MajCN}_i, \mathrm{MinCN}_i)$; HRDTimer phase classification $\phi(i) \in \{\mathrm{Early}, \mathrm{Late}, \mathrm{NA}\}$; soft pre/post-WGD probabilities $(p_i^G, p_i^S)$ for NA-phase mutations; the total SBS3 burden $N_{\mathrm{SBS3}}$; and the per-phase non-SBS3 signature mixture $\mathbf{m}_\phi$.

The latter two quantities are derived from the MuSiCal exposure vector obtained by fitting mutational signatures to the original tumour mutational catalogue. The probability that a mutation originated from SBS3, $p_i^{\mathrm{SBS3,\ real}}$, is used as the baseline in Step 3, whereas the remaining signature exposures define the phase-specific mixture vectors $\mathbf{m}_\phi$ used in Step 4.

**What changes between scenarios.** The temporal distribution of SBS3 activity is controlled by the rate function $\lambda_3^{(s)}(t)$. Different scenarios correspond to different choices of $\lambda_3^{(s)}(t)$, whereas all other quantities remain unchanged. For a given scenario, the rate function is fixed across all bootstrap replicates (Steps 1–3).

**What changes between replicates.** Within each (sample, scenario) pair, 200 bootstrap replicates are generated. In each replicate, mutation-level signature labels $S_i \in \{\text{SBS1}, \text{SBS3}, \dots\}$are resampled according to the scenario-specific SBS3 probabilities obtained after application of $\lambda_3^{(s)}(t)$ (Step 3). SBS96 trinucleotide channels $c_i$ are then resampled conditional on these signature assignments and the fixed per-phase $\phi(i) \in \{\text{Early}, \text{Late}\}$ signature mixtures (Steps 4–5). Consequently, the original trinucleotide contexts observed in the real tumour are not preserved in the simulated data.

##### Step 1 — Define the scenario rate function

A non-negative rate function $\lambda_3^{(s)}(t)$ is defined on $[t_{\text{HRD}}, 1]$ (SBS1-based molecular time) and rescaled so that

$$\int_{t_{\text{HRD}}}^{1} \lambda_3^{(s)}(t)\, dt = N_{\text{SBS3}}, \tag{1}$$

ensuring that all scenarios carry the same *expected* total SBS3 burden and differ only in *when*—before or after genome doubling—that burden is deposited. The actual SBS3 count in any individual replicate may vary slightly around $N_{\text{SBS3}}$ due to stochastic sampling in Step 4. The four scenarios and their rate functions are defined in Section 2.

##### Step 2 — Place mutations on the molecular timeline and compute scenario-specific SBS3 scaling factor

The only quantity that differs between scenarios is the SBS3 probability assigned to each mutation. All other tumour properties—phase classification $\phi(i)$ of mutations as Early/Late/NA, copy-number state, and non-SBS3 signature composition—are held fixed at their real-tumour values throughout.

To compute the per-mutation SBS3 rate adjustment, each mutation is assigned a representative time point $w_i$ within its phase window: $[t_{\text{HRD}}, t_{\text{WGD}}]$ for Early-phase mutations and $[t_{\text{WGD}}, 1]$ for Late-phase mutations, sampled according to the scenario rate function $\lambda_3^{(s)}(t)$. NA-phase mutations are first assigned to either Early or Late phase via a Bernoulli draw weighted by their pre- and post-WGD timing probabilities $(p_i^G, p_i^S)$.

The sole purpose of $w_i$ is to evaluate the scenario rate $\lambda_3^{(s)}(w_i)$ and derive the scenario-specific scaling factor $r_i^{(s)}$ used in Step 3. It has no effect on any signature other than SBS3.

##### Step 3 — Update each mutation's SBS3 probability

Each mutation is assigned a scaling factor equal to the ratio of the scenario rate to the baseline rate at its position $w_i$:

$$r_i^{(s)} = \frac{\lambda_3^{(s)}(w_i)}{\lambda_3^{(\text{base})}(w_i)}. \tag{2}$$

The original SBS3 probability from the real tumour, $p_i^{\text{real,SBS3}}$, is multiplied by $r_i^{(s)}$ and the resulting values are rescaled by a global constant so that the expected total SBS3 count is preserved:

$$p_i^{\text{SBS3},(s)} = p_i^{\text{SBS3, real}} \cdot r_i^{(s)} \cdot \frac{N_{\text{SBS3}}}{\sum_j p_j^{\text{SBS3, real}} \cdot r_j^{(s)}}. \tag{3}$$

Because the original probability $p_i^{\text{real,SBS3}}$ enters as a multiplicative factor, mutations with $p_i^{\text{real,SBS3}} \approx 0$ remain non-SBS3 regardless of the scenario.

##### Step 4 — Assign one signature label per mutation

For each mutation and bootstrap iteration, a single signature label $S_i$ is sampled:

$$S_i = \begin{cases} \text{SBS3}, & \text{with probability } p_i^{\text{SBS3},(s)}, \\ S_i' \sim \text{Categorical}(\mathbf{m}_{\phi(i)}), & \text{with probability } 1 - p_i^{\text{SBS3},(s)}, \end{cases} \tag{4}$$

where $\mathbf{m}_\phi$ is the MuSiCal exposure vector for phase $\phi$, taken from the real tumour, with the SBS3 component set to zero and the remaining entries renormalised. By construction, two properties hold: (i) each mutation has marginal probability $p_i^{\mathrm{SBS3},(s)}$ of being assigned to SBS3; (ii) within each replicate, the expected counts of *non-SBS3* signatures match the phase-specific mixture $\mathbf{m}_\phi$ observed in the real tumour. The SBS3 component of the phase mixture intentionally changes between scenarios — this redistribution between Early and Late phases is precisely what drives differences in $\hat{t}_{\mathrm{HRD}}$ across scenarios and is the quantity under investigation. Mutations with high $p_i^{\mathrm{real,SBS1}}$ — predominantly C>T transitions at CpG sites — carry $p_i^{\mathrm{real,SBS3}} \approx 0$ and are therefore unaffected by the scenario adjustment regardless of how SBS3 is redistributed. Only mutations with genuinely ambiguous SBS3 probability are reassigned between scenarios. This separation is what makes SBS1 a reliable molecular clock for HRD timing.

##### Step 5 — Generate SBS96 channels from assigned signatures

The mutation's original SBS96 trinucleotide channel from the real tumour is discarded and replaced by a fresh draw from the COSMIC spectrum $\mathbf{w}_{S_i} \in \Delta^{96}$ of the assigned signature:

$$c_i \sim \mathrm{Multinomial}(1, \mathbf{w}_{S_i}). \tag{5}$$

Regenerating channels from $\mathbf{w}_{S_i}$ produces a catalogue consistent with the imposed mixture. As a result, MuSiCal recovers the target exposures in expectation, while still reflecting realistic sampling variation across replicates.

##### Step 6 — Refit and re-estimate

The simulated catalogue is identical to the real tumour in every respect except the temporal distribution of SBS3 mutations, which is governed by $\lambda_3^{(s)}$. The standard HRDTimer pipeline is applied to this catalogue to obtain scenario-specific estimates $\hat{t}_{\mathrm{HRD}}^{(s)}$ and $\hat{t}_{\mathrm{WGD}}^{(s)}$, which are then translated to calendar age.

Repeating this procedure across 200 bootstrap replicates and all four scenarios yields a distribution of age-at-HRD estimates for each sample. These estimates are compared to the reference age obtained directly from the original tumour using HRDTimer (Section 3), allowing us to quantify how sensitive the inferred age is to the assumed post-HRD SBS3 rate.

#### 2 Scenario definitions

Four scenarios were implemented, spanning the range of biologically plausible post-HRD SBS3 behaviours (Table 1). The **baseline** assumes SBS3 accumulates at a constant rate once HRD is established, and serves as the reference model. The **ramp-up** scenario captures the idea that HRD damage may take time to fully manifest, with SBS3 accumulation accelerating gradually after the initial event. The two **oscillating** scenarios model episodic rather than continuous damage: the oscillate scenario uses a smooth sinusoidal modulation, while oscillate-sat imposes a square-wave pattern with complete quiescence between active phases.

All rate parameters are scaled by the post-HRD window $\omega = 1 - t_{\mathrm{HRD}}$, ensuring that scenarios are biologically comparable across samples with different HRD timing: a sample with late HRD (small $\omega$) has a proportionally shorter ramp and tighter oscillation period than one with early HRD. All rate functions are renormalised to carry the same total SBS3 burden, satisfying Eq. (1).

The period $T = 0.25\,\omega$ gives approximately four oscillation cycles across the post-HRD window for every sample—with the number of post-WGD oscillations after WGD depending on the timing of the latter. The amplitude $A = 0.2$ and square-wave parameters ($c_{\mathrm{high}} = 1.5$, duty cycle $d = 0.70$) were chosen to represent clearly marked but moderate departures from the constant-rate baseline, while keeping the rate non-negative everywhere.

Two design-level considerations are worth noting. First, the ramp-up scenario concentrates SBS3 mutations towards the post-HRD window by construction. Because HRDTimer estimates $\hat{t}_{\mathrm{HRD}}$ from the Late-phase SBS3-to-SBS1 ratio, this inflates that ratio and is expected to shift $\hat{t}_{\mathrm{HRD}}$ to later values. We explicitly quantify this induced bias to obtain an estimate of how much the inferred age changes under shifts from the observed tumour profiles, and we find that the resulting deviations remain within a reasonable range, indicating stable behaviour under this scenario (Figure 2D).

Second, for oscillating scenarios, instability can arise when the oscillation period $T$ becomes comparable to the length of the Late window $(1 - t_{\mathrm{WGD}})$. In this regime, the Late SBS3-to-SBS1 ratio captures only part of a cycle and may

**Table 1.** SBS3 rate function $\lambda_3^{(s)}(t)$ for each scenario, $t \in [t_{\mathrm{HRD}}, 1]$. The shape function $f(t)$ is multiplied by a normalisation constant $c = N_3 \big/ \int_{t_{\mathrm{HRD}}}^{1} f(t)\,dt$ to satisfy Eq. (1); $\omega = 1 - t_{\mathrm{HRD}}$. Amplitude, duty cycle, and timescale parameters are fixed design choices; only $c$ is determined from the data. Parameter values were chosen so that the resulting Early- and Late-phase SBS3 fractions remain consistent with the range observed across samples, ensuring each scenario represents a biologically plausible departure from the constant-rate baseline rather than an extreme redistribution.

| Scenario | Rate function $\lambda_3^{(s)}(t),\ t \in [t_{\mathrm{HRD}}, 1]$ | Biological interpretation |
|---|---|---|
| **Baseline** | $c$ | Constant SBS3 accumulation; reference model |
| **Ramp-up** | $c\exp\Big(\frac{t-t_{\mathrm{HRD}}}{\tau}\Big), \quad \tau = 0.10\,\omega$ | Gradual SBS3 activation |
| **Oscillate** | $c\Big(1+0.2\cos\frac{2\pi(t-t_{\mathrm{HRD}})}{T}\Big), \quad T = 0.25\,\omega$ | Sinusoidal ±20% modulation around the mean rate |
| **Oscillate-sat** | $c_{\mathrm{high}} \cdot \mathbf{1}\big[(t - t_{\mathrm{HRD}}) \bmod T < 0.70T\big],\ T = 0.25\,\omega,\ c_{\mathrm{high}} = 1.5\,c$ | Square-wave bursts (duty cycle 70%); rate ≈1.4× above mean during active phases, zero between |

not reflect the long-run average rate, which can lead to unstable or phase-dependent $\hat{t}_{\mathrm{HRD}}$ estimates in samples with short post-WGD windows.

In both cases, uncertainty is quantified and propagated through 200 bootstrap replicates, and we provide an empirical assessment of the impact of these design choices on the final age-at-HRD estimates.

#### 3 Results

The four SBS3 rate scenarios produce qualitatively distinct temporal profiles of SBS3 activity while preserving Early- and Late-phase SBS3 burdens within the range observed across the cohort. An example tumour is shown in Figure 2A,B. The oscillate scenario preserves a pre/post-WGD split close to the constant-rate baseline, whereas both the ramp-up and oscillate-sat scenarios shift burden toward the post-WGD phase (28% and 29% pre-WGD respectively, versus 35% under the baseline). For ramp-up this reflects the gradual accumulation of SBS3, shifting the overall burden towards the late period; for oscillate-sat it arises because the square-wave begins in the quiescent phase immediately after $t_{\mathrm{HRD}}$, effectively delaying the onset of SBS3 mutagenesis and reducing the burden accumulated before WGD. Both effects are captured directly in the phase split in Figure 2B, illustrating how different mechanistic assumptions translate into measurable differences in the Early/Late SBS3 distribution.

Across all 20 samples, age-at-HRD estimates under the baseline, oscillate, and oscillate-sat scenarios are tightly consistent with the reported HRD onset age (Figure 2C–D) and centred near zero bias. The ramp-up scenario introduces a systematic positive shift (+3.8 yr): by concentrating SBS3 in the late post-HRD window, it inflates the Late-phase SBS3-to-SBS1 ratio, causing HRDTimer to infer a later $\hat{t}_{\mathrm{HRD}}$ and consequently an older age at HRD onset. This bias is expected by design and quantifies the maximum effect of a systematically mis-specified rate assumption on the final age estimate.

Although point estimates remain largely robust across scenarios, confidence interval widths can vary between samples (Figure 2C). The widest intervals are concentrated in two groups of tumours. Samples with early HRD ($t_{\mathrm{HRD}} < 0.15$, $n = 6$) exhibit consistently broad confidence intervals across all scenarios (CI half-width 25–55,yr), reflecting overlap between the inferred HRD event and the period of normal-tissue SBS1 accumulation. A second group of samples with short post-WGD windows ($t_{\mathrm{WGD}} > 0.90$) shows greater sensitivity to the assumed SBS3 rate function. In these tumours, the very short Late phase results in less accurate estimates of the SBS3:SBS1 rate, reducing the precision of $\hat{t}_{\mathrm{HRD}}$. Under oscillating scenarios, this limitation can be further amplified when the oscillation period is comparable to the duration of the Late phase. Overall, these results indicate that uncertainty is driven primarily by the precision with which $\hat{t}_{\mathrm{HRD}}$ can be estimated, rather than by the specific SBS3 rate assumption, with the latter having only a modest effect except in tumours with very short post-WGD evolutionary windows.

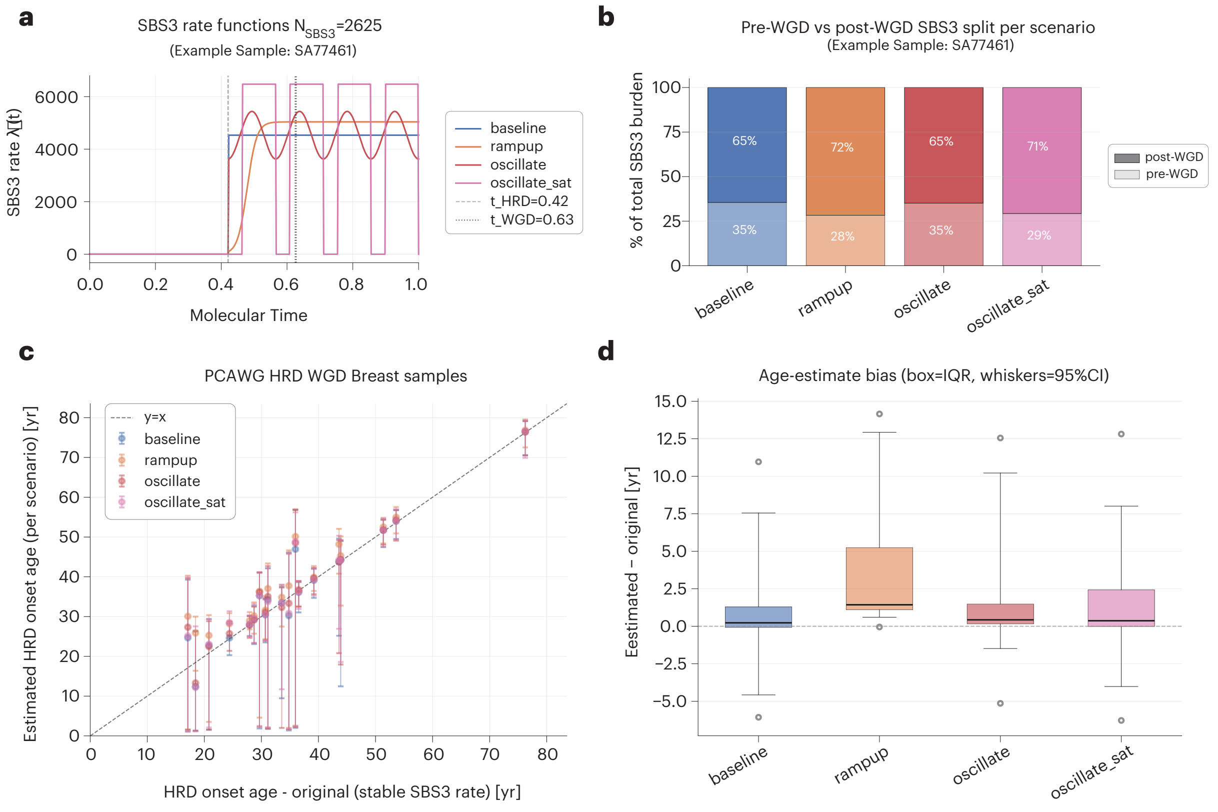


**Figure 2. HRD-onset age estimates are robust to post-HRD SBS3 rate assumptions across biologically motivated scenarios.** **a,** SBS3 rate functions $\lambda_3^{(s)}(t)$ for the four scenarios applied to an example sample (SA77461; $N_{\text{SBS3}} = 2{,}625$, $t_{\text{HRD}} = 0.42$, $t_{\text{WGD}} = 0.63$). All functions carry identical total SBS3 burden; vertical dashed lines mark $t_{\text{HRD}}$ and $t_{\text{WGD}}$. **b,** Fraction of total SBS3 burden falling in the pre-WGD and post-WGD phases for each scenario (same sample as **a**). The ramp-up and oscillate-sat scenarios shift burden toward the pre-WGD phase (28% and 29% post-WGD, respectively) compared to the constant-rate baseline (35%). **c,** Estimated median age-at-HRD versus HRD onset age derived under the original (constant-rate) HRDTimer model, across all $n = 20$ PCAWG breast cancer samples and four scenarios. Error bars, 95% bootstrap confidence intervals; dashed line, identity. **d,** Signed bias (estimated − original) per scenario. Box, IQR; whiskers, 95% bootstrap CI; dashed line, zero. Baseline, oscillate, and oscillate-sat estimates are centred near zero; ramp-up introduces a systematic positive shift of ≈2–4 yr by design.

Taken together, these results demonstrate that HRD-onset age estimates are robust to the assumed post-HRD SBS3 accumulation rate. Across three of the four scenarios tested — spanning constant, sinusoidal, and episodic SBS3 dynamics — estimates are statistically indistinguishable from those obtained under the constant-rate assumption that underlies the main HRDTimer model. Even under the ramp-up scenario, which represents a deliberate and systematic stress-test of the Late-phase SBS3 signal, the induced bias remains within 4 yr — smaller than the uncertainty inherent to the HRDTimer posterior for most samples. Samples that do show wide confidence intervals do so for reasons unrelated to the SBS3-rate assumption. In paricular, uncertainty is driven by limited molecular-time resolution resulting from early HRD timing or short post-WGD windows, both of which are intrinsic properties of the samples and are already apparent in the original HRDTimer analysis. The constant-rate assumption is therefore suitable for robust age-at-HRD estimation across the cohort.
